# Resonating with replicability: factors shaping assay yield and variability in microfluidics-integrated silicon photonic biosensors

**DOI:** 10.1101/2025.07.16.664198

**Authors:** Lauren S. Puumala, Samantha M. Grist, Karyn Newton, So Jung Kim, Stephen Kioussis, Sajida Chowdhury, Maggie Wang, Myra Wei, Yas Oloumi Yazdi, Avineet Randhawa, Yuting Hou, Lukas Chrostowski, Sudip Shekhar, Karen C. Cheung

## Abstract

The integration of biosensors and microfluidics has facilitated the development of compact analytical devices capable of performing automated and information-rich detection of myriad targets, both in the lab and at the point of need. However, optimization of microfluidics-integrated biosensor systems and replicability challenges present roadblocks in validation and commercialization. Understanding factors contributing to yield and replicability in biosensor performance is key to the development of biosensor optimization frameworks and technology translation beyond the research setting. Hence, for the first time, we present a detailed analysis of factors affecting performance, assay yield, and intra- and inter-assay replicability in microfluidics-integrated silicon photonic (SiP) evanescent-field microring resonator biosensors. Strategies for mitigating bubbles—a major operational hurdle and contributor to instability and variability in microfluidics-integrated biosensors—are analyzed to improve assay yield. Effective bubble mitigation is demonstrated by combining microfluidic device degassing, plasma treatment, and microchannel pre-wetting with surfactant solution. Both intrinsic and analyte-detection performance metrics and their replicability are quantified for the first time for sub-wavelength grating-based SiP biosensors, highlighting a path to further optimization. Lastly, the effects of sensor functionalization on analyte detection performance and replicability are evaluated. We compare polydopamine-vs. Protein A-mediated bioreceptor immobilization chemistries and spotting-vs. flow-mediated bioreceptor patterning approaches. We find that simple polydopamine-mediated, spotting-based functionalization improves spike protein (1 µg/mL) detection signal by 8.2× and 5.8× compared to polydopamine/flow and Protein A/flow approaches, respectively, and yields an inter-assay coefficient of variability below the standard 20% threshold for immunoassay validation. Overall, this work proposes a practical framework through which evanescent-field SiP biosensors, and microfluidics-integrated biosensors more generally, can be characterized, compared, and optimized to facilitate efficient biosensor development.

## 1. Introduction

Microfluidics-integrated biosensors have enabled the development of both information-rich lab-scale assays as well as portable, rapid, and mass-scale biomarker-based testing formats that obviate the need for highly-trained technicians, costly laboratory infrastructure, and long wait times (1–3). Microfluidic devices integrated with biosensors facilitate precise fluid handling and complete sample analysis on-chip, performing functions such as reagent mixing, separation, pre-concentration, routing, and continuous sampling, often in an automated manner (1,4). These miniaturized devices open up the opportunity to detect countless biomarker targets with high specificity, selectivity, sensitivity, and throughput, combined with low cost and ease-of-use (1,5–7). Nevertheless, challenges remain in biosensor validation and commercialization, especially with regard to optimizing devices to achieve replicable and robust sensing performance (5,8–11).

Microfluidics, in addition to the biosensor’s transducer and recognition element, contribute to the device’s performance and replicability. Critical performance metrics and their variability need to be quantified and considered throughout the development process—from early research to commercialization. However, the importance of replicability and the sources of variability from the transducer, recognition element, and microfluidics are often inadequately considered in research, hindering translation of novel technologies (12,13). Validation guidelines for biosensors and point-of-care analytical devices established by authorities such as the Clinical and Laboratory Standards Institute (CLSI), European Pharmacopeia (EP), United States Pharmacopeia (USP), and International Council for Harmonisation (ICH) include characteristics related to variability such as repeatability, reproducibility, robustness, and ruggedness (13,14). Nevertheless, underreporting and inconsistent reporting of metrics related to variability have been highlighted as major challenges in biosensors research that have slowed commercialization (10,15,16). In the fields of optical and electrochemical biosensors, some publications have explored improvements to variability-related metrics through optimizing transducer fabrication, surface functionalization, and signal processing/referencing/calibration approaches (14,17–19). Recently, Robbiani et al. (20) reviewed confounding factors that lead to variability in gas sensor performance for electronic nose applications and potential mitigation strategies. Developing a strong understanding of the sources of variability in each biosensor system can lead to the creation of adaptable frameworks for streamlined optimization of assay detection/quantification limits, yield, replicability, repeatability, and robustness at each stage of development toward clinical validation.

Evanescent-field silicon photonic (SiP) biosensors are one class of microfluidics-integratable optical biosensors that can facilitate information-rich sensing in a compact and low-cost format. They are produced using wafer-scale semiconductor fabrication techniques that can enable reproducible and inexpensive production at large scales (21). SiP sensors consist of nanoscale silicon or silicon nitride structures that guide and manipulate light due to their high refractive index (RI) contrast with the substrate (e.g., silicon dioxide) and surrounding media (21,22). Various SiP biosensing architectures have been demonstrated, including microring resonators (MRRs), which consist of a waveguide that is looped back on itself in a ring and a straight bus waveguide that couples light into the ring; such architectures have been described in detail elsewhere (21–24). Briefly, a change in the local RI (Δn_surface_, e.g., due to analyte binding) within the MRR’s evanescent field changes the effective RI (Δn_eff_) of the waveguide, which is transduced to a quantifiable shift in the resonance peak (Δλ_res_) of the MRR’s optical transmission spectrum (22,25). A millimeter-scale SiP sensor chip can be patterned with tens of MRRs whose transmission spectra can be simultaneously monitored in real time for multiplexed sensing (26,27). By functionalizing their surfaces with analyte-specific bioreceptors, SiP sensors can be used to selectively detect myriad biomarker targets including proteins (28–30), nucleic acids (31–33), toxins (34,35), viruses (36,37), and bacteria (38), with limits of detection down to the pg/mL scale and dynamic ranges spanning 2–3 orders of magnitude (30,39,40). These biosensors are typically integrated with microfluidics to control the sequential delivery of small volumes of functionalization reagents, analyte samples, wash buffers, and signal amplification reagents to the sensor surface (41).

As for all biosensor systems, low variability in sensing performance—both intra-assay (i.e., across multiple sensing structures on a chip probed during a single assay) and inter-assay (i.e., across replicate assays performed on different chips)—is essential for validation of diagnostic SiP biosensors. Fig. 1 outlines example system factors that contribute to variability in SiP sensing assays and Table 1 summarizes how each of these factors is expected to vary across sensors, microfluidic channels, and replicate chips/assays. Factors related to microfluidics integration, such as reagent depletion due to biomolecule adsorption in the fluidic system, flow rate instability, and alignment of sensing structures within microfluidic channels, may contribute to variability by affecting the concentration of sample delivered to the surface, c_surface_. Gas bubble formation in microfluidic channels is a frequent operational hurdle in microfluidic assays, especially those using polydimethylsiloxane (PDMS)-based devices (42). Bubbles can damage sensor surface functionalization chemistry (42–44) and interfere with the sensing signal, presenting a major, and often unpredictable, source of variability. Factors related to surface functionalization, such as the orientation, density, and stability of immobilized bioreceptors, reagent variability, and nonspecific/off-target binding, contribute to a biosensor’s variability by affecting the amount of material bound at the surface and, in the case of SiP sensors, Δn_surface_ (15). These factors depend on the choice of bioreceptor, immobilization chemistry, and patterning technique (27). For a given bioreceptor, covalent rather than noncovalent immobilization may enhance the stability of the functionalized surface (27). Immobilization chemistries employing reaction protocols that are robust to day-to-day variations can improve yields and reduce inter-assay variability (45,46). In addition, the method by which the bioreceptor solution is patterned and incubated on the sensor surface (e.g., in-flow or spotting) dictates the mass transport phenomena governing bioreceptor delivery to the surface, influencing the uniformity and density of immobilized bioreceptors, thus affecting detection performance and variability (27). Finally, factors related to the transducer, including fabrication variations in waveguide geometry and oxide open processes as well as surface cleanliness, can contribute to variability by affecting how Δn_surface_ is transduced to Δλ_res_. For example, SiP MRRs based on more complicated waveguide geometries, such as sub-wavelength gratings (SWGs), offer improved sensitivity to conventional strip waveguide devices but may be more susceptible to variability in fabrication and waveguide wetting due to their smaller feature sizes (47). Variability in wetting from trial to trial would cause performance variability, and potentially bubbles during experiments. Potential strategies to reduce the impact of each of these factors are included in Table 1; these strategies span the system design (e.g., sensor design, use of reference sensors and the functionalization chosen for those sensors (48), or microfluidics materials), functionalization strategy (e.g., type of bioreceptor), assay design and protocol (e.g., sample dilution), and data analysis (e.g., resonance peak curve-fitting and averaging of replicate sensors).

**Fig. 1.**
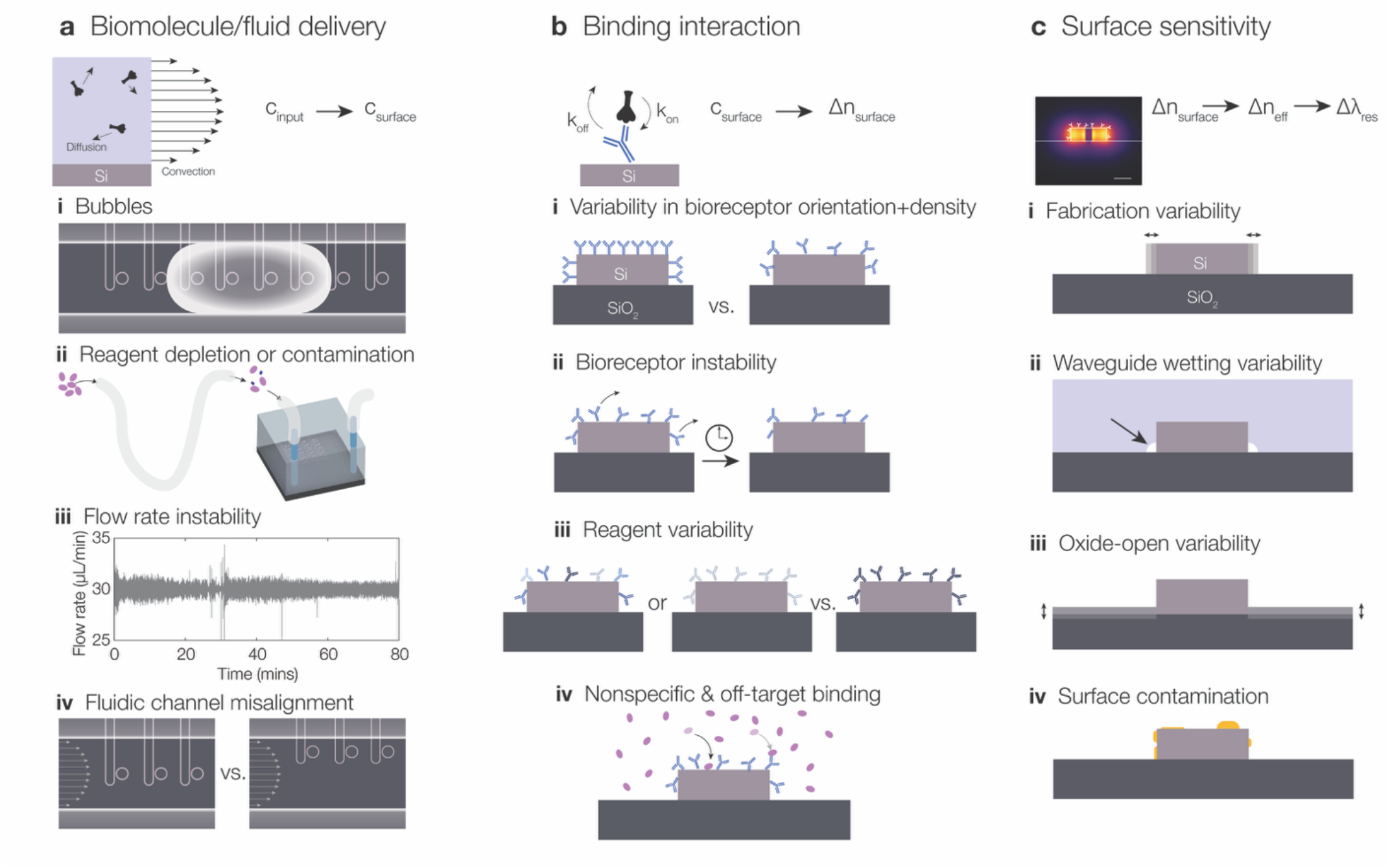
Example system factors contributing to analyte detection replicability using SiP biosensors. The sensor’s signal (resonance peak shift Δλ_res_) is related to the analyte concentration in the input sample (c_input_), but the signal depends on (a) the biomolecule delivery to the surface via convection and diffusion (which controls the concentration of the analyte in the fluid at the surface of the sensor c_surface_ as a function of c_input_), (b) the binding interaction between the analyte and the functionalized resonator surface (which controls how the RI at the surface of the sensor, Δn_surface_, changes as a function of the bulk analyte concentration in the fluid at the sensor surface), and (c) the sensor’s surface sensitivity (which controls how the effective RI, Δn_eff_, and the resonance peak position change in response to the changes in RI at the surface of the sensor). There are factors within each of these stages that can impact analyte detection replicability; 4 example factors are illustrated for each stage. **(a)** (i) Bubbles can disrupt delivery of the analyte to the sensor surface, cause significant sensor instability, and potentially damage sensor functionalization. (ii) Reagents can be depleted and the fluid can be contaminated with other materials within the microfluidics or pumping system used to deliver the analyte to the sensor surface. (iii) Fluid pumping systems can have instability in the delivered flow rate changing the rate of convective transport. (iv) Depending on where the sensors are located in the microfluidic channels, they can be exposed to different fluid velocities, again changing rates of convective transport. **(b)** (i) Depending on the type of bioreceptor used and how it is immobilized on the surface, there can be wide variation in the orientation and density of the bioreceptors, which regulate how much analyte can bind to the surface. (ii) Bioreceptors can be unstable on the surface of the sensor, causing bioreceptor removal or damage over time between functionalization and detection. (iii) There can be variation in the performance of different lots of bioreceptors used for functionalization, and polyclonal antibodies have inherent variability and a range of affinities. (iv) Surface functionalization is susceptible to both nonspecific binding (e.g., to the bare surface of the resonator if inadequately blocked, depicted by the black arrow) and off-target binding between the bioreceptors and non-analyte components of the sample matrix (depicted by the grey arrow). **(c)** (i) Fabrication variability in the waveguide geometry affects the resonators’ intrinsic sensing performance. (ii) Depending on the geometry, sensor surface hydrophilicity, and how fluid is delivered to the surface, there can be variations in waveguide wetting, resulting in different proportions of the waveguide cross-sections being exposed to fluid. (iii) If a silicon dioxide cladding is used during SiP sensor fabrication (standard for many foundries), it needs to be removed over the sensors to permit fluid access to the sensor surface. This oxide-open etch typically has variability in etch depth, resulting in different heights of exposed waveguide and thus different sensitivities. (iv) The surface cleanliness of the sensor can change the sensitivity (if the binding interaction is moved further from the sensor surface) as well as how much analyte can bind.

**Table 1.**
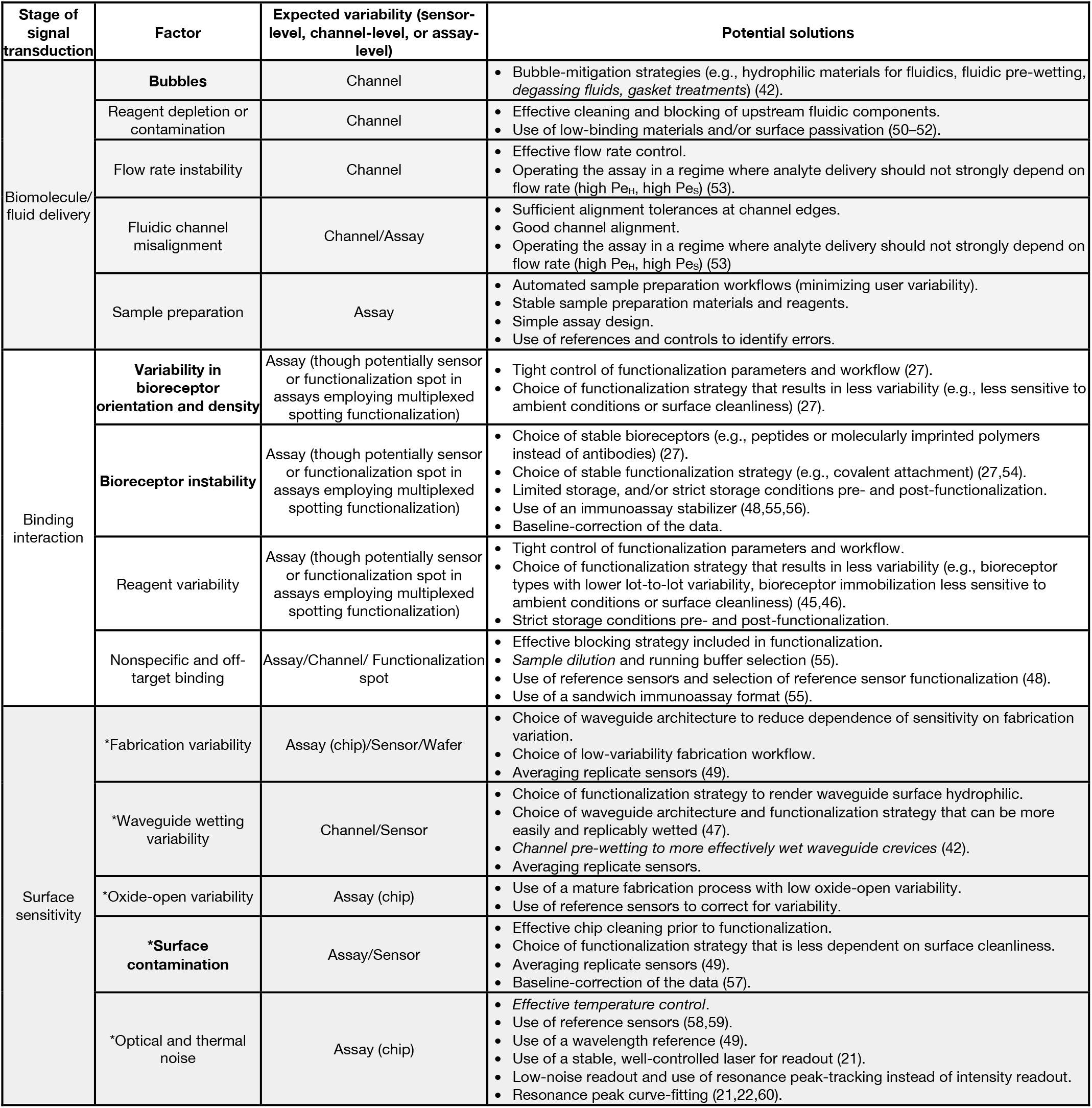
Factors affecting intrinsic and analyte-detection replicability, in each of the 3 stages of detection/signal transduction described in Fig. 1. We note for each factor whether we may expect it to present variability on the sensor level, the microfluidic channel level, or the assay level, as well as the potential solutions. *Italics* denote solutions that may need to be implemented/performed at the point-of-use rather than during design/manufacturing (more challenging to implement for decentralized testing applications). Factors preceded by an asterisk (^*^) are specific to SiP biosensors, while others are applicable to microfluidics-integrated biosensors more generally. **Bolding** highlights factors that we aimed to modulate and assess through our experimental work in this manuscript, focusing on bubble-mitigation strategies and surface functionalization approaches.

| Stage of signal transduction | Factor | Expected variability (sensor-level, channel-level, or assay-level) | Potential solutions |
| --- | --- | --- | --- |
| Biomolecule/fluid delivery | **Bubbles** | Channel | Bubble-mitigation strategies (e.g., hydrophilic materials for fluidics, fluidic pre-wetting, *degassing fluids, gasket treatments*) (42). |
|  | Reagent depletion or contamination | Channel | Effective cleaning and blocking of upstream fluidic components.<br>Use of low-binding materials and/or surface passivation (50–52). |
|  | Flow rate instability | Channel | Effective flow rate control.<br>Operating the assay in a regime where analyte delivery should not strongly depend on flow rate (high $Pe_H$, high $Pe_S$) (53). |
|  | Fluidic channel misalignment | Channel/Assay | Sufficient alignment tolerances at channel edges.<br>Good channel alignment.<br>Operating the assay in a regime where analyte delivery should not strongly depend on flow rate (high $Pe_H$, high $Pe_S$) (53) |
|  | Sample preparation | Assay | Automated sample preparation workflows (minimizing user variability).<br>Stable sample preparation materials and reagents.<br>Simple assay design.<br>Use of references and controls to identify errors. |
| Binding interaction | **Variability in bioreceptor orientation and density** | Assay (though potentially sensor or functionalization spot in assays employing multiplexed spotting functionalization) | Tight control of functionalization parameters and workflow (27).<br>Choice of functionalization strategy that results in less variability (e.g., less sensitive to ambient conditions or surface cleanliness) (27). |
|  | **Bioreceptor instability** | Assay (though potentially sensor or functionalization spot in assays employing multiplexed spotting functionalization) | Choice of stable bioreceptors (e.g., peptides or molecularly imprinted polymers instead of antibodies) (27).<br>Choice of stable functionalization strategy (e.g., covalent attachment) (27,54).<br>Limited storage, and/or strict storage conditions pre- and post-functionalization.<br>Use of an immunoassay stabilizer (48,55,56).<br>Baseline-correction of the data. |
|  | Reagent variability | Assay (though potentially sensor or functionalization spot in assays employing multiplexed spotting functionalization) | Tight control of functionalization parameters and workflow.<br>Choice of functionalization strategy that results in less variability (e.g., bioreceptor types with lower lot-to-lot variability, bioreceptor immobilization less sensitive to ambient conditions or surface cleanliness) (45,46).<br>Strict storage conditions pre- and post-functionalization. |
|  | Nonspecific and off-target binding | Assay/Channel/ Functionalization spot | Effective blocking strategy included in functionalization.<br>*Sample dilution* and running buffer selection (55).<br>Use of reference sensors and selection of reference sensor functionalization (48).<br>Use of a sandwich immunoassay format (55). |
| Surface sensitivity | *Fabrication variability | Assay (chip)/Sensor/Wafer | Choice of waveguide architecture to reduce dependence of sensitivity on fabrication variation.<br>Choice of low-variability fabrication workflow.<br>Averaging replicate sensors (49). |
|  | *Waveguide wetting variability | Channel/Sensor | Choice of functionalization strategy to render waveguide surface hydrophilic.<br>Choice of waveguide architecture and functionalization strategy that can be more easily and replicably wetted (47).<br>*Channel pre-wetting to more effectively wet waveguide crevices* (42).<br>Averaging replicate sensors. |
|  | *Oxide-open variability | Assay (chip) | Use of a mature fabrication process with low oxide-open variability.<br>Use of reference sensors to correct for variability. |
|  | **\*Surface contamination** | Assay/Sensor | Effective chip cleaning prior to functionalization.<br>Choice of functionalization strategy that is less dependent on surface cleanliness.<br>Averaging replicate sensors (49).<br>Baseline-correction of the data (57). |
|  | *Optical and thermal noise | Assay (chip) | *Effective temperature control.*<br>Use of reference sensors (58,59).<br>Use of a wavelength reference (49).<br>Use of a stable, well-controlled laser for readout (21).<br>Low-noise readout and use of resonance peak-tracking instead of intensity readout.<br>Resonance peak curve-fitting (21,22,60). |

In the field of SiP biosensors, several papers have reported intra-and/or inter-assay variability in intrinsic (i.e., sensing of changes in the bulk RI of the fluid cladding the sensor surface) and/or analyte sensing performance, quantified in terms of the coefficient of variability (CV), as summarized in ESI Table S1. For example, Mudumba et al. (49) thoroughly characterized variability in SiP MRR biosensor performance in the commercial Genalyte system, reporting CVs for intrinsic and analyte sensing performance within (ring-to-ring and between groupings of rings) and between assays (chip-to-chip and across replicate experiments). However, this type of thorough analysis is not typically reported for the multitude of SiP sensors reported at the research stage, with the replicability of some newer sensor architectures like SWGs not yet being established. Additionally, to the best of our knowledge, no prior works have combined this type of thorough analysis of SiP biosensor intra- and inter-assay replicability and performance with a detailed exploration of how these metrics are affected by microfluidics integration-, functionalization-, and transducer-related factors.

In this paper, we present for the first time a detailed analysis of factors impacting SiP biosensor replicability and assay yield, based on literature references, our team’s experience, and new experimental data. We present a general framework for studying the factors that contribute to biosensor replicability, and outline a specific manifestation of this framework for SiP biosensors including characterization and comparison of several conditions. Our experimental results in Section 3, supported by detailed methods in Section 2 and the Electronic Supplementary Information as well as comparison to existing literature, clarify critical factors limiting replicability and performance in our system. For the same sets of silicon SWG MRRs, we present a comparison of intrinsic sensor performance and inter- and intra-assay replicability (bulk RI sensitivity, stability, and system limit of detection) with performance and replicability for analyte detection in a SARS-CoV-2 spike protein demonstration assay. We also discuss factors contributing to assay yield, considering a failed assay as one in which the data cannot be used or trusted, e.g., due to major sensor signal instability due to air exposure, equipment failure, reagent contamination/deterioration, and operator errors. In this work, we focus on factors related to microfluidics integration that contribute to yield, such as bubbles in microfluidic devices, and their mitigation. Finally, we show how the choice of sensor functionalization strategy impacts sensor signal and replicability in demonstration assays comparing noncovalent Protein A-mediated and covalent polydopamine (PDA)-mediated bioreceptor immobilization chemistries as well as in-flow and spotting-based bioreceptor patterning techniques. To the best of our knowledge, we present the first demonstration of PDA-mediated bioreceptor immobilization on a silicon MRR architecture and show its feasibility as a simple and stable functionalization approach for these sensors. Together, these contributions will highlight a practical interdisciplinary strategy for developing and optimizing SiP biosensor systems with broad applicability to microfluidics-integrated biosensors more generally.

### 2. Experimental

In order to understand and improve variability in our system, we applied the following framework (generally applicable to all biosensors) to our SiP system:

1. Deconstruct the system into different stages of detection and identify factors in the system at each stage that impact performance and replicability, and how they may be expected to manifest at the inter- and intra-assay levels, as well as any other levels relevant to the system of interest (e.g., inter-channel for our system).
2. Identify a set of performance metrics that isolate different contributing factors and assess which factors contribute to variability in each performance metric.
3. Characterize the performance metrics and their replicability at the inter- and intra-assay levels to identify factors that are limiting performance and replicability in the system.

Table 1 presents a specific example of applying the first stage of the framework to SiP biosensors. Although specifically applied to our system (SiP biosensors integrated with pressure-driven continuous-flow microfluidics), portions of Table 1 are also applicable to other classes of biosensors (e.g., the factors in the Biomolecule/fluid delivery stage are applicable to other classes of surface sensors integrated with pressure-driven continuous-flow microfluidics).

Table 2 presents the manifestation of the second stage of the framework for SiP biosensors. Each performance metric defined in Table 2 has a set of factors at the biomolecule delivery, analyte-binding, and surface sensitivity stages of detection that can impact its replicability. Table 2 also presents a summary of some of these factors for each metric. Using SiP ring resonator evanescent-field biosensors integrated with pressure-driven continuous-flow microfluidics as an example system, we first worked to improve assay yield by reducing bubble formation within the microfluidic channels and subsequently characterized this range of common performance metrics and their replicability. In this section, we report a summary of the methods that we used for this work, with detailed methods supporting inter-laboratory reproducibility presented in the Electronic Supplementary Information (ESI) Section S2. In addition, product, vendor, and lot information for chemicals and reagents used in this work are provided in ESI S2.1. Experimental data describing the characterization of our system (implementing stage 3 of the framework) are presented in Section 3, with the experimental performance metric summary included in Table S19.

**Table 2.**
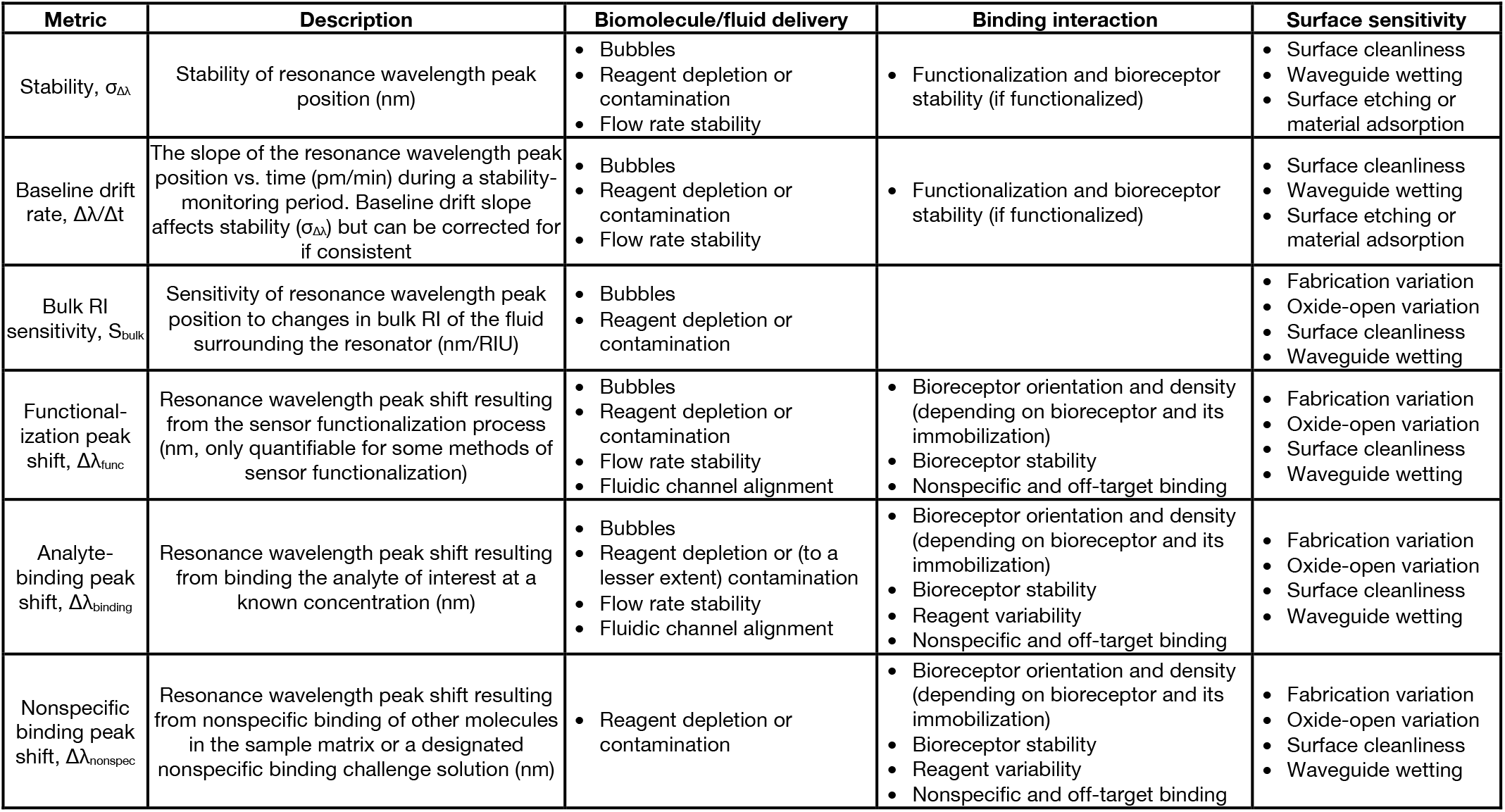
Common performance metrics quantified for SIP biosensors, and their dependence on factors affecting analyte-detection replicability in each of the 3 stages of detection/signal transduction described in Fig. 1.

### 2.1. Sensor chip and microfluidics design, fabrication, and assembly

#### 2.1.1. Photonic sensor chip

Photonic SWG MRR circuits were designed with the following design parameters: ring radius R = 30 µm, coupling gaps g_c_ = [500, 550] nm, grating period Λ = 250 nm, duty cycles δ = [0.65, 0.7], waveguide width w = 500 nm, and waveguide thickness t = 220 nm, as previously described by our group (47). All photonic experiments reported in this paper used rings with an SWG duty cycle of 0.7 and a coupling gap of 500 nm. The layout included input and output grating couplers designed to couple 1550 nm light between the air-clad chip and benchtop tunable laser and detectors. The photonic chips (Fig. 2(a)) were fabricated on silicon-on-insulator (SOI) wafers through the Applied Nanotools Inc. (ANT, Edmonton, AB, Canada) NanoSOI Fabrication Service Silicon Device Layer process using 100 keV electron beam lithography and reactive ion etching (61). Further details regarding the photonic design and fabrication are provided in ESI S2.2. The SWG MRRs used in this work were found to exhibit the following performance metrics when clad with ultrapure water: extinction ratio ER = 22.3 ± 3.7 dB, quality factor Q = 8.39 × 10^3^ ± 2.13 × 10^3^, and free spectral range FSR = 4.24 ± 0.08 nm (all metrics are reported as the mean ± standard deviation calculated across 8 replicate chips with 8 replicate MRRs analyzed per chip). Further details regarding the photonic design, fabrication, and resonator performance metric quantification are provided in ESI S2.2

**Fig. 2.**
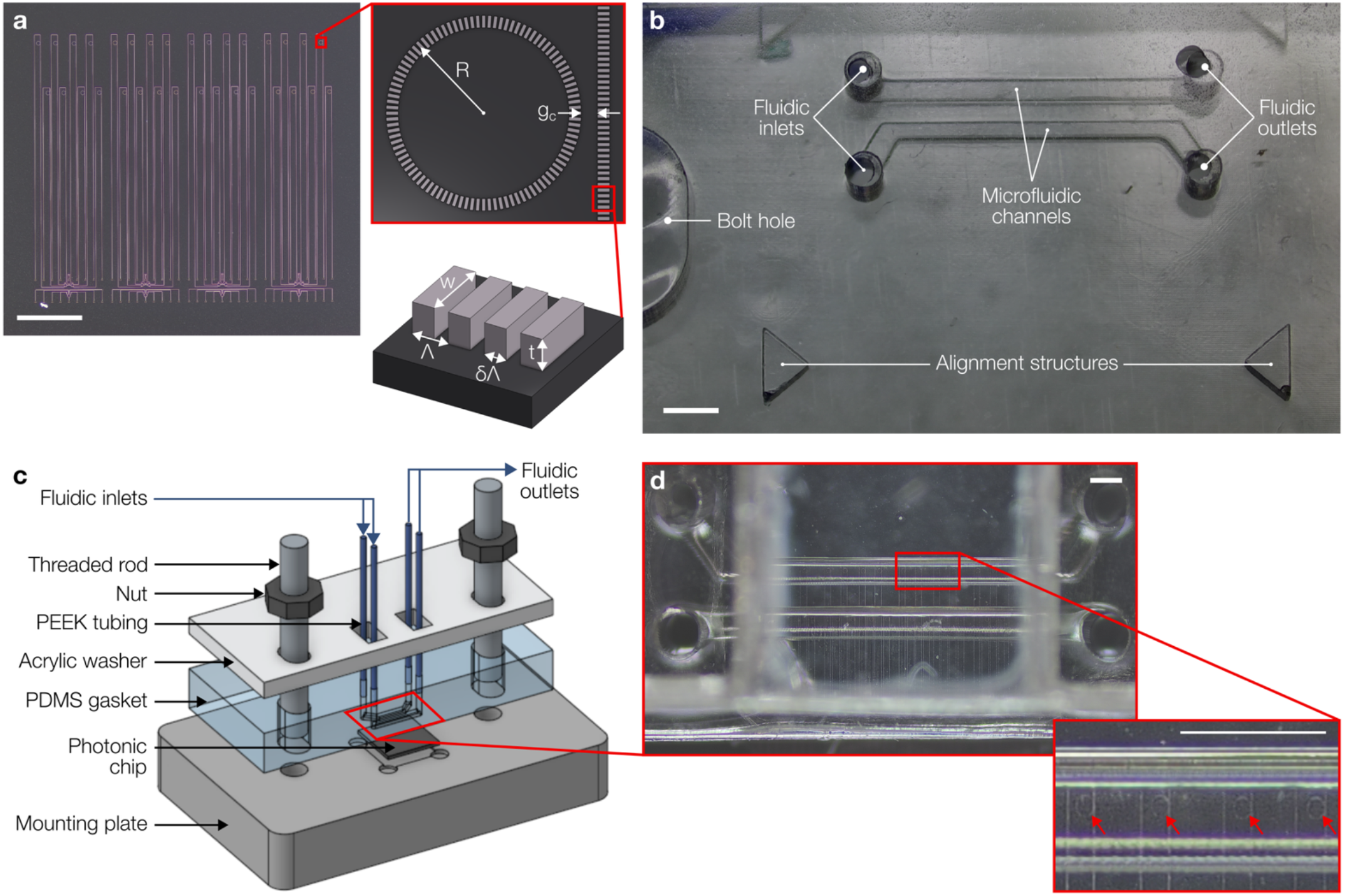
Photonic chip and microfluidic gasket design and integration. **(a)** Optical micrograph of SiP MRR chip (scale bar represents 1000 µm) with insets showing annotated illustrations of a SWG MRR and SWG waveguide (not to scale). **(b)** Optical micrograph of PDMS gasket without assembly (scale bar represents 1000 µm). **(c)** Illustration of photonic chip-microfluidic gasket assembly. **(d)** Optical micrograph of the photonic MRRs aligned within the microfluidic channels of the microfluidic gasket with inset showing a zoomed-in view of four MRRs aligned within the top microfluidic channel (scale bars represent 500 µm). Red arrows point to MRRs in (d). Rings are circular as depicted in (a) but appear oval-shaped in (d) due to optical aberrations from imaging through the gasket assembly.

#### 2.1.2. Microfluidics

Microfluidic gaskets (Fig. 2(b–d)) to deliver liquid solutions to SiP chips for sensor performance characterization were fabricated using Sylgard™ 184 poly(dimethylsiloxane) (PDMS) (Ellsworth Adhesives, Hamilton, ON, Canada) molded against 3D printed molds through soft lithography, as previously described by our team (47). The microfluidic PDMS gasket designs include two parallel microfluidic channels, 300–400 µm in width and 500 μm in height over the region of the photonic chip containing the sensors, expanding into 600 μm diameter circular input/output regions, as illustrated in Fig. 2(b). Depending on the experiment, PDMS gaskets were assembled with the photonic chip without further treatment or degassed overnight in a vacuum desiccator then treated with air plasma immediately prior to assembly with the photonic chip for bubble mitigation (refer to Section 2.4). To assemble the setup for fluidic testing (Fig. 2(c)), the photonic chip was placed in the machined recess of an aluminium mounting plate and the PDMS gasket was aligned to the chip under visual inspection (Fig. 2(d)). An acrylic washer, threaded rods, and nuts were used to seal the fluidics against the photonic chip. Additional details are provided in ESI S2.3.

### 2.2. Photonic-fluidic testing setup

#### 2.2.1. Photonic testing setup

Sensing experiments were performed using a custom optical testing setup (Maple Leaf Photonics, Seattle, WA, USA), which has been described in detail elsewhere (47). Briefly, the photonic chip-gasket assembly was mounted on a motorized and temperature-controlled stage. A 12-channel lidless fiber array (VGA-12-127-8-A-14.4-5.0-1.03-P-1550-8/125-3A-1-1-0.5-GL-NoLid-Horizontal, OZ Optics, Ottawa, ON, Canada) was aligned to the chip’s grating couplers to interface with a C-band swept tunable laser (Agilent 81682A, Agilent Technologies, Santa Clara, CA, USA) and optical detectors (Agilent 8164A and Keysight N7744C, Keysight Technologies, Santa Rosa, CA, USA). This setup enabled simultaneous readout from up to eight photonic MRR sensors. Alignment of the fiber array to the photonic chip was performed using open-source PyOptomip software (Python 2.7, 32-bit) (62), which controlled the position of the optical testing setup’s linear motorized stages and communicated with the tunable laser and detectors. During sensing experiments, data were acquired from the photonic chip using a custom Python software GUI, which swept the C-band tunable laser source over a user-defined wavelength range (∼15 nm wide) and recorded and saved the output transmission spectra from the photonic chip every 20–30 s. Additional details and a schematic of all connections between the photonic chip and external equipment are provided in ESI S2.4.

#### 2.2.2. Fluidic testing setup

A two-channel modular Fluigent LineUp™ series fluid control system (Fluigent, Le Kremlin-Bicêtre, France), controlled by OxyGEN software (SSFT-OXY), was used to control and monitor the delivery of reagents to both channels of the microfluidic gasket mounted to the photonic chip. This system is illustrated in Fig. S2 and described in detail in ESI S2.5. Briefly, each fluidic channel included 10 reagent reservoirs, a pressure-based flow controller, 10-position bidirectional valve, and flow rate sensor. Bubble traps (PG-BT-REC25UL, PreciGenome, San Jose, CA, USA) were connected to the outlets of each flow rate sensor. In experiments using aqueous solutions for sensor pre-wetting (refer to Section 2.6), the ends of PEEK capillary tubing (IDEX 1531B, Cole-Parmer Canada, Quebec, QC, Canada) connected to the bubble trap outlets were directly inserted into the inlet ports of the microfluidic gasket. In experiments using ethanol-based pre-wetting, an additional pre-wetting assembly was included in the path between the bubble trap outlets and microfluidic gasket inlet ports for each microfluidic channel to facilitate bubble-free switching between ethanol, manually delivered using a syringe, and the Fluigent system. This assembly was required due to the hydrophobic PTFE bubble trap membranes becoming wetted by organic solvent solutions, making them no longer able to remove bubbles from the flowing fluid. A ∼2 cm piece of PEEK tubing friction fit to Tygon® 0.02” ID, 0.06” OD microbore tubing (Masterflex® Microbore Transfer Tubing, Tygon® ND-100-80, Cole-Parmer Canada, Quebec, QC, Canada) was inserted into each of the outlet ports of the microfluidic gasket to direct effluent into a waste container. In ESI 2.6. we have outlined strategies that we have employed for mitigating common sources of error with these types of automated pressure-driven flow control systems.

### 2.3. Fluidic priming and pre-wetting

Before connecting the fluidic delivery system to the microfluidic gasket, all lines of the Fluigent system that were to be used for each experiment were primed with appropriate solutions to avoid pushing air from empty lines through the microfluidic gasket after pre-wetting (which could dry out channel crevices and introduce bubble nucleation sites). The complete map of solutions with which each reservoir was primed for each assay type is presented in Table S12.

We then proceeded to the pre-wetting process. For the experiments that used ethanol pre-wetting (bubble monitoring experiments testing ethanol pre-wetting and assays that detected 20 µg/mL spike protein using the Protein A/flow-mediated functionalization process), a pre-wetting assembly (details provided in ESI S2.5) was employed rather than the Fluigent system. The pre-wetting assembly was used to manually deliver 1–2 mL of anhydrous ethanol to each channel of the dry microfluidic gasket to fully wet all crevices, followed by flushing the channels with ∼2 mL of ultrapure water. Next, the pre-wetting assembly was disconnected and the primed Fluigent system was connected to the gasket without introducing air bubbles and set to flow ultrapure water at 30 µL/min prior to proceeding with sensing experiments. For the experiments that used Triton X-100 pre-wetting (bubble monitoring experiments testing Triton X-100 pre-wetting and all assays that detected 1 µg/mL spike protein), the Fluigent system was primed with 0.3 mM Triton X-100 in PBS after priming all other lines. The Triton X-100-primed lines were then connected to the inlet ports of the microfluidic gasket and the pre-wetting solution was delivered at a constant flow rate of 1– 2 µL/min to fully wet the channels. After ∼5 minutes of pre-wetting solution flow, the Fluigent reservoir being delivered was switched to the assay’s running buffer and the flow rate was increased to 30 µL/min. Detailed descriptions of fluidic priming and pre-wetting protocols are provided in ESI S2.6.

### 2.4. Characterizing bubbles in microfluidic devices

Static contact angles were measured (ESI S2.7) for all three pre-wetting liquids (ultrapure water, ethanol, and 0.3 mM Triton X-100 in PBS) on the surface of a molded PDMS gasket before and after treating the PDMS piece with air plasma for 75 seconds at 400 mTorr and high power in a Harrick Plasma cleaner (PDC-001-HP, Harrick Plasma, Ithaca, NY). Post-plasma contact angles were measured immediately after plasma treatment and at 30-minute intervals over a 300-minute period following plasma treatment during which the gasket was stored under atmospheric conditions.

*In situ* bubble-monitoring experiments were performed to evaluate the efficacy of bubble mitigation strategies for PDMS microfluidics, including channel pre-wetting with the three different pre-wetting liquids (ultrapure water, ethanol, and 0.3 mM Triton X-100 in PBS), PDMS degassing, and PDMS air plasma treatment. Table S3 lists the combinations of bubble mitigation strategies that were tested and the number of trials for each. These experiments were performed using photonic chip-gasket assemblies, as described in Section 2.2. Where applicable, PDMS gaskets were degassed in a vacuum desiccator overnight (connected to our house vacuum line, which ranges from 13–23 HgV) until use, and/or treated with air plasma (as described above). Gaskets were then promptly assembled with the rest of the chip-gasket assembly within 30–40 minutes. Channels were pre-wet with the specified liquids prior to introducing working liquids.

Bubble nucleation and entry within the microfluidic channels was monitored via time-lapse imaging over several hours of aqueous working liquid flow (protocol details provided in ESI S2.8). From reviewing the compiled monitoring videos for each trial, the following metrics were recorded: number of bubble nucleation sites, number of bubbles entering and exiting the microfluidic channel, duration of time during which each bubble was present in the microfluidic channel, and maximum bubble size. To normalize bubble dynamics across experiments, the number of bubbles entering and nucleating in the channel per hour, as well as the proportion of the monitored time where bubbles were visible in the channel, were also calculated.

Percent reduction values for each metric of bubble frequency and duration/impact (y) were calculated for each experimental group relative to the control condition using the formula:

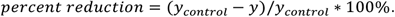

### 2.5. Polydopamine deposition, capture antibody spotting, and blocking

PDA coating of SiP sensors for assays using PDA-based antibody immobilization was performed via a one-pot aqueous reaction found to yield a PDA film thickness of 1.8 ± 0.1 nm (ESI S2.9). The bare sensor chip was first placed in a glass crystallization dish containing 40 mL of freshly prepared dopamine hydrochloride solution (2 mg/mL in Tris buffer). PDA coating of the chip was allowed to proceed for 30 minutes at room temperature under stirring at ∼100 rpm. After 30 minutes, the chip was transferred to a beaker containing fresh Tris buffer, thoroughly rinsed with ultrapure water, then dried with nitrogen gas. The chip was stored under ambient conditions until use.

To prepare sensor chips for assays using spotting-based antibody immobilization, PDA coated chips were manually spotted with 20 µL of 500 µg/mL spike protein capture antibody in PBS, containing 10% (v/v) glycerol and 0.005% (v/v) Triton X-100 using a micropipette (63). Glycerol and Triton X-100 were included to slow droplet evaporation and minimize the coffee ring effect (55,64). Spotted chips were incubated at room temperature for one hour in a closed petri dish lined with cleanroom wipe dampened with ultrapure water to create a humid environment. After incubation, each chip was thoroughly rinsed with PBS. Sensor chips were then blocked with 20 mg/mL BSA in PBS for one hour at room temperature, followed by thorough rinsing in PBS (65,66). Chips were dried with nitrogen, then immediately assembled with a microfluidic gasket.

### 2.6. Characterizing sensor performance

#### 2.6.1. Intrinsic sensor performance

Bulk RI sensitivity (S_bulk_) and stability experiments were performed on 4 replicate SiP chips. After pre-wetting the microfluidic channels with ethanol, ultrapure water was delivered to the photonic chip-gasket assembly at 30 µL/min while the optical fibre array was coupled to the chip. An automated fluidic protocol defined in Fluigent’s OxyGEN software was then used to supply five standard solutions of [0, 62.5, 125, 250, and 375] mM sodium chloride (NaCl) in ultrapure water (see ESI Table S4 for measured RIs) to the sensor in ascending and then descending order of concentration for 20 minutes per step at 30 µL/min while photonic sensing data were acquired. Resonance peak shift data from both the ascending and descending concentrations were used to compute S_bulk_ for each sensor. A 10-minute duration of the ultrapure water (0 mM NaCl) flow at the beginning of the experiment was used for computing baseline drift and stability metrics. Additional details and data are provided in ESI S2.10.

#### 2.6.2. Analyte detection performance

Spike protein detection assays were performed using the following functionalization approaches to compare both noncovalent (Protein A) and covalent (PDA) antibody immobilization chemistries and in-flow and spotting-based antibody patterning: Protein A/flow, PDA/spotting, and PDA/flow. Detection of 20 µg/mL spike protein in buffer was demonstrated for Protein A/flow functionalization only. Detection of 1 µg/mL spike protein in buffer was demonstrated for all three functionalization approaches. To perform each assay, the appropriate sensor chip was interfaced with a microfluidic gasket (Section 2.1). Protein A/flow assays used bare SiP sensor chips and surface functionalization with Protein A, BSA blocking, and capture antibody immobilization were performed in-assay. PDA/spotting assays used sensor chips already functionalized via PDA coating, antibody spotting, and BSA blocking (Section 2.5) prior to fluidic integration. PDA/flow assays used PDA-coated sensor chips; antibody immobilization and BSA blocking were performed in-assay. Chip-gasket assemblies were mounted on the photonic-fluidic testing setup (Section 2.2) and the fluidic delivery system was loaded with all required functionalization and assay reagents and primed (Section 2.3) prior to connection with the chip-gasket assembly. Gasket treatment conditions and pre-wetting fluids used in these assays are summarized in ESI S2.11. Fluidic protocols prepared in Fluigent’s OxyGEN software were used to automate reagent delivery while sensing data were acquired. Both fluidic channels were subject to identical fluidic protocols, allowing us to quantify intra-channel replicability (all sensors in one microfluidic channel), intra-assay replicability (all sensors across both channels in an assay) and inter-assay replicability (across trials).

Where applicable, assays began by delivering functionalization reagents (Protein A/flow assays: Protein A (100 µg/mL), BSA (1 mg/mL), and capture antibody (20 µg/mL); PDA/flow assays: capture antibody (20 µg/mL) and BSA (20 mg/mL)) to the sensor, interspersed with PBS buffer flow. For all assays using PDA-based antibody immobilization, the functionalized sensors were rinsed with a pH 2.2 glycine-HCl buffer prior to the detection assay to remove excess loosely-bound capture antibodies (67,68); this step was not included in Protein A/flow assays, where it would be expected to remove adsorbed Protein A/antibody complexes from the surface. All detection assays were then performed by subjecting the functionalized sensors to an initial running buffer stabilization period, followed by exposure to 1 mg/mL BSA in PBS to challenge the surface blocking, a 10-minute running buffer rinse, spike protein detection (1 or 20 µg/mL in PBS), and a final running buffer stabilization period. Fluidic protocols for all assay formats, including fluidic reservoir configurations, reagent compositions, flow rates, and flow times are provided in ESI S2.11.

Initial and final peak shift drift rates and resonance peak shifts due to BSA and spike protein binding were quantified for each assay, in addition to the intra- and inter-assay CVs for each metric. Where relevant, Protein A and capture antibody binding shifts were quantified in the same manner as BSA and spike protein binding shifts. Quantification was performed on baseline-corrected sensor data, with baseline fitting being performed on the initial drift rate quantification regions. For the Protein A/flow assays, the initial draft rate region was defined as the first running buffer stabilization period before Protein A deposition. For all other assays, it was defined as the running buffer stabilization period after the glycine-HCl buffer rinse. To mitigate the effects of any bulk RI-induced signal while different solutions were being delivered to the sensors, all resonance peak shifts were quantified by comparing the resonance peak positions during the washes before and after each assay stage while the fluid surrounding the sensor was running buffer. The final drift rate quantification region for all assays was defined as the running buffer stabilization period immediately after spike protein delivery. Bulk RI sensitivities were also quantified for each sensor, with details and data provided in ESI S2.10.

### 2.7. Sensing data analysis

An overview of the SiP MRR sensing data acquisition and analysis workflow is visualized in Fig. S6, with details reported in ESI S2.12. Briefly, acquired optical spectra were analyzed using a custom Python script to Lorentzian-fit each resonance peak to extract its central wavelength and perform peak-tracking over time (across sequential sweeps in each experiment). Lorentzian fitting was used to more accurately discern the central wavelength and reduce the impact of noise on the measured optical power, in comparison with peak identification on the raw data. Our peak-tracking algorithm uses information from all resonance peaks in the sweep range rather than a single peak, to further reduce the impact of noise on the measurement of a single peak. After generating tracked resonance peak shift vs. time datasets, these data were postprocessed to extract the metrics of interest (Table 2). All binding assay data were baseline-corrected to mitigate the effects of any sensor drift and functionalization dissociation on the subsequent quantification of peak shifts and baseline drift slopes for each assay stage.

## 3. Results and discussion

### 3.1. Improving SiP assay yield by reducing bubble nucleation in microfluidic devices

In an effort to improve the yield and performance of SiP biosensor assays, we first sought to identify and empirically characterize the efficacy of strategies to reduce bubble nucleation in pressure-driven continuous flow microfluidics integrated with SiP biosensors. The impact of bubbles on SiP assays is twofold: (1) bubbles disrupt the ability of the sensor to read out meaningful data and (2) exposure to air can damage proteins on functionalized sensor surfaces. Bubbles can have a catastrophic impact on the performance and replicability of optical biosensors that read out RI, since the large RI contrast between air and aqueous solutions causes drastic instability in the sensor signal as bubbles come into contact with the sensors. Examples of such sensor instability during exposure to bubbles are provided in ESI Section S4. Moreover, drying of proteins (e.g., bioreceptors) on the sensor surface due to exposure to air can destabilize their three-dimensional structures, changing or diminishing functionality (69). Bubbles often repeatedly form in pressure-driven continuous-flow microfluidic systems after bubble nuclei are generated when the fluidic channels are first wet with liquid (Fig. 3(a)) (42). High advancing contact angles between the liquid and channel walls cause air to be readily trapped in surface crevices, creating Harvey nuclei that can lead to cyclic bubble expansion and release throughout assays as liquids containing super-saturation levels of gases (due to the pressure that is used to drive flow) travel past the bubble nuclei (42).

Though it is a popular material in microfluidics-integrated biosensors, particularly in academia (70,71) but also in some commercial devices (e.g., Fluidigm, Emulate (71,72)), PDMS suffers from some well-known limitations, including susceptibility to protein adsorption and gas bubble formation due to high surface hydrophobicity as well as absorption of small hydrophobic molecules due to high permeability (21,41,73,74). Alternative materials have been used for the fabrication of pressure-driven continuous flow microfluidics integrated with SiP biosensors, including mylar and teflon (39,68,75), SU-8 (76,77), and off-stoichiometry thiol-ene (OSTE) polymer (78). Digital microfluidics have also been demonstrated as an alternative to pressure-driven continuous flow microfluidics on SiP sensors (79–81).

Despite the limitations discussed above, PDMS was selected for microfluidics fabrication in this work owing to its low cost, ease of rapid prototyping, and optical transparency, which enables visually-guided alignment of microchannels to on-chip optical sensing structures (21,73). While susceptible to bubbles, PDMS’s gas solubility and permeability also create opportunities for using the PDMS itself as a power-free vacuum source to assist with bubble mitigation and reduce bubble nuclei (82,83). Degassing the PDMS piece under vacuum for an extended period of time reduces the concentration of air dissolved in the material (82). When the degassed device is returned to atmospheric pressure, the “stored vacuum” (the low concentration of dissolved air) in the PDMS can drive diffusion of air down the concentration gradient into the PDMS, potentially helping to eliminate bubbles and Harvey nuclei formed during channel wetting if wetting is conducted quickly after device removal from vacuum (82). Vacuum-aided bubble traps can also be integrated into PDMS devices, leveraging the high gas permeability of the material (84,85). Other bubble mitigation strategies target the reduction of advancing contact angles during the critical initial channel wetting process to minimize the formation of Harvey nuclei (42). To this end, PDMS microfluidics can be treated with plasma to temporarily increase surface hydrophilicity and channels can be pre-wetted with solvent or surfactant solutions prior to introducing working fluids.

In this work, we explored the efficacy of channel pre-wetting with solvent and surfactant solutions, PDMS degassing, and air plasma treatment as bubble mitigation strategies for biosensor-integrated PDMS microfluidics. We evaluated the effect of these strategies on bubble introduction, nucleation, and growth in the channels of PDMS microfluidic gaskets mounted to silicon chips. Bubbles were monitored over several hours of fluid flow via image capture. From these captures, we quantified bubble duration (the proportion of monitored time when bubbles were present) and frequency (number of bubbles introduced or nucleated per hour) in the microchannels as metrics to assess the efficacy of the tested bubble mitigation strategies. Bubble duration is a critical metric, quantifying the proportion of a microfluidic experiment during which the biosensor results might be assumed to be unreliable due to the presence of bubbles. Bubble nucleation frequency quantifies the presence and impact of Harvey nuclei within the microfluidic channel, while bubble introduction frequency quantifies bubbles that were nucleated or introduced elsewhere in the microfluidic system (e.g., in connectors or components upstream of the microfluidic channels, or potentially at the inlet of the microfluidic channel). Because experiments where bubbles are visible in the channels for a large proportion of the experiment time are likely to fail, we further define an assay yield metric as the proportion of trials in which bubbles were visible in the channels for <10% of the monitored time. Together, these metrics describe how many bubbles were present in experiments under each condition as well as the impact that they would have on the biosensor system or assay.

Three pre-wetting liquids were evaluated: ultrapure water, ethanol, and 0.3 mM Triton X-100 in PBS. Ultrapure water typically demonstrates a hydrophobic contact angle (>90º) with PDMS and was used here as a control. As illustrated in Fig. 3(b), ethanol and 0.3 mM Triton X-100 exhibit lower static contact angles on untreated PDMS (ethanol: 28.1º ± 3.2º, 0.3 mM Triton X-100: 44.7º ± 4.9º) compared to ultrapure water (94.4º ± 4.8º), and their use as pre-wetting liquids was, therefore, hypothesized to mitigate bubbles by reducing the formation of Harvey nuclei during gasket wetting. As shown in Fig. 3(b), air plasma treatment transiently reduces the static contact angles of all three pre-wetting liquids on PDMS (ultrapure water: < 5º, ethanol: < 5º, 0.3 mM Triton X-100: 16.0º ± 1.9º, measured immediately after plasma treatment), with contact angles increasing over time after treatment due to the migration of low molecular weight chains from the bulk to the surface of the PDMS (86,87). Hence, plasma treatment shortly before channel wetting was tested to evaluate its effect on reducing the occurrence of bubbles in the microfluidic devices. Degassing the PDMS gasket under vacuum overnight prior to mounting to the silicon chip and introducing liquids was also tested. Where controllable with our automated fluid control system (ultrapure water and Triton X-100 pre-wetting conditions), pre-wetting was performed at a low flow rate of 1 µL/min (resulting in average linear flow velocities of ∼5–6.7 mm/min—depending on channel width—within our microfluidic channels) to further reduce advancing contact angles during liquid introduction. The delivery method for ethanol was more complex due to inline bubble trap compatibility issues with organic solvents. Ethanol pre-wetting required manual injection into the PDMS channels using a syringe connected with segments of Tygon® and PEEK tubing, rather than being regulated by the automated flow control system. In addition, ethanol is not suitable for pre-wetting when denaturation-prone biological elements are present in the device (42); hence, this approach cannot be used to pre-wet microfluidic devices interfaced with biosensor chips that have already been functionalized with antibodies or other denaturation-prone bioreceptors.

Fig. 3(c–e) summarizes the quantified bubble duration and frequency metrics we used to compare the effects of plasma treatment, degassing, and pre-wetting liquid on bubble formation, and Table 3 presents the average ± standard deviation among the trials for each metric and condition. Consistent with the reduction in contact angle conferred by air plasma treatment of the gasket, we found a reduction in bubble incidence and impact when the gasket was plasma-treated prior to Triton X-100 introduction, as described in Fig. 3(c). While these results suggest that plasma treatment confers a 68% reduction in the proportion of an experiment that could be impacted by the presence of bubbles in the channel, it alone is not sufficient for robust, reliable biosensing as it leaves 10 ± 24% of the experiment with potentially unreliable measurements, and an assay yield (percentage of trials with bubbles visible for <10% of the monitored time) of only 83.3%. We expect that both plasma treatment and PDMS degassing would primarily affect bubble nucleation rates and not bubble entry rates, since they would not be expected to impact bubble formation in locations upstream of the gasket in the fluidic system. However, there may be bubble nucleation sites where the tubing interfaces with the gasket I/Os that cannot be easily resolved in the monitoring images and result in bubbles being quantified as “entering” rather than “nucleating”.

**Table 3.**
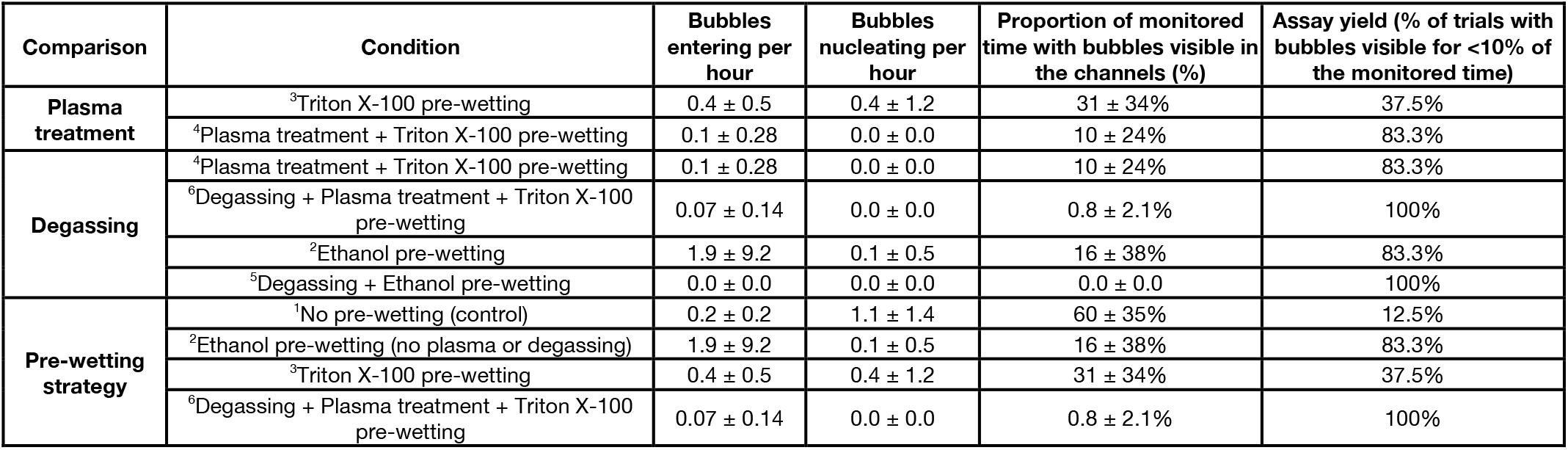
Quantification of the bubble duration and frequency metrics reported in Fig. 3(c–e), organized by the conditions compared in each subfigure. Superscript numbers at the beginning of the “Condition” column report which of the 6 tested conditions described in Table S3 the row represents. Duplicate rows are present as some of the same datasets were used for different comparisons.

As seen in Fig. 3(d) and Table 3, when the PDMS was degassed for >12 hours and removed from the desiccator immediately prior to use, we observed improvements in bubble duration and incidence for both Triton X-100 pre-wetting after plasma treatment as well as ethanol pre-wetting. These results show that gasket degassing confers a further 92% reduction in the proportion of an experiment that could be impacted by the presence of bubbles for gaskets treated with plasma and pre-wetted with Triton X-100, and a 100% reduction for gaskets pre-wetted with ethanol. Assay yields increased from 83.3% to 100% for both the Triton + plasma and ethanol conditions when the PDMS gasket was degassed prior to use. While we are not aware of previous work demonstrating degassed PDMS as a passive pumping source to reduce Harvey nucleus formation and bubble incidence, a study by Zhou et al. (88) demonstrated that nanoliter volumes of liquid could be dispensed into arrays of microwells using degassed PDMS microchannels that acted as an internal vacuum pumping source. The authors showed that, even after exposure to atmospheric pressure for 30 minutes after degassing and prior to connecting to the filling fluid, the degassed PDMS could drive fluid flow, filling the microwells within 11 minutes. This previous work is consistent with our results, demonstrating PDMS’s ability to effectively store vacuum and enable air removal from fluidic channels within the PDMS device after moderate periods of exposure to atmospheric pressure. In this work, we have shown how PDMS degassing can also be an effective bubble mitigation strategy in PDMS microfluidics by helping to remove Harvey nuclei even after they form.

From the results presented in Fig. 3(c-d), we hypothesized that a gold-standard pre-wetting strategy combining Triton X-100 pre-wetting with plasma treatment and PDMS degassing might perform similarly to ethanol-based pre-wetting, while also bringing advantages like compatibility with sensor functionalization, tubing, and system materials and components that are incompatible with organic solvents. We tested this condition against Triton X-100 pre-wetting without gasket treatment as well as against ethanol pre-wetting and a control. The results presented in Fig. 3(e) show high rates of bubble entry into the channels and considerable bubble nucleation in the control condition using ultrapure water for pre-wetting without plasma treatment or degassing of the PDMS gasket. The high nucleation rate indicates that the strategy was ineffective at sufficiently wetting all crevices within the PDMS structure, likely due to the hydrophobic contact angle with the PDMS surface.

Despite ethanol having the lowest contact angle on PDMS, ethanol pre-wetting without plasma treatment or degassing resulted in intermediate levels of bubble occurrence in the channels, potentially due to the more complex and less controllable nature of the manual ethanol pre-wetting protocol. This process may be susceptible to bubble introduction during the manual fluid injection process and when connecting the microfluidic gasket to the Fluigent system for the delivery of working fluids, as well as the additional potential for bubble nucleation in crevices and dead volume regions in the tubing assembly and connectors added inline to permit ethanol pre-wetting. Less control over the flow rate during the manual pre-wetting process may have also resulted in higher advancing contact angles. Pre-wetting with 0.3 mM Triton X-100 was facilitated by the Fluigent system in combination with bubble traps, allowing for controlled flow delivery while effectively removing bubbles from the liquid before entering and pre-wetting the PDMS microfluidic channels, but bubble nucleation and entry were still observed. When Triton X-100 pre-wetting was combined with plasma treatment and degassing of the gasket (both individually found to reduce bubble incidence and impact as discussed above), we found a large reduction in bubbles entering (82.5% reduction) and nucleating (100% reduction). In summary, compared to the control condition, ethanol pre-wetting, Triton X-100 pre-wetting alone, and pre-wetting with Triton X-100 after gasket plasma treatment and degassing conferred 73%, 48%, and 99% reduction in the proportion of the experiment time that could be unreliable due to the presence of bubbles. As a result of these improvements, the assay yield increased from 12.5% in the control condition up to 83.3%, 37.5%, and 100%, respectively.

Large variability in standard deviations and CVs was present in all bubble-monitoring metrics. We hypothesize that this large variability results from the probability of bubble nucleation sites forming in the monitored channel region or upstream of it: not all trials will generate bubble nuclei, but those that do will often have a large effect on our monitored metrics (large number and/or duration of bubbles). We thus expect high variation across trials when computing the average and standard deviation of these metrics across the 3–24 trials for each condition (Table S3), including both bubble-free and bubble-containing trials. Indeed, in Table 3 we observe that conditions that yielded high values in our metrics also showed high standard deviation values due to the probability of trials in which bubble nuclei were not formed. This high variability complicates the statistical analysis and significance testing. Although we do not observe a significant impact of degassing due to this large variation, the only conditions in which we measured 100% assay yield (bubbles visible <10% of the monitored time) used degassing, suggesting that this strategy is effective.

The reduced bubble nucleation observed here with both ethanol and surfactant solutions is consistent with work reported by Pereiro et al. (42), which showed that wetting a PDMS pillared microfluidic chamber with ethanol or surfactant solution (10% sodium dodecyl sulfate in water) allowed for complete filling of the chamber with liquid, while directly wetting the chamber with water led to trapped air. Pereiro et al. reported that although the degree of wetting was greater for ethanol than surfactant solution on PDMS, both of these solutions yielded initially bubble-free chambers via microscopy (which may miss sub-resolution Harvey nuclei). Building upon this previous work, here we have additionally quantified the reductions in the incidence and impact of bubbles in PDMS microfluidic channels during >2 hours of working fluid flow offered by alcohol- and surfactant-based pre-wetting, combined with plasma treatment and degassing. We have also employed a nonionic surfactant solution that is much gentler than 10% SDS and is compatible with antibody-functionalized sensors.

Overall, our experimental findings highlight that pre-wetting microfluidic channels with Triton X-100 solution, combined with degassing and plasma treatment of the PDMS microfluidic gasket shortly before use, are effective strategies for bubble mitigation in PDMS microfluidics-integrated SiP biosensors. Together, these strategies yielded a 99% reduction in the proportion of the experiment time that could be unreliable due to the presence of bubbles. The degassing and plasma treatment likely work to reduce the presence of Harvey nuclei in the microfluidic gasket itself, while pre-wetting with low-concentration Triton X-100 may better wet other crevices of the fluidic system upstream of the gasket and at the inlet interface as compared to a surfactant-free solution. Although solvent (ethanol) pre-wetting is also an effective strategy, the complexity of the required fluidic setup as well as the incompatibility with biofunctionalized sensors led us to choose the Triton X-100 pre-wetting fluid with plasma and degassing gasket pretreatment for our assays involving pre-functionalized resonators. A statistical analysis of the data presented in Fig. 3 is provided in ESI S3.

### 3.2. Replicability of SiP intrinsic performance and analyte detection

Seeking to understand how the choice of specific performance metric affects the measured performance and replicability of SiP biosensor-based measurements, we characterized and compared several performance metrics on a representative set of SiP biosensors. Using SiP integrated circuits containing a set of 8 SWG ring resonator biosensors operating near 1550 nm (47), we first integrated the SiP chip with a PDMS microfluidic gasket (89), such that 4 sensors were aligned in each of two separate microfluidic channels. We then delivered automated microfluidic protocols to characterize their intrinsic performance and subsequently to characterize their analyte-detection performance in a simple demonstration assay format using the spike protein of the SARS-CoV-2 virus in buffer as a representative biomarker. More complex samples (e.g., real clinical specimens) introduce additional contributions to variability and require additional controls (e.g., negative control samples, control or reference sensors lacking bioreceptors). The simple assay format used here was selected in order to increase protocol complexity in a stepwise manner and to facilitate the analysis of contributions to assay variability related to the transducer, microfluidics integration, and surface functionalization.

We first sought to understand the intrinsic performance of our SiP biosensors, and characterized the bulk RI sensitivity, stability, system limit of detection, and baseline drift rate, as well as the intra- and inter-assay replicability of these performance metrics. Briefly, we exposed the sensors to RI standard solutions (0–375 mM NaCl) through the microfluidic channels after ethanol pre-wetting and analyzed the resonance peak positions during the water baseline (yielding measurements of stability and drift rate) and during exposure to each standard solution (yielding measurements of S_bulk_). The results of this intrinsic performance characterization are presented in Fig. 4. Fig. 4(a) shows the quantified resonance peak shift vs. time as the sensors were exposed to each RI standard solution, and Fig. 4(b) shows the resonance peak shift vs. bulk RI change of each standard solution, linearly fitted to extract the S_bulk_ for each sensor in each trial. The 4 sensors in each microfluidic channel (plotted as different line traces) show near-identical signals in Fig. 4(a). The quantified S_bulk_, baseline drift rate (the slope of a linear fit to the peak position vs. time data), stability σ_Δλ_ (standard deviation of the peak position across a 20-min exposure to water), and system limit of detection (sLoD = 3σ_Δλ_/S_bulk_) (21,90) are reported as beeswarm plots with each point representing the data from one sensor in Fig. 4(c-f), respectively. The measured RIs of the standard solutions and sensor-by-sensor data are reported in ESI S2.10.1.

**Fig. 4.**
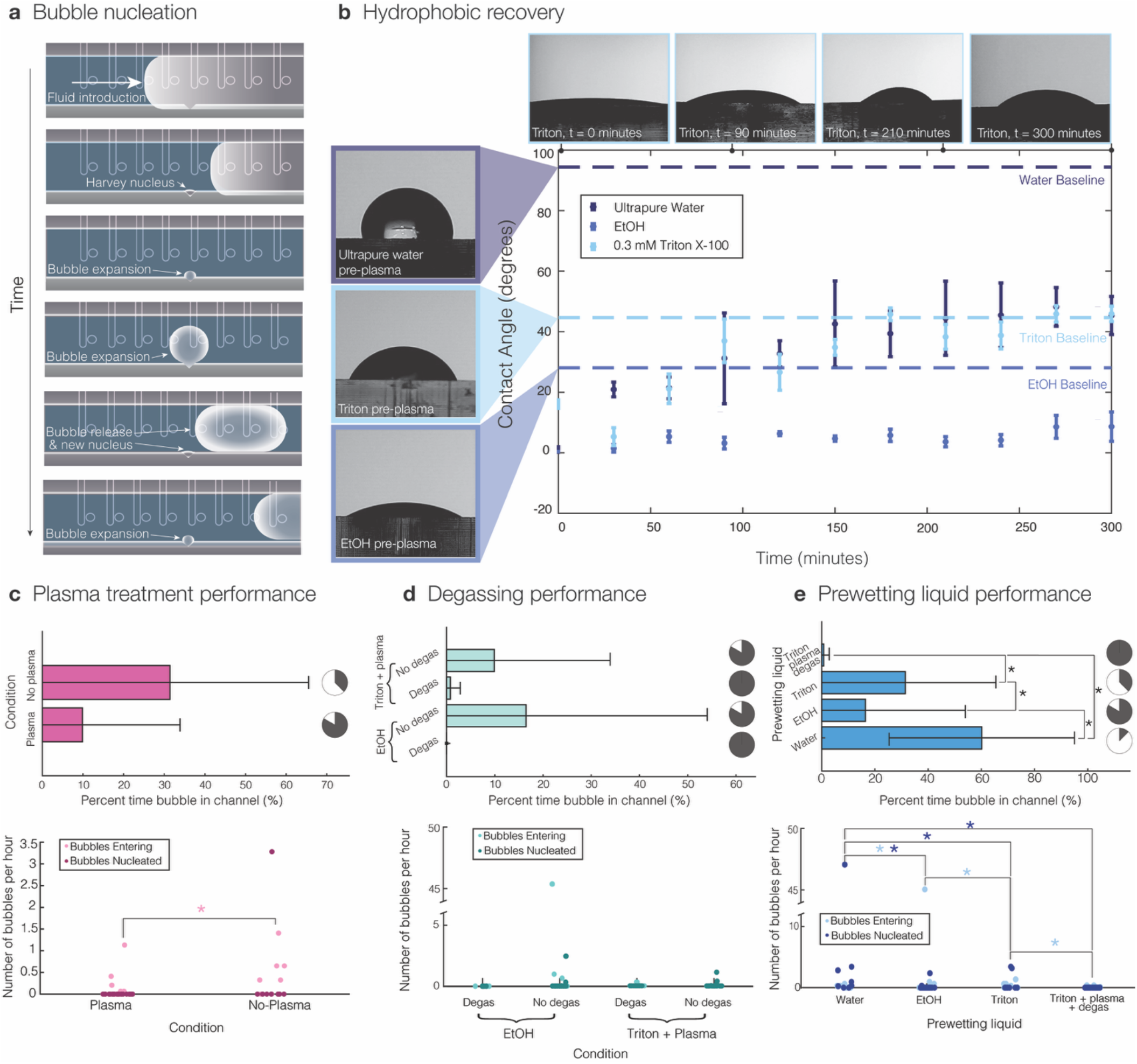
Mitigating bubble formation in microfluidic systems using PDMS gasket pre-treatment and pre-wetting has the potential to improve assay yield by reducing the impact of bubbles by 99-100%. **(a)** Illustration showing the formation of a Harvey nucleus at a crevice in the microfluidic channel wall during pre-wetting and its subsequent growth, detachment, and re-growth. **(b)** Contact angle measurements depicting the hydrophobic recovery of PDMS after air plasma treatment show that the static contact angles for ultrapure water, ethanol, and 0.3 mM Triton X-100 solutions on PDMS remain <20° for ∼30-60 minutes after treatment. Baseline (pre-plasma) values are plotted as dashed lines. **(c-e)** quantification of bubble duration (top) and incidence (bottom) for the bubble-mitigation strategies. Beside each bubble duration bar plot, pie charts depict the assay yield (with successful assays defined as those with bubbles in the channels for <10% of the monitored time); dark grey pie slices denote successful assays. **(c)** Quantification of bubble incidence and duration shows that air plasma treatment mitigates bubble formation in PDMS microchannels, reducing bubble duration by 68%. Data are shown for non-plasma-treated and plasma-treated PDMS gaskets (not degassed) that were pre-wetted with 0.3 mM Triton X-100 (no plasma: n = 8, plasma: n = 16). **(d)** Quantification of bubble incidence and duration shows that PDMS degassing reduces bubble duration by a further 92% compared to plasma treatment when channels are pre-wetted with Triton X-100 solution, and reduces bubble duration by 100% for gaskets pre-wetted with ethanol (EtOH). Data are shown for non-degassed (Triton: n = 8, EtOH: n = 24) and degassed (Triton: n = 18, EtOH: n = 3) PDMS gaskets, either treated with plasma and pre-wetted with Triton X-100 solution or pre-wetted with ethanol (no plasma treatment). **(e)** Quantification of bubble incidence and duration comparing the performance of three different pre-wetting liquids. We compare pre-wetting with water (n = 8), ethanol (n = 24), and 0.3 mM Triton X-100 in PBS (n = 8) with no gasket degassing or plasma treatment, and also compare a gold-standard condition that uses pre-wetting in 0.3 mM Triton X-100 in PBS immediately after gasket degassing and plasma treatment (n = 18). Triton X-100, ethanol, and gold-standard Triton + plasma + degassing reduce bubble duration by 48%, 73%, and 99%, respectively, compared to water. In each of (c-e), we quantified the proportion of monitoring time where bubbles were visible, bubble introduction frequency, and bubble nucleation frequency in the microchannels during several hours of aqueous solution flow after pre-wetting (>2 h duration for all trials). ^*^p<0.05 in two-tailed Mann-Whitney U-tests. Further details regarding significance testing are provided in ESI S3. Significance indicators in bubble frequency plots match the colour of the dataset to which they refer.

**Fig. 4.**
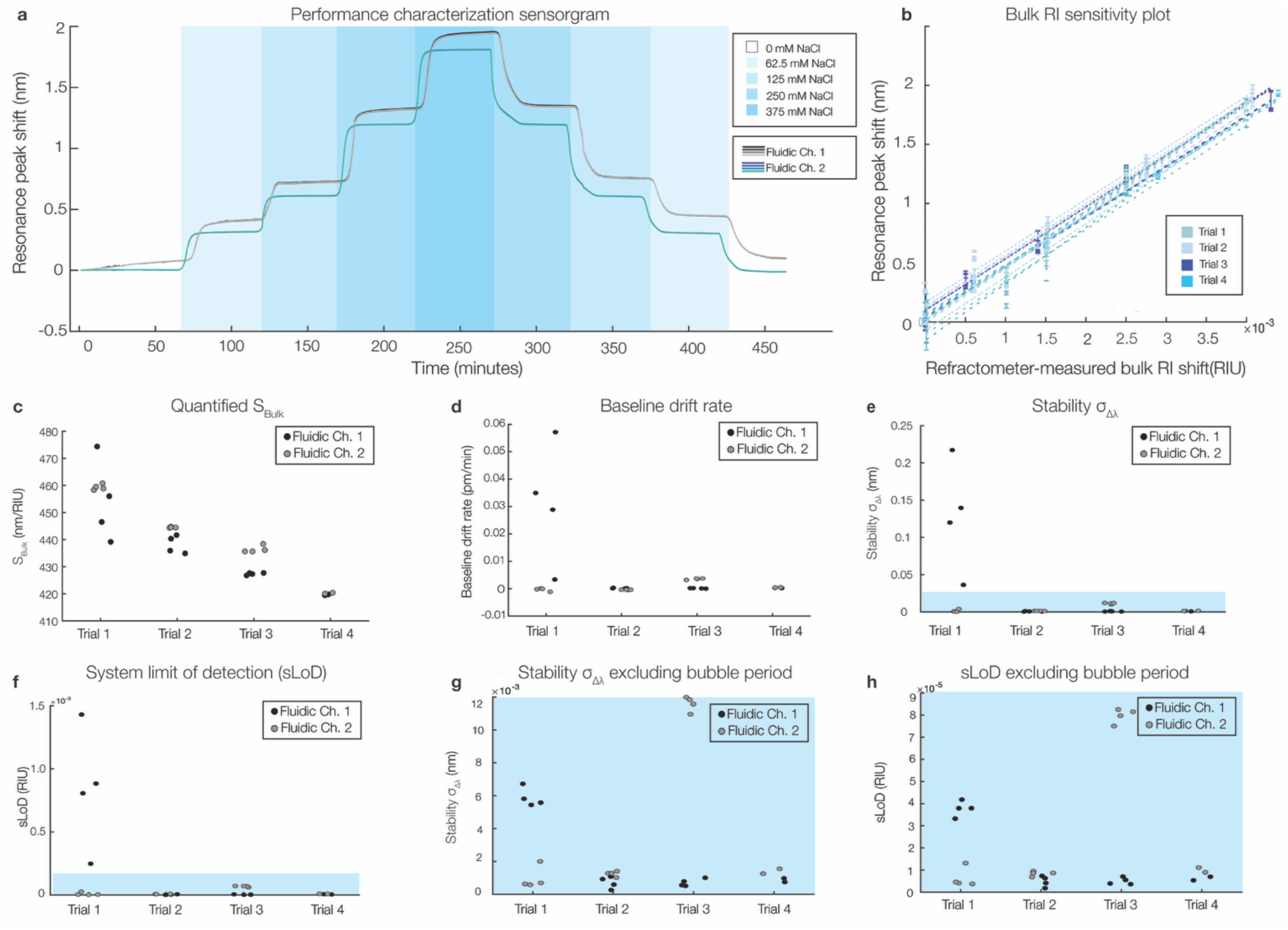
Intrinsic performance and replicability of SiP biosensor system. **(a)** Peak shift vs. time (sensorgram) plot for a representative performance characterization experiment, with overlaid regions highlighting bulk RI standard solution delivery. Eight lines (corresponding to 8 sensors) are plotted, with the signals from the 4 sensors in each channel tightly coinciding in this trial. **(b)** Bulk RI sensitivity plot (average peak shift vs. shift in standard solution RI), with the data from all sensors in all 4 trials (along with the linear fits used to extract the bulk RI sensitivity) overlaid. Error bars depict 3 standard deviations of the sensorgram peak shift signal in 25 measurements (∼3-5 mins) during exposure to each RI standard solution. **(c–f)** Beeswarm plots showing **(c)** quantified S_Bulk_, **(d)** baseline drift rate, **(e)** stability σ_Δλ_, and **(f)** sLoD over 4 trials, with 8 sensors/trial and using a 10-minute stability-monitoring period. A bubble that entered the second microfluidic channel during the stability monitoring period of trial 1 resulted in poor measured stability during that trial. **(g–h)** Re-analyzing the data to exclude the bubble-containing period from the analysis yields plots of **(g)** the stability σ_Δλ_ and **(h)** sLoD with much lower spread and better performance. Blue shading denotes the same y-axis range in (e) and (g) and in (f) and (h). Note that due to a data acquisition error, Trial 4 had only 4 sensors (2 sensors per channel).

Across the 8 replicate sensors in each trial and 4 replicate trials, we observed S_bulk_ values ranging from 419–474 nm/RIU. The measured stability σ_Δλ_ values show a much larger spread, ranging from 0.26– 217 pm. Most of this large spread is driven by Trial 1, where timelapse monitoring confirmed that a bubble passed over the channel 2 sensors during the stability-monitoring period (ESI Fig. S7). This bubble also affected the quantification of the baseline drift and sLoD. Re-analyzing the data to exclude the period containing this bubble yielded stability and sLoD data presented in Fig. 4(g-h), showing a drastic improvement in the stability σ_Δλ_, sLoD, and replicability. The average values for each metric after this re-analysis and their inter- and intra-assay coefficients of variation (CVs) are reported in Table 4.

**Table 4.**
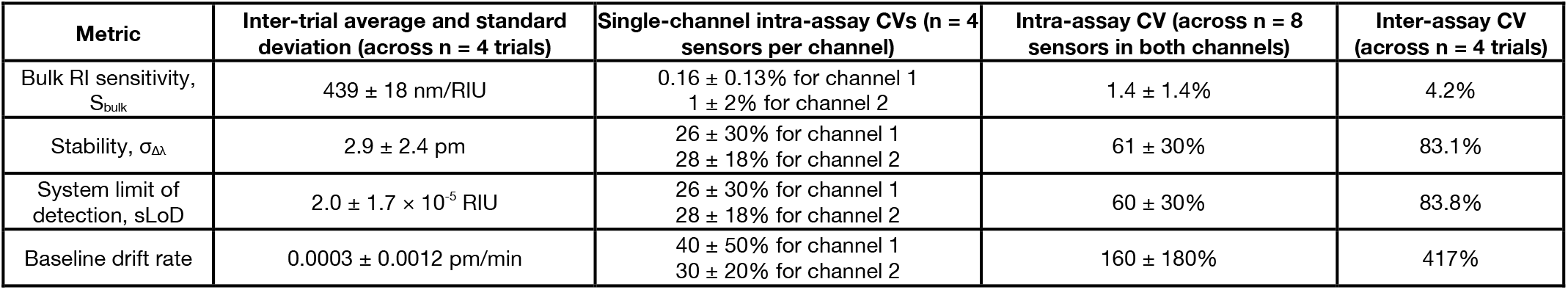
Quantified replicability of SiP biosensors for intrinsic performance metrics (data shown in Fig. 4; bubble-excluded data are reported in the table).

The inter- and intra-assay CVs for S_bulk_ are all low (<5%), with inter-assay CV approximately 3 times that of the average intra-assay CVs. This suggests that both variability between sensors (for example due to variation in fabricated SWG waveguide dimensions or due to how the crevices of these sensors are wetted with fluid) and variability between trials contribute to overall replicability. Variability between trials may be due to other factors noted in Tables 1–2, such as surface etching and material adsorption, bubbles, and reagent depletion or contamination. These factors might be expected to impact all 4 sensors in each microfluidic channel (all located within <1 mm of each other), and drive up the inter-assay and 2-channel intra-assay CVs compared to the single-channel intra-assay CV. Another factor that may contribute to the inter-assay variability reported here is the accuracy of our reference (refractometer) measurements of the bulk RI of our standard solutions. Each solution was measured prior to each assay in case pipetting errors or solution evaporation during storage resulted in slight variation of the solution RI; however, from the refractometer measurements presented in ESI S2.10, even the RI of the ultrapure water varied slightly across the trials (with standard deviation of 2.5 × 10^-4^ RIU across the 4 trials). This suggests that there was some variability in this reference measurement across different trials or perhaps different operators, and the small bulk RI shifts used in these trials were approaching the resolution of the measurement. Because the maximum difference in bulk RI between our standard solutions (ΔRI) was 4.4 × 10^-3^ RIU, the measurement variability (6% of the maximum ΔRI) may have had an appreciable effect on the inter-assay replicability. The absolute S_bulk_ values are subject to systematic error due to the use of a visible light Abbe refractometer for our reference measurements while our sensors operate near 1550 nm. However, all sensor measurements are subject to the same error in S_bulk_ measurement so this would not be expected to impact the measured CVs. A comparison between our measured refractive index step sizes and predicted values at 1550 nm from a previously reported model (91) is presented in Table S4; this comparison shows that all predicted 1550 nm step sizes lie within two standard deviations of our measured visible light step sizes.

Nevertheless, our reported S_bulk_ replicability across these trials is similar to that reported previously in the literature. Analyzing strip waveguide ring resonator sensors, Iqbal *et al*. report an S_bulk_ of 163 nm/RIU with 3.9% intra-trial (ring-to-ring) CV in the resonance shifts magnitude resulting from a bulk RI shift, as well as a cycle-cycle variance of 0.47% (the same ring resonators exposed to multiple cycles of bulk RI shifts) (92). Using the commercial Genalyte system with 128 ring resonator sensors, Mudumba *et al*. tracked the,resonance shift due to a bulk RI increase resulting from the addition of 0.5 M NaCl to the buffer, and report intra-trial CVs in the averaging 0.87% (ranging from 0.5–1.8% over 192 chips) (49). Chip-to-chip inter-assay CVs in the same work ranged from 0.4–1.8%. These previous results, using strip waveguides that might be expected to be less sensitive to variations in fabrication uniformity and wetting than the SWGs used here, are similar in magnitude to our results.

The inter-assay CVs for the stability σ_Δλ_, sLoD, and baseline drift rate are all considerably larger and reflect the variability in stability and noise levels from trial to trial. Removing the impact of the bubbles reduced the inter-assay CVs in the stability σ_Δλ_ and sLoD from 170% and 169% to 83.1% and 83.8%, respectively. The average stability σ_Δλ_ was reduced from 18.3 ± 31 pm to 2.9 ± 2.4 pm, illustrating how the presence of bubbles can have a drastic effect on both the stability and the replicability of these types of sensors. Nevertheless, even without the presence of bubbles, the CVs of these stability metrics are much larger than those for S_bulk_. This is likely due to the overall small magnitude of the stability σ_Δλ_ and baseline drift rate and the stochastic nature of the noise effects that impact stability. Because the baseline drift rate in particular is so small (ranging from −2.1 × 10^-3^ to 3.8 × 10^-3^ pm/min), even the largest-magnitude slope would only contribute to a ∼0.1 pm resonance shift over a ∼20–30-min binding step. This contribution is small both in comparison to typical shifts observed during bulk RI and binding assays (>100 pm) and also in comparison to the overall stability σ_Δλ_.

Next, we sought to characterize the performance and replicability of these same SiP devices as specific analyte sensors. We hypothesized that these same sensors’ replicability for performance metrics relevant to biomolecule detection would be poorer than that for intrinsic performance metrics due to the larger set of factors that impact these metrics, as described in Table 2. Comparing different metrics of performance and replicability has the potential to help isolate factors that can be subsequently tuned to improve the ultimate performance of an assay. To test this hypothesis, we exposed the same sets of sensors used for intrinsic performance characterization to a demonstration binding assay to detect the spike protein of the SARS-CoV-2 virus (20 µg/mL in PBS). We used a microfluidic flow-based functionalization process employing noncovalent Protein A-mediated capture antibody immobilization for these measurements so that we could extract and compare the sensor signal during each stage of functionalization.

The biomolecule-detection performance of our sensors is presented in Fig. 5. Fig. 5(a) depicts a representative sensorgram showing the assay stages and resulting peak shift signal of the 8 replicate sensors across the duplicate microfluidic channels for a single trial, encompassing sensor functionalization and spike protein detection. Clear peak shift signal is visible for each stage of sensor functionalization and analyte detection. Quantifying this peak shift signal yields the data shown in Fig. 5(b–e), in which the shifts for each of the 8 sensors across the two duplicate channels are plotted for the Protein A, BSA, capture antibody functionalization, and spike protein detection assay stages. Across the 4 replicate trials, we measured average peak shifts (± one standard deviation) of 1.5 ± 0.3 nm as the Protein A adsorbed to the waveguide surface, 0.21 ± 0.18 nm as BSA nonspecifically attached to the surface to block it, 0.7 ± 0.3 nm as the capture antibody bound to the Protein A on the sensor surface, and 0.84 ± 0.15 nm as the spike protein bound to the functionalized surface. Quantifying the sensor drift at the beginning and end of the assay yields the results shown in Fig. 5(f–g), with an average initial baseline drift rate of 0.6 ± 1.2 pm/min, and an average final drift rate of −3.4 ± 1.6 pm/min.

**Fig. 5.**
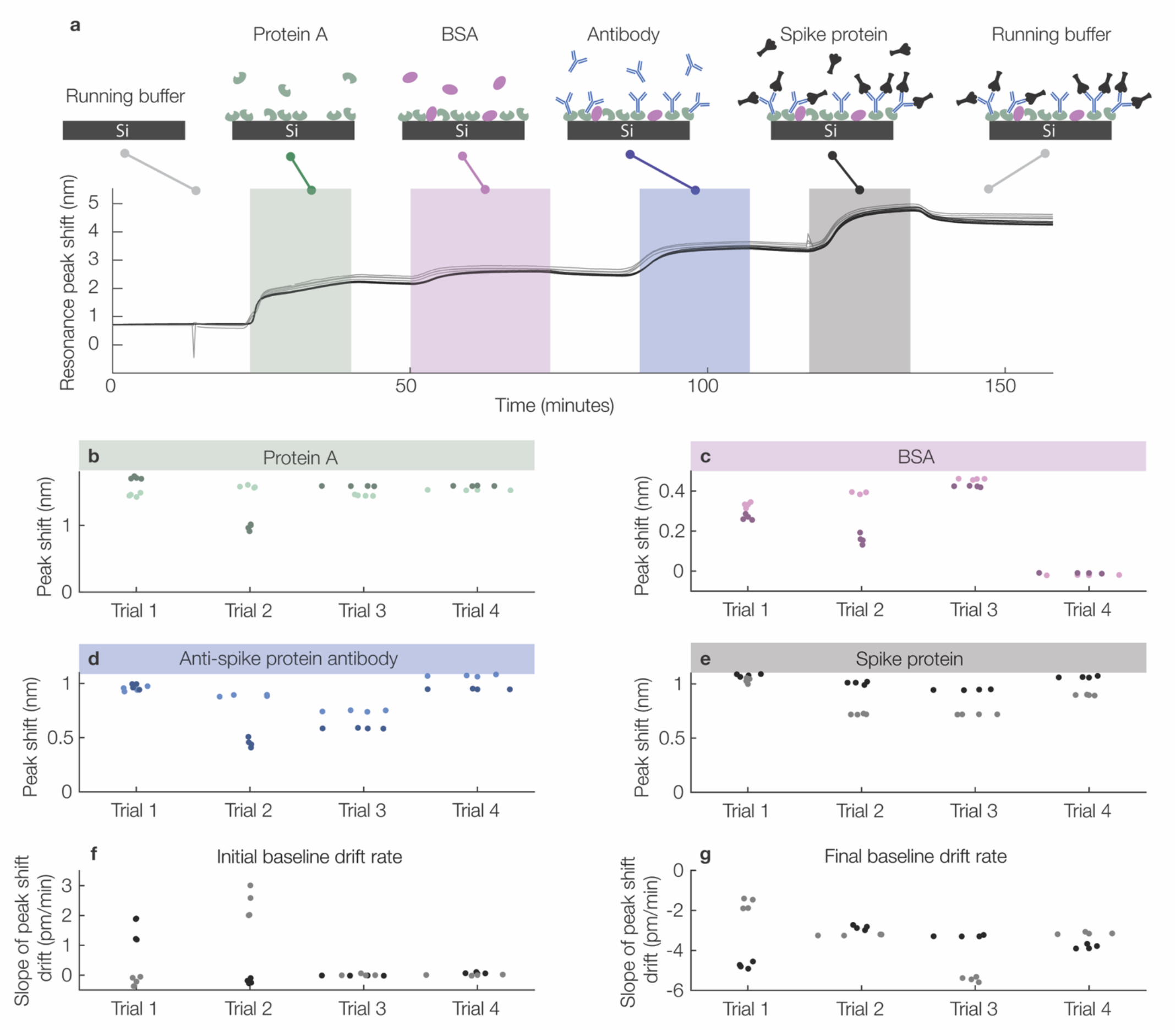
Performance and replicability of SiP biosensor analyte detection reveals our sensors’ analyte-detection replicability with 11.1% inter-assay CV using a multi-step oriented bioaffinity-based (non-covalent) biofunctionalization approach, reflecting the increased number of factors impacting analyte detection in comparison to intrinsic bulk sensitivity. Analyte-detection replicability exceeds that of functionalization (antibody-attachment) shifts. **(a)** Sensorgram plot for example binding assay, illustrating the assay stages. **(b–e)** Beeswarm plots showing quantified peak shifts for each of the 8 sensors/trial across two microfluidic channels per trial and 4 trials. **(b)** Quantified Protein A adsorption shifts. **(c)** Quantified nonspecific bovine serum albumin (BSA) binding shifts. **(d)** Quantified capture antibody binding shifts.**(e)** Quantified SARS-CoV-2 spike protein binding shifts. **(f)** Beeswarm plot showing quantified initial baseline drift rates during the initial PBS stability region, after baseline correction. **(g)** Beeswarm plot showing quantified final baseline drift rates after spike protein binding, after baseline correction. Trial 1 is shown in (a). Line and datapoint colours denote the sensors located in each of two microfluidic channels run with identical assays, with the darker colours consistently denoting the same microfluidic channel (#1) across subplots.

We subsequently analyzed the peak shift and baseline drift data presented in Fig. 5 to quantify the CVs of the sensor signals in response to each binding stage as a measure of replicability. We quantified inter-assay CV as well as intra-assay CV across both microfluidic channels for each trial and the separate intra-assay CVs for each microfluidic channel and trial; these results are summarized in Table 5.

**Table 5.** Quantified replicability of SiP biosensors for functionalization and biomarker detection using an example binding assay with microfluidic flow-mediated functionalization (data shown in Fig. 5). For all intra-assay CV values, the mean and standard deviation of the intra-assay CVs in 4 trials are reported. ^*^Some trials showed very small initial baseline drift rates (e.g., 0.0009 ± 0.016 pm/min for the Trial 4 channel 2 intra-assay CV), which led to spuriously high inter- and intra-assay CV values.

| Assay stage | Inter-trial average and standard deviation (across n = 4 trials) | Single-channel intra-assay CVs (across n = 4 sensors in each channel) | Intra-assay CV (across n = 8 sensors in both channels) | Inter-assay CV (across n = 4 trials) |
| --- | --- | --- | --- | --- |
| Protein A adsorption | $1.48 \pm 0.14$ nm | $1.0 \pm 0.7\%$ for channel 1<br>$1.5 \pm 2.2\%$ for channel 2 | $10 \pm 10\%$ | 9.5% |
| BSA blocking/nonspecific adsorption | $0.25 \pm 0.19$ nm | $2.7 \pm 2.0\%$ for channel 1<br>$10 \pm 8\%$ for channel 2 | $24 \pm 20\%$ | 76.5% |
| Capture antibody functionalization | $0.83 \pm 0.18$ nm | $1.2 \pm 0.7\%$ for channel 1<br>$3 \pm 4\%$ for channel 2 | $14 \pm 14\%$ | 22.3% |
| Spike protein detection | $0.93 \pm 0.10$ nm | $0.9 \pm 0.4\%$ for channel 1<br>$0.9 \pm 1.0\%$ for channel 2 | $11 \pm 7\%$ | 11.1% |
| Initial baseline drift* | $0.45 \pm 0.53$ pm/min | $31 \pm 7\%$ for channel 1<br>$500 \pm 800\%$ for channel 2 | $300 \pm 300\%$ | 117.7% |
| Final baseline drift | $-3.5 \pm 0.6$ pm/min | $2.8 \pm 1.2\%$ for channel 1<br>$5 \pm 7\%$ for channel 2 | $24 \pm 20\%$ | 16.6% |

As shown across most metrics in Table 5, the single-channel intra-assay CVs (replicate sensors within a single microfluidic channel) are nearly an order of magnitude lower than the intra-assay and inter-assay CVs across both replicate channels. This suggests that for the analyte-detection performance metrics presented in Table 5, the factors tabulated in Tables 1–2 that differ between microfluidic channels might play a larger role in the overall inter-assay replicability in comparison to intra-assay variation between sensors or other inter-trial factors like the preparation of the protein solutions that do not vary between channels (since the same protein solutions were split between the two microfluidic channels in each trial). These channel-specific factors include those related to the pre-wetting process or the cleanliness and performance of the microfluidic system that may result in reagent depletion within the tubing before it reaches the sensors, dilution, or contamination. The exceptions to this finding are the initial baseline drift and BSA blocking assay stages, where other factors may have dominated the observed variability.

We do not observe any consistent significant relationship between spike protein detection signal and sensor position along the length of the channel (ESI S5), suggesting that analyte depletion along the length of the channel likely does not contribute to the intra-assay CV for this relatively high analyte concentration. The effects of analyte depletion are expected to become more prominent at lower analyte concentrations (53), contributing to increased variability. As highlighted in Fig. 1(a), microfluidic channel alignment has the potential to impact analyte-detection signals due to the impact of nonuniform fluid velocity across the width of the channel on convective mass transport to the sensor surface. In our system, however, our flow rate of 30 µL/min yields a Péclet number (Pe_H_) of 2.5 ×10^4^ (ESI S6), indicating that mass transport in our channels during the spike protein binding stage is diffusion-limited, and variations in flow rate across the channel width are unlikely to result in a large impact on the overall mass transport to the sensor surface. Depletion of small-molecule compounds due to sorption into and onto peristaltic pump tubing has been previously found to dominate over PDMS device sorption (50); although we use different tubing materials in this work (FEP and PEEK) it is possible that biomolecule depletion (either during the functionalization or detection stage of the assay) due to protein adsorption to the tubing walls before it reaches the chip impacts our sensing performance and replicability.

The 11.1% inter-assay CV for spike protein detection observed here is comparable to or lower than the typical CV range reported for commercial enzyme-linked immunosorbent assays (∼10–20%), and lower than the 20% CV threshold commonly employed for immunoassay validation (12,93,94), suggesting acceptable replicability for analyte detection. Our replicability is also comparable to previously published analyte measurements using SiP biosensors (ESI Table S1), although many works do not report the replicability of their sensors. A demonstration of an immersible SiP biosensor used for spike protein serology (detection of anti-spike protein antibody after functionalization of the chip with the spike protein’s receptor binding domain (RBD)) reported intra-assay CVs ranging from 3.6–5.7%, and inter-assay CVs of 6.9–8.6% from duplicate measurements (95). A demonstration assay interrogating the biotin-neutravidin interaction using an 8-channel silicon nitride ring resonator system reported intra-assay CVs of 10% and 13.9% and inter-assay CVs of 13.9% and 11% for neutravidin analyte concentrations of 50 and 10 μg/mL, respectively (96). Inter-assay CVs for calibration fit parameters for a hybridization assay to detect microRNA-21 using bimodal waveguide interferometers were reported as 5.3–5.7% in simple buffer and 2.6–3.4% in undiluted plasma, while inter-assay CVs for the assay LoDs were 51% in buffer and 17% in plasma (97). In a large study using the commercial Genalyte platform (which has 128 ring resonator sensors in 32 clusters) to detect anti-SS-A antibodies, CVs were reported across all 128 rings (2.4% average, ranging from 0.6–6.4%), within each cluster (ranging from 0.04–16%), across clusters (average 2.2%, ranging from 0.3–6.1%), and across chips (averaging 5% CV across 72 chips) (49). This comparison shows that the SWG sensors employed in our study show comparable replicability to those used in previous works, and our comparison of the inter- and intra-assay CVs shows that the sensors themselves are likely not the limiting factor for assay replicability.

Higher CVs are observed in our experimental data for the BSA blocking/nonspecific binding test (CV = 76.5%) as well as the capture antibody attachment stage (CV = 22.3%) of the assay. The variability in the BSA stage can be partly attributed to the low overall shifts observed during that binding stage, but also results from very low/negligible BSA signal in the fourth trial. The variability in the signal from antibody attachment may be due to antibody aggregates or other factors that led to different degrees of mass attachment during this binding stage without a large effect on analyte-detection performance.

The initial baseline drift tended to have very high CVs alongside measured drift slopes <3 pm/min. Low average drift slopes resulted in very high intra-assay CVs in some cases. There was also variation between channels and between trials, with some assays showing a slight positive initial baseline drift and others showing a slight negative initial baseline drift as shown in Fig. 5. The baseline drift should ideally be zero; however, baseline drift in the resonance peak position of SiP biosensors exposed to PBS solutions is well documented and can arise from factors such as waveguide etching (98,99), silicon oxidation (100,101), or deposition or removal of residues from the surface of the waveguide due to the microfluidic system. Oxidation, etching, or removal of previous residues would be expected to result in a negative baseline drift, while deposition of residue would be expected to result in a positive baseline drift. Indeed, we observed a larger magnitude of baseline drift rate in PBS at the start of the assay as compared to the baseline drift slope in water reported in Fig. 4, which may reflect the slow etching of silicon that can occur in PBS buffer solution (98,99). Baseline correction was used to reduce the effects of drifts on the assay results; however, some drift remained after correction in these trials. This remaining slope in the resonance peak shift data suggests that the drift was not completely consistent/linear over time; however, the magnitudes were small in comparison to the analyte-detection shifts. The use of ethanol pre-wetting for these trials, alongside long exposures to fluids prior to the start of the immunoassay (with the intrinsic performance characterization as well as an overnight incubation in flowing water), may have contributed to residue deposition on the sensor surface before and during this beginning stage of the immunoassay. These types of residues, depending on the type and time of deposition (whether material was attaching to or detaching from the waveguide during the initial stability monitoring period), could introduce either positive or negative baseline drift rates. Because the ethanol pre-wetting process was conducted manually, it may be susceptible to increased inter-experiment variability in potential residue deposition from syringe and tubing materials, syringe lubricants, and salt precipitation. The final baseline drifts are consistent in negative slope direction, larger in magnitude than the initial drifts, and more replicable between trials with 16% inter-assay CV. We expect that these baseline drifts are dominated by the dissociation kinetics and desorption of the biomolecules involved in the binding assay. Some dissociation between antibody and antigen will always be expected after switching to a solution no longer containing free antigen; this dissociation is governed by the dissociation rate constant for the antibody-antigen pair (102). Desorption of non-covalently immobilized Protein A, capture antibody, and BSA are also expected to contribute to these final baseline drifts.

Overall, our experimental characterization of SWG sensors shows spike protein detection signal replicability comparable to similar immunoassays. As expected, the CVs for spike protein detection are higher than those for bulk RI sensitivity, reflecting the greater number of factors that contribute to the variability in this metric (Table 2), although still well within an acceptable range for commercial immunoassays (12,93,94). However, our analysis reveals that we typically see more variation in the stability and baseline drift metrics, the nonspecific binding signal, and the antibody attachment signal than in the analyte-detection signal. CVs in these other metrics are reported less frequently than those for analyte detection and bulk RI sensitivity. It is possible that prior to functionalization, the bare sensor is particularly sensitive to the state of the fluidic system and any potential contamination or etching that could be introduced, leading to higher variation in the stability and baseline drift from trial to trial and between microfluidic channels. We hypothesized that our functionalization and assay design could be modified to improve replicability, e.g., by functionalizing the sensors prior to microfluidic integration (55,103), at the cost of no longer being able to monitor the sensor during each stage of functionalization.

### 3.3. Functionalization strategy impacts analyte detection

Having characterized our sensors’ intrinsic and analyte detection performance and replicability in a single assay format, we sought to quantify the effect of sensor surface functionalization on analyte detection performance and replicability. We aimed to evaluate the effect of two aspects of functionalization: bioreceptor (1) immobilization chemistry and (2) patterning technique. The bioreceptor choice (antibody) was kept constant. Regarding immobilization chemistry, we sought to compare noncovalent and covalent approaches. We hypothesized that, compared to the noncovalent Protein A-mediated antibody immobilization previously used by our group (100,104,105) and discussed in Section 3.2, covalent PDA-mediated immobilization may offer improved sensor stability, replicability, and sensitivity (45,65,106). PDA is a bioinspired surface coating that can be deposited on various surfaces from simple one-pot room temperature aqueous reactions to yield nanometer-scale films (107). Under neutral pH, PDA contains quinone groups that are susceptible to nucleophilic attack by amines and thiols present on proteins through Michael addition and Schiff base reactions, facilitating direct unoriented covalent attachment of protein-based bioreceptors (108,109). Bakshi et al. (46,110) recently demonstrated covalent antibody immobilization on SiP sensors using PDA chemistry, which improved sensor yield compared to common silane-based approaches that typically involve moisture-sensitive reactions and tight process controls (27,111,112). Unlike Protein A-mediated antibody immobilization, covalent approaches offer more stable bioreceptor attachment, which can enable sensor regeneration for reuse; this is valuable for both assay development in the lab and serial biomarker measurements in end-use settings (113,114). Regarding bioreceptor patterning technique, we wished to compare in-flow antibody application, as previously used by our group (100,104,105) and discussed in Section 3.2, and spotting-mediated application. Compared to in-flow patterning, spotting-mediated antibody application may address channel-level factors that dominate variability in the flow-based approach. Moreover, spotting-mediated functionalization, combined with inkjet or pin printing technologies, can offer improved patterning resolution for multiplexed surface functionalization toward multi-analyte detection on a single sensor chip and also permits the use of higher antibody concentrations due to reduced fluid consumption (27).

To compare these different functionalization approaches, we performed SARS-CoV-2 spike protein demonstration immunoassays on sensors functionalized via noncovalent Protein A-based immobilization chemistry + in-flow antibody application (Protein A/flow), covalent PDA-based immobilization chemistry + spotting-based antibody application (PDA/spotting), and covalent PDA-based immobilization chemistry + in-flow antibody application (PDA/flow). Fig. 6 illustrates each functionalization approach and assay design. Detection of 1 µg/mL spike protein was performed on sensors functionalized using all three strategies, while detection of 20 µg/mL spike protein was also demonstrated for Protein A/flow functionalization, as discussed in Section 3.2. For the covalent functionalization approaches (PDA/spotting and PDA/flow), we compared the effect of antibody patterning technique on intra-assay performance (initial and final baseline drift rates, BSA challenge peak shifts, and spike protein detection peak shifts) and replicability (intra-assay CVs); results are shown in Fig. 7. We also compared all three functionalization approaches (Protein A/flow, PDA/spotting, and PDA/flow) in terms of inter-assay performance and replicability (inter-assay CVs); results are shown in Fig. 8.

**Fig. 6.**
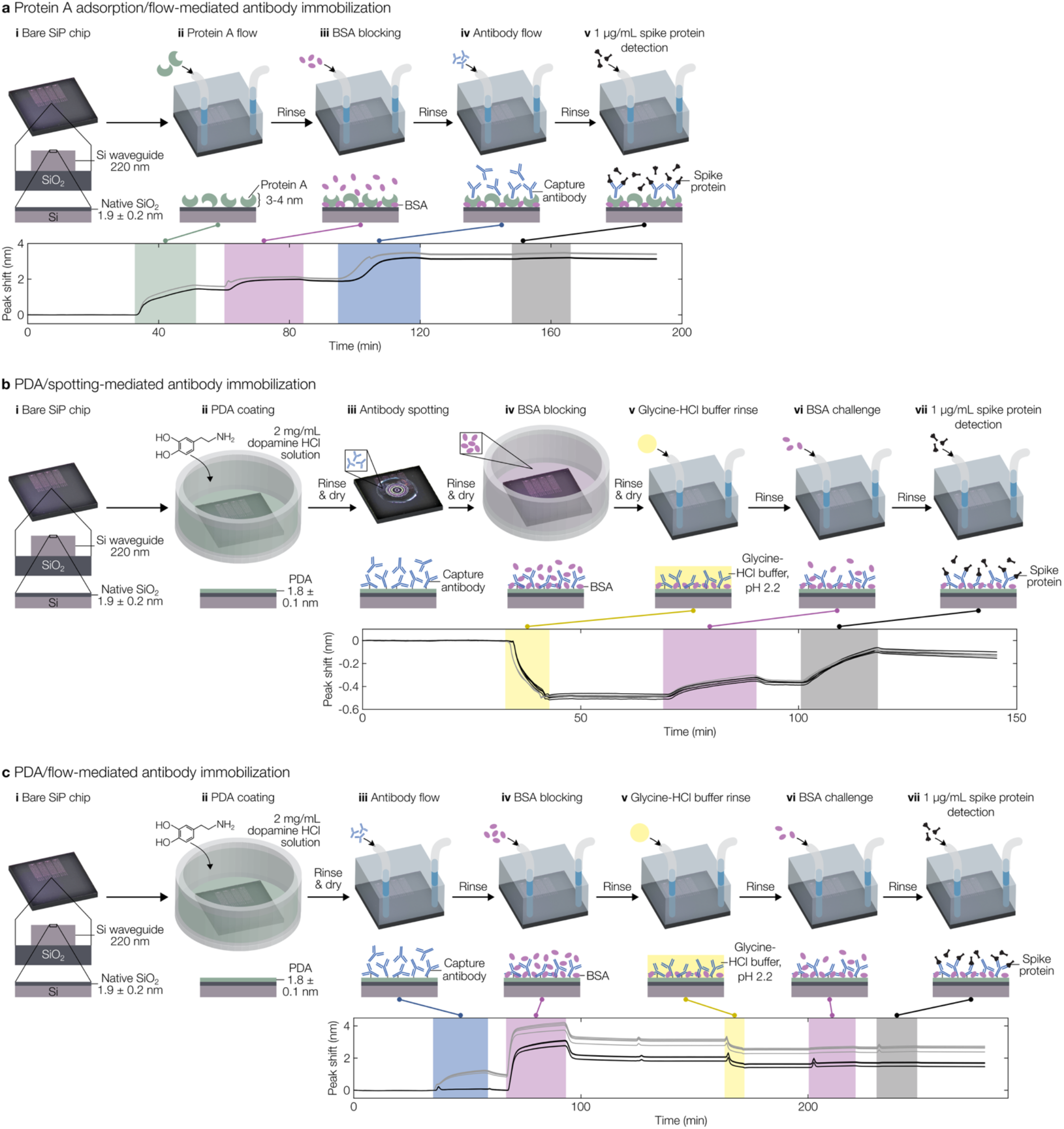
Functionalization strategy comparison, depicting the steps of functionalization and analyte detection performed prior to and after fluidic integration for each strategy. **(a)** Protein A adsorption/flow-, **(b)** PDA/spotting-, and **(c)** PDA/flow-mediated antibody immobilization. Each illustration depicts the functionalization process flow, assay steps, and a representative experimental peak shift trace. For each peak shift trace, data from 8 sensors located in two microfluidic channels that were exposed to identical assay conditions are plotted. Line colours denote the sensors located in each of two microfluidic channels, with the darker colours consistently denoting the same microfluidic channel across subplots. A spike protein concentration of 1 µg/mL was used in all assays depicted in these representative traces. Peak shift traces highlight the potential improvements to analyte detection sensitivity and inter-channel replicability offered by performing functionalization prior to fluidic integration.

**Fig. 7.**
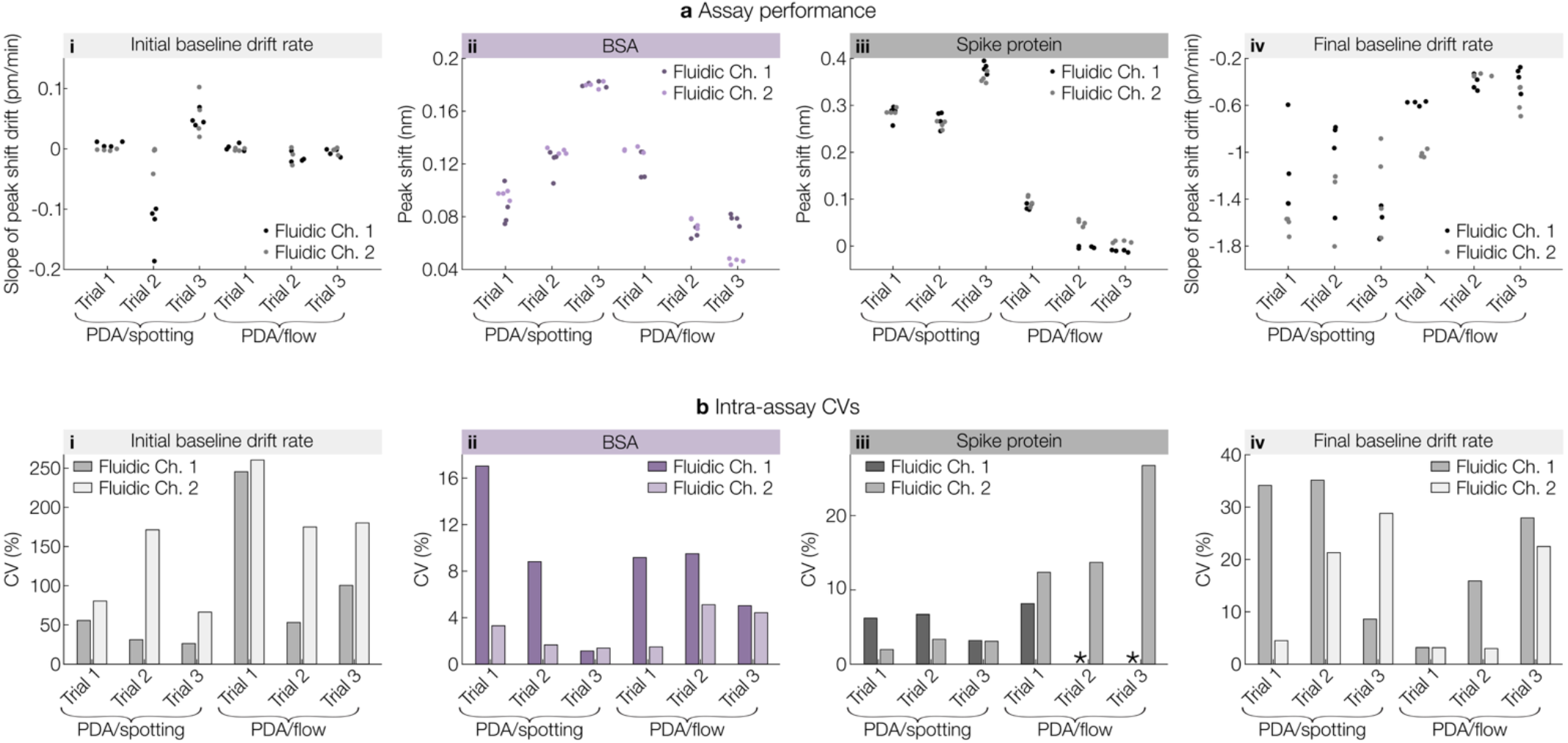
The antibody immobilization strategy (spotting versus in-flow patterning) impacts the performance and replicability of PDA-modified SiP biosensors in SARS-CoV-2 spike protein demonstration assays, with spotting at high antibody concentration facilitating higher analyte detection signals than flow-based deposition using equivalent antibody consumption (lower concentration). **(a)** Beeswarm plots comparing quantified assay performance metrics, including **(i)** initial baseline drift rate, **(ii)** BSA binding shift, **(iii)** spike protein binding shift, and **(iv)** final baseline drift rate. Data were collected from eight sensors/trial across two microfluidic channels/trial and three trials/functionalization approach. All metrics were quantified from baseline-corrected data. **(b)** Clustered bar charts comparing intra-assay CVs across sensors in each fluidic channel for quantified assay performance metrics, including **(i)** initial baseline drift rate, **(ii)** BSA binding shift, **(iii)** spike protein binding shift, and **(iv)** final baseline drift rate. ^*^CVs are not shown for fluidic channel 1 in trials 2 and 3 for PDA/flow due to negligible mean binding shifts, which led to spuriously high CV values.

**Fig. 8.**
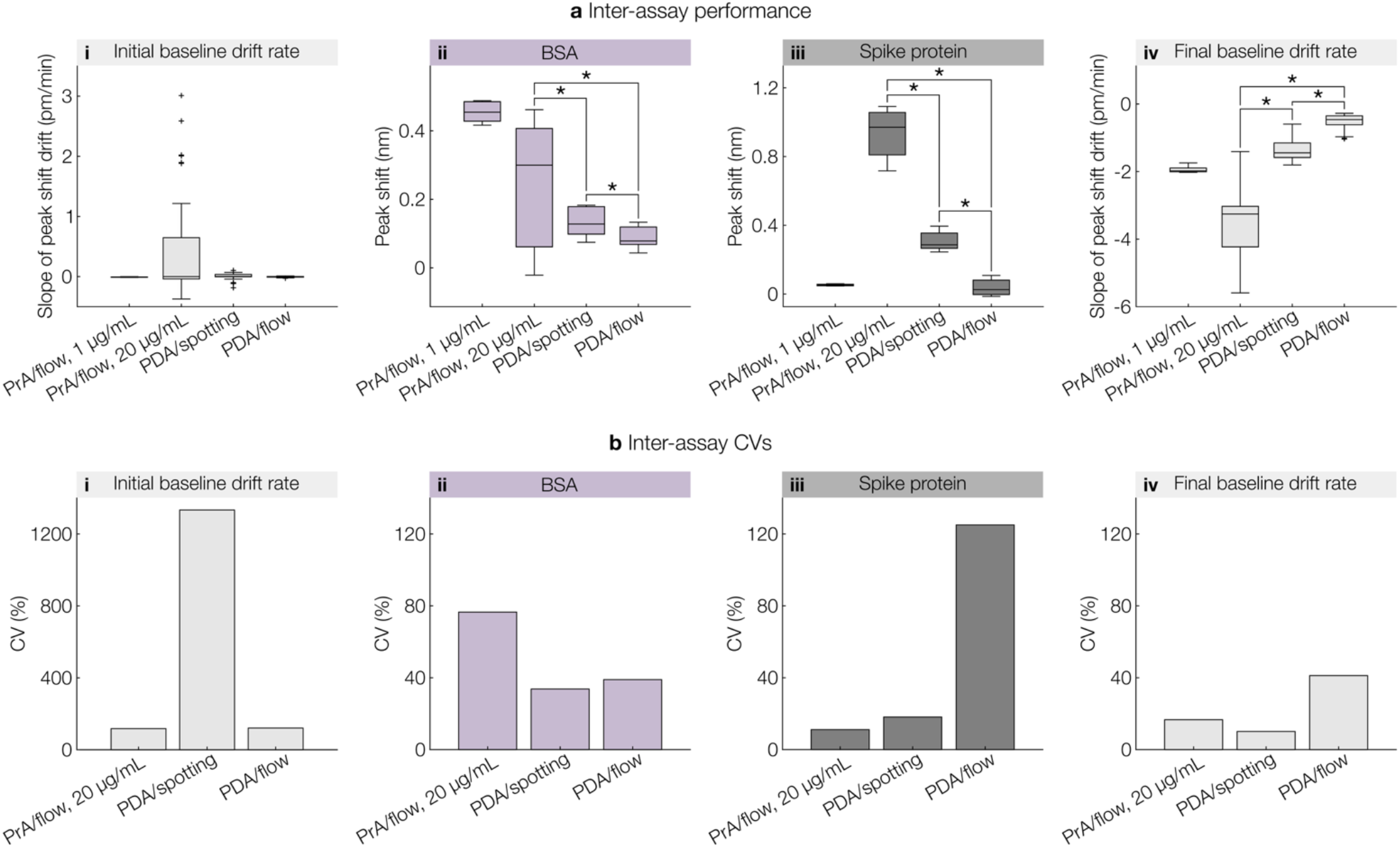
Impact of functionalization approach on the inter-assay performance and replicability of SiP biosensors in SARS-CoV-2 spike protein demonstration assays.**(a)** Box and whisker plots comparing quantified performance metrics for assays performed on sensors functionalized via Protein A adsorption/flow-, PDA/spotting-, and PDA/flow-mediated antibody immobilization. Plotted performance metrics include: **(i)** initial baseline drift rate, **(ii)** BSA binding shift, **(iii)** spike protein binding shift, and **(iv)** final baseline drift rate. Visualized data include the sensor response of eight sensors/trial across two microfluidic channels/trial. Data are shown for assays performed with a 1 µg/mL spike protein concentration for the Protein A/flow (1 trial), PDA/spotting (3 trials), and PDA/flow (3 trials) functionalization approaches. Data are also shown for assays performed using a 20 µg/mL spike protein concentration for the Protein A/flow functionalization approach (4 trials). All metrics were quantified from baseline-corrected data. ^*^p<0.05 in two-tailed Mann-Whitney U-tests. Further details regarding significance testing are provided in ESI S3. **(b)** Bar charts comparing inter-assay CVs for assays performed on sensors functionalized via Protein A/flow-, PDA/spotting-, and PDA/flow-mediated antibody immobilization. Inter-assay CVs are shown for the following metrics: **(i)** initial baseline drift rate, **(ii)** BSA binding shift, **(iii)** spike protein binding shift, and **(iv)** final baseline drift rate.

Initial drift rates were close to zero for all assays detecting 1 µg/mL spike protein, regardless of functionalization approach, indicating good performance of the baseline corrections performed during post processing. Initial drift rates of −0.0080 pm/min, −0.0046 ± 0.0614 pm/min, and −0.0051 ± 0.0062 pm/min were measured for the Protein A/flow, PDA/spotting, and PDA/flow functionalization strategies, respectively (values are reported as inter-assay mean ± standard deviation where multiple trials were performed). For the Protein A/flow assays demonstrating 20 µg/mL spike protein detection, a mean initial drift rate of 0.45 ± 0.53 pm/min was measured, representing a larger drift rate and variability compared to the other assay formats. This may be attributed to setup protocols, which included ethanol pre-wetting and long exposures to fluids prior to the start of the immunoassay, as discussed in Section 3.2. In contrast, all other assays included Triton X-100 solution pre-wetting, informed by the results of Section 3.1, and only short exposures (∼1 hour) to fluids prior to the start of the immunoassay. Inter-assay CVs for initial drift rate are high for all assay formats (Fig. 8(bi)), consistent with the findings of Section 3.2. Initial drift rate CVs for the PDA/spotting and PDA/flow assays may be considered spuriously high due to the exceptionally small mean drift rates by which the standard deviations are normalized for this assay metric (115). Low initial drift rates for the assays employing PDA-mediated antibody immobilization (in which the sensor is already functionalized by this stage of the assay) also show that the stability of the functionalization chemistry and blocking is not a dominant factor leading to high drifts with unpredictable baselines that might be expected to degrade analyte-detection performance.

Average peak shifts due to nonspecific BSA adsorption were significantly greater (p<0.05, Mann-Whitney U test, ESI S3) for assays employing Protein A/flow functionalization (0.29 ± 0.19 nm, averaged across all five Protein A/flow assays) compared to those employing PDA/spotting (0.13 ± 0.04 nm) and PDA/flow (0.087 ± 0.034 nm) functionalization. In the Protein A/flow assays, these peak shifts correspond to in-flow blocking of Protein A-coated sensors with 1 mg/mL BSA. In the PDA-based functionalization assays, these represent peak shifts due to exposure of previously blocked sensors to 1 mg/mL BSA. The lack of prior surface blocking in the Protein A/flow assays likely contributed to more BSA adsorption (65). The Protein A/flow assay format involving 20 µg/mL spike protein detection also exhibited the greatest inter-assay CV for nonspecific BSA binding (77%). This may be related to inter-assay variations in sensor surface cleanliness and hydrophobicity following ethanol pre-wetting and fluid exposure prior to the assay (116). Among the assays employing covalent PDA-based functionalization, intra-assay variability was smaller than inter-assay variability for peak shifts due to nonspecific BSA adsorption. For PDA/spotting, the intra-assay CVs (across n = 8 sensors in both channels) were 6.7 ± 5.5%, while the inter-assay CV was 34%. For PDA/flow, the intra-assay CVs were 14 ± 11%, while the inter-assay CV was 39%. This suggests that procedural variations from day-to-day (e.g., variations in BSA stock solutions due to freeze/thaw cycles, timings of procedure steps, PDA coating, antibody spotting, and blocking performed on each chip, and pipetting accuracy) have a greater impact on nonspecific adsorption than differences between fluidic channels (e.g., flow rate stability and fluidic channel alignment) and individual sensors (e.g., fabrication differences or nonuniformities in surface cleanliness, wetting, and functionalization chemistry across the chip) within a given trial. The PDA/spotting assays exhibited greater nonspecific BSA adsorption shifts than PDA/flow, which may be attributed to the off-chip surface blocking employed in the PDA/spotting functionalization approach. Here, immersion-based blocking with BSA was performed prior to rinsing and drying the chip, mounting fluidics, pre-wetting the fluidic channels, and aligning optical I/Os. Between blocking the sensor and starting the binding assay, the sensor was exposed to air (i.e., during microfluidic gasket alignment) and flowing buffers (i.e., during pre-wetting and optical alignment) for variable amounts of time. Moreover, the pre-wetting buffer included surfactant, which can disrupt protein-surface interactions and solubilize proteins (117). These drying and buffer flow steps may have increased BSA dissociation from the surface, reducing blocking efficacy. On the other hand, in the PDA/flow assays, the sensors remained hydrated and were exposed to identical fluidic protocols between surface blocking and the binding assay, potentially reducing nonspecific binding.

Average peak shifts for specific detection of 1 µg/mL spike protein varied across functionalization approaches, with PDA/spotting yielding the greatest mean spike protein detection signals (0.31 ± 0.06 nm) compared to PDA/flow (0.038 ± 0.047 nm) and Protein A/flow (0.053 nm). Regarding immobilization chemistry, the low overall shifts as well as the low number of replicates challenge statistical comparison; however, the slightly greater analyte detection peak shifts achieved for Protein A/flow than PDA/flow may be related to the well-oriented nature of antibodies immobilized on Protein A (118,119). In contrast, covalent immobilization via PDA targets amine groups that are abundant on the antibody surface and may result in the formation of multiple bonds between the antibody and PDA film (107,108,120). This may cause conformational changes and render some binding sites inaccessible for spike protein capture.

We find large differences in performance due to antibody patterning technique. PDA-modified sensors functionalized via spotting yielded significantly larger analyte detection signals than in-flow antibody application (p = 3.1 ×10^-9^, Mann-Whitney U test, ESI S3). The PDA/flow assays also exhibited a greater inter-assay CV for spike protein detection. These differences in analyte detection performance suggest differences in the surface density and/or affinity of capture antibodies immobilized via these two application methods. These differences in capture antibody surface density and affinity may result from a combination of the antibody concentration (500 µg/mL in spotting-based application vs. 20 µg/mL in flow-based deposition, although the total antibody consumption was the same for the two approaches) as well as other factors related to flow-based delivery. We performed order-of-magnitude mass transfer calculations to model antibody delivery to the sensor surface via spotting- and flow-based capture antibody application (53) (ESI S6). In the PDA/spotting method, mass transfer of antibody molecules to the sensor surface is assumed to be solely driven by diffusion, while in the PDA/flow method, it is driven by diffusion and convection. We found that mass transfer in both protocols should be sufficient to provide >10×excess antibody molecules to the sensor surface than what is required to fully saturate the surface with receptors. This suggests that other experimental factors likely impacted antibody immobilization and subsequent analyte detection. During flow-based immobilization, antibody adsorption to the walls of the microfluidic tubing upstream of the SiP sensor may deplete antibody molecules from solution. In turn, this reduction in antibody concentration may slow mass transport kinetics, potentially leading to a lower surface concentration of immobilized capture antibodies. These slower kinetics may also render capture antibodies more susceptible to conformational changes due to stronger surface-protein interactions at low surface coverage and side-on—rather than vertical—antibody orientations (116,121). Gonzalez-Guerrero et al. (122) compared the performance of nanophotonic silicon nitride sensors functionalized with covalent silane chemistry and patterned with bioreceptors using both in-flow and droplet printing approaches. Similarly to our findings, the authors demonstrated improved analyte detection signal for the droplet printed sensors when using BSA/anti-BSA monoclonal antibody as a model receptor/analyte system. They attributed the improved performance of the droplet printed sensors to the improved reorganization capacity of the bioreceptor layer when allowed to react with the surface under static conditions, yielding more available binding sites. In addition to these mechanisms for improved signal with droplet-based functionalization, we hypothesize that in-flow antibody delivery may also increase the risk of functionalization solution contamination. Small volumes of fluid (e.g., blocking solution) captured in the fluidic delivery system’s rotary valve during reagent switching (e.g., 1.7 µL carryover fluid volume within the valve in our system (123)) may cause contamination and inhibit antibody immobilization. Protein that may have adsorbed to the walls of the fluidic system tubing and components and been incompletely removed by system cleaning protocols could also lead to functionalization solution contamination. Overall, the increased complexity of in-flow antibody immobilization compared to spotting-based immobilization appears to increase the risk of unsuccessful surface functionalization, which is illustrated in the representative peak shift trace shown in Fig. 6(c). In this PDA/flow trial, peak shifts were largely absent from the antibody immobilization and analyte detection stages in fluidic channel 1, suggesting unsuccessful surface functionalization despite confirmation of delivery of the same (split) functionalization solutions delivered to channel 2 (via monitoring of reservoir depletion and microfluidic outlet flow).

Final drift rates were negative for all sensors, regardless of functionalization approach and analyte concentration. These negative slopes indicate the removal of material from the sensor surface (21) and can be largely attributed to dissociation of capture antibody-spike protein complexes as the sensor is rinsed with analyte-free buffer. Among the assays employing covalent functionalization, the average final drift rate for PDA/spotting assays (−1.3 ± 0.1 pm/min) was significantly greater in magnitude (p = 4.9 ×10^-8^) than that measured in the PDA/flow assays (−0.54 ± 0.22 pm/min). This is likely correlated with the greater amount of analyte binding in the PDA/spotting assays. Assuming 1:1 Langmuir binding kinetics between the surface-immobilized capture antibodies and spike protein, the rate of dissociation of the antibody-antigen complexes is proportional to the concentration of surface-bound spike protein at equilibrium (124–126). Similarly, the Protein A/flow assays involving 20 µg/mL spike protein detection, which exhibited the greatest analyte binding signals due to the high spike protein concentration, also exhibited significantly greater final drift rates at −3.5 ± 0.6 pm/min. The Protein A/flow assay involving 1 µg/mL spike protein detection exhibited a greater final drift rate (−1.9 pm/min) than the other assays involving 1 µg/mL spike protein detection, even though this assay exhibited a lower analyte-specific signal than the PDA/spotting assays. This may be indicative of other contributors to the drift rate, such as Protein A and capture antibody removal from the sensor surface due to the relatively weak non-covalent interactions used to immobilize these proteins (27). Meanwhile, PDA films have been found to exhibit good stability in PBS and antibodies covalently attached to the PDA-coated sensors are not expected to desorb from the surface (46,110). For all functionalization approaches, desorption of BSA molecules introduced during the BSA challenge may contribute to drift rates.

The inter-assay CV for spike protein detection using PDA/spotting functionalization was 18.1%, which is lower than the ∼20% CV threshold typically used for immunoassay validation (12,93,94). Nevertheless, this inter-assay variation is larger than the single-channel intra-assay variability (CVs = 5.4 ± 1.9% and 2.8 ± 0.7%) and channel-level intra-assay variability (CVs = 4.7 ± 0.6%), highlighting that future efforts to improve replicability in this critical analyte detection stage should target factors that contribute to assay-level, rather than sensor- and channel-level variability. Indeed, tight process controls should be practiced in reagent storage/handling, reagent preparation, and functionalization procedures to the extent possible. We have included in ESI Table S19 a comparison of the sensor-level, channel-level, and assay-level CVs for each of the performance metrics and functionalization strategies discussed here. Examining these data shows that the PDA/spotting approach appears to reduce the channel-level intra-assay variability by ∼57% compared to the 20 µg/mL Protein A/flow strategy. This suggests that the spotting-based functionalization strategy may effectively address some of the channel-level factors that dominate variability in the flow-based approach (e.g., functionalization reagent depletion or contamination due to the tubing upstream of the chip). However, the inter-assay CV increased from 11.1% to 18.1%. Although some of this increase may be related to the ∼3× smaller analyte-binding signals (due to detection of spike protein at a concentration of 1 µg/mL rather than 20 µg/mL), this increase in the inter-assay CV suggests that the PDA/spotting approach (in which functionalization is performed at the chip-level rather than the channel-level) may introduce additional factors that vary across assays. For example, there may be variability in how and for how long the chips were dried after functionalization and prior to fluidic integration that contributed to variability in the activity of the immobilized antibodies. Future work could explore the use of immunoassay stabilizers (as previously used for SiP biosensor functionalization protocols (55)) to reduce this contributor to assay-level variability. In addition, further replicability improvements could be made by increasing the analyte detection signal by way of amplification strategies, such as enzymatic or particle-based amplification (27). Such signal enhancement would increase the mean detection signal, making the assay more robust to noise and day-to-day variations, while also allowing for detection of lower analyte concentrations. These potential benefits should, however, be weighed with tradeoffs associated with more complicated and time-consuming assay protocols and the introduction of additional reagents.

Taken together, these results highlight that PDA-mediated antibody immobilization offers comparable or improved analyte detection performance compared to Protein A-mediated functionalization, with the added benefit of sensor regenerability. As PDA can be deposited on many types of surfaces, PDA-mediated functionalization approaches are applicable to numerous classes of biosensors, including electrochemical sensors, other optical sensors (e.g., SPR, SERS), piezoelectric sensors, and lateral flow assays (109,127–129). Spotting-based antibody immobilization on PDA-functionalized sensors is a simple protocol that allows for significantly improved analyte detection signal compared to flow-based application and is amenable to multiplexing (e.g., via high-throughput precision spotting techniques such as piezoelectric inkjet and pin printing). Toward developing reliable point-of-care multiplexed diagnostics, future work should focus on minimizing inter-assay variability and combining the PDA/spotting functionalization approach reported herein with high-precision patterning tools. It should be noted, however, that these precise spotting techniques use nL-scale droplets that are much smaller than the 20 µL spots used in this work (27,130,131). Order-of-magnitude mass transport calculations (ESI S6.1) suggest that nL-volume spots with diameters of 150 µm still enable the transport of 10× excess antibodies to fully saturate the sensor surface in the spotting region within the first 1.5 minutes of incubation. However, controlling drying rates of these tiny spots to ensure sufficient time for receptor conjugation to the surface, minimizing droplet splashing upon impact with the surface, controlling spot morphology, and minimizing the coffee ring effect are among the factors introduced by precision spotting techniques that may increase variability in functionalization and analyte detection performance (55,64). In contrast to flow-based immobilization, offline spotting-based functionalization also does not enable real-time monitoring of sensor data throughout the functionalization process, which can provide helpful insight into whether or not bioreceptors have been successfully attached to the sensor surface prior to analyte detection.

The use of clinically relevant biological sample matrices, lower analyte concentrations, and more complicated assay formats (e.g., involving signal amplification) are expected to contribute to assay variability and increase the risk of false positive signals and should, therefore, be studied for assays employing PDA-functionalized SiP sensors. In ESI Section S9, we present preliminary data for assays to detect a different target analyte with different physicochemical properties, aimed at beginning to study these issues for the PDA/spotting functionalization approach. We demonstrate the detection of Interleukin 8 (IL-8) on PDA-functionalized SiP sensors in both simple (PBS-BSA) and complex (complete cell culture medium containing 10% fetal bovine serum (CCM)) sample matrices using a sandwich assay format with streptavidin-HRP (SA-HRP)-based signal amplification. Surface blocking with BSA alone and in combination with CCM are compared. We also demonstrate the use of capture antibody-free reference sensors to serve as a control for nonspecific binding. In both simple and complex media, we observe significantly larger amplification signals for samples containing 3.125 ng/mL IL-8 compared to a negative control (0 ng/mL IL-8), validating specific detection of IL-8 in complex media. We find that blocking with BSA combined with CCM significantly reduces nonspecific binding of matrix components in the IL-8/CCM sample compared to BSA blocking alone, potentially due to the greater molecular diversity of the BSA + CCM blocking solution, which may improve saturation of unoccupied sites on the sensor surface (117). However, this reduction in nonspecific binding is not sufficient to permit direct label-free detection of 3.125 ng/mL IL-8 in CCM compared to the control, requiring the sandwich assay format for detection in complex media. We also observe a higher inter-assay CV for the amplification stage of this IL-8 detection assay (e.g., 27% inter-assay CV for detection of IL-8 in simple buffer) compared to the direct label-free detection of 1 μg/mL spike protein reported in Fig.8. The ∼320× lower concentration used in this assay likely contributes to this higher variability. The more complex assay format may also contribute to this higher variability (with SA-HRP binding shifts dependent not only on the capture antibody-IL-8 binding interaction but also on the detection antibody-IL-8 interaction and the biotin-streptavidin interaction). Further studies should explore a rigorous analysis of factors contributing to variability and strategies for improving replicability in such assays performed in biological media using low analyte concentrations and signal amplification. The optimization of antifouling treatments including protein-(e.g., serum proteins, casein), polymer-(e.g., oligoethylene glycol/polyethylene glycol layers, zwitterionic layers), and peptide-based blockers for PDA-functionalized sensors should also be rigorously evaluated to facilitate accurate and replicable analyte detection in complex biological media (65,132).

## 4. Conclusions

We have characterized and compared, for the first time, a set of both intrinsic and analyte-detection performance metrics and their replicability at the sensor-, channel-, and assay-levels using SiP SWG MRR biosensors (summarized in ESI Table S19). We measured comparable CVs for S_bulk_ using our SWG-based ring resonator sensors compared to those previously reported in the literature, despite the smaller fabrication feature sizes used for SWG waveguides. These data support the use of SWG waveguides for biosensors, leveraging their strengths of high sensitivity and tunable optical properties. We also found that CVs for the metrics incorporating stability were considerably higher than those for sensitivity, perhaps due to the stochastic nature of the measured noise. Sensor instability due to bubbles in the microfluidic system had a critical effect on the sensor performance, illustrating the importance of mitigating bubbles in microfluidics-integrated SiP biosensor systems. We found that effective bubble mitigation in PDMS microfluidics integrated with SiP biosensors can be achieved by strategies such as pre-wetting fluidic channels with Triton X-100 surfactant solution and degassing and plasma treating PDMS gaskets prior to use. Through SARS-CoV-2 spike protein demonstration assays (1 µg/mL), we showed that functionalization via covalent PDA/spotting-mediated antibody immobilization improves analyte detection signal by 8.2×and 5.8×compared to PDA/flow and noncovalent Protein A/flow approaches, respectively. When comparing the replicability of intrinsic (S_bulk_) and analyte-detection (Δλ_binding_) performance, the intrinsic performance (S_bulk_) showed higher replicability, consistent with the lower number of factors contributing to variability of this metric (Table 2). We found that for analyte-detection replicability when using Protein A/flow functionalization, factors unique to each fluidic channel (e.g., factors related to the pre-wetting process or the cleanliness and performance of the microfluidic system that may result in reagent depletion, dilution, or contamination) appear to dominate the variability in our system. In contrast, with PDA/spotting functionalization it appears that we were able to address some of the channel-level contributors, but factors unique to each assay (e.g., stability of the functionalized surface through the drying and rehydration phases) dominated the variability in our system, highlighting a path to future optimization. Future efforts should focus on reducing inter-assay variability for biosensors using PDA-based functionalization, improving surface blocking, developing robust surface regeneration protocols, and exploring signal amplification strategies.

Overall, this work describes a practical example of a framework to analyze and improve replicability that is applicable to many classes and configurations of biosensors: deconstructing the system and identifying factors that contribute to performance and replicability at each stage; identifying a set of performance metrics that can isolate different contributing factors; and characterizing the performance metrics and their replicability at different levels (intra-channel, inter-channel, inter-assay). By detailing the different performance metrics and different representations of variability for our system, we have provided a demonstration of this framework and proposed a structure by which SiP biosensor systems—broadly composed of the sensors, fluidic integration, and binding assay—can be analyzed, characterized, compared, and optimized. We have then used this structure with a representative system of SWG MRR sensors and demonstration binding assays detecting the SARS-CoV-2 spike protein to better understand some of the important factors limiting system performance. We anticipate that with future optimization and isolation of different components, this structure will serve as a valuable, methodical tool for silicon photonics researchers and biosensors researchers more broadly to approach the challenging and interdisciplinary task of biosensor development.

## Supporting information

ESI

## Author contributions

**Conceptualization**, L.S.P. and S.M.G.; **Data curation**, L.S.P., S.M.G., K.N., S.J.K., S.K., S.C., M. Wang, and M. Wei; **Formal analysis**, L.S.P., S.M.G., K.N., and M. Wang; **Funding acquisition**, S.M.G., L.C., S.S., and K.C.C.; **Investigation**, L.S.P., S.M.G., K.N., S.J.K., S.K., S.C., M. Wang, M. Wei, Y.O.Y., and Y.H.; **Methodology**, L.S.P., S.M.G., K.N., S.J.K., S.K., S.C., M. Wang, M. Wei, Y.O.Y., A.R., and Y.H.; **Project administration**, L.S.P., S.M.G., L.C., S.S., and K.C.C.; **Software**, S.M.G. and A.R.; **Resources**, S.M.G., L.C., S.S., and K.C.C.; **Supervision**, S.M.G., L.C., S.S., and K.C.C.; **Validation**, L.S.P., S.M.G., and K.N.; **Visualization**, L.S.P., S.M.G., K.N., and M. Wang; **Writing - original draft**, L.S.P., S.M.G., and K.N.; **Writing - review & editing**, L.S.P., S.M.G., K.N., S.J.K., S.K., S.C., M. Wang, M. Wei, L.C., S.S., and K.C.C.

## Data availability

Data supporting this article have been included as part of the Electronic Supplementary Information.

## Acknowledgements

The acknowledgements come at the end of an article after the conclusions and before the notes and references.

