## Supplementary material for "Resonating with replicability: factors shaping assay yield and variability in microfluidics-integrated silicon photonic biosensors": ESI

#### Table of Contents

S1. Intra- and inter-assay CVs reported for SiP sensors 2
S2. Supplementary experimental details 3
- S2.1. Chemicals and reagents 3
- S2.2. Sensor chip design and fabrication 3
- S2.3. Microfluidic gasket design, fabrication, and assembly 3
- S2.4. Photonic testing setup 4
- S2.5. Fluidic testing setup 5
- S2.6. Fluidic priming, pre-wetting, and risk mitigation 6
  - S2.6.1. Priming 6
  - S2.6.2. Pre-wetting 6
  - S2.6.3. Residues due to salt precipitation and fluidic components when ethanol is used for pre-wetting 7
  - S2.6.4. Minimizing contamination during fluidic protocols 8
  - S2.6.5. Pressure-driven flow control system maintenance 8
- S2.7. Contact angle measurements 8
- S2.8. Characterizing bubbles in microfluidic devices 9
- S2.9. Polydopamine film thickness 9
- S2.10. Bulk refractive index sensitivity testing 11
  - S2.10.1. Intrinsic sensor performance experiments and demonstration assays (20 µg/mL spike protein) 11
  - S2.10.2. Demonstration assays (1 µg/mL spike protein) 13
- S2.11. Reservoir setups and fluidic protocols for demonstration assays 13
- S2.12. Sensing data analysis workflow 15
  - S2.12.1. Resonance peak tracking 15
  - S2.12.2. Bulk refractive index sensitivity extraction 15
  - S2.12.3. Baseline correction 15
  - S2.12.4. Peak shifts and slopes quantification 16

S3. Statistical analysis 16
- S3.1. Bubble mitigation strategy comparisons 16
- S3.2. Demonstration assay comparisons 18

S4. Sensor instability due to bubble exposure 18
S5. Sensor signal dependence on channel position 20
S6. Mass transport estimates 20
S6.1. Spotting-mediated antibody immobilization 21
S6.2. Flow-mediated antibody immobilization 21
S6.3. In-flow analyte detection 22
S6.4. Limitations 22
S7. Summary of sensor-level, channel-level, and assay-level CVs 22
S8. Baseline-correction fit functions 25
S9. Demonstration of a sandwich immunoassay for IL-8 detection in complex media on polydopamine-functionalized SiP sensors 27
S9.1. Methods 27
S9.1.1. IL-8 detection assay methods 27
S9.1.2. Assays incorporating reference sensors 29
S9.2. Performance comparison for IL-8 detection in simple and complex media 30
S9.3. Statistical testing 33
References 33

### S1. Intra- and inter-assay CVs reported for SiP sensors

**Table S1.** Intra- and inter-assay coefficients of variation (CVs) reported in the literature for silicon photonic (SiP) sensors used for bulk refractive index (RI) and biological analyte detection.

| Sensor type | Analyte | Metric | Intra-assay CV (%) | Inter-assay CV (%) | Refs. |
|---|---|---|---|---|---|
| SiP MZI | Cardiac troponin | Analyte detection signal (phase shift) | – | 10 | (1) |
| $Si_3N_4$ MRR | Neutravidin | Analyte detection signal (resonance shift), 10 µg/mL | 13.9 | 11.0 | (2) |
| | | Analyte detection signal (resonance shift), 50 µg/mL | 10.0 | 13.9 | |
| Si MRR | NaCl | Bulk RI detection signal (resonance shift), 0.5 M | 0.5–1.8 (ring-to-ring)<br>0.3–1.7 (between 4-ring clusters) | 0.4–1.8 (chip-to-chip) | (3) |
| | | Bulk RI detection signal (resonance shift), 2.0 M | 0.4–1.1 (ring-to-ring)<br>0.2–1.1 (between 4-ring clusters) | 0.5–1.8 (chip-to-chip) | |
| | Anti-SS-A antibodies in diluted serum | Analyte detection signal | 0.6-6.4 (ring-to-ring)<br>0.3–6.1 (between 4-ring clusters) | 2.3–3.7 (chip-to-chip within an array$^{\dagger}$)<br>5.0 (total) | |
| Si MRR | Salt | Bulk RI detection signal (resonance shift), 0.15 M | 3.9 (ring-to-ring)<br>0.47 (cycle-to-cycle) | – | (4) |
| $Si_3N_4/SiO_2$ TriPleX MRR | Neutravidin (adsorption) & BSA-biotin (capture) | Analyte detection signal (resonance shift) | 0.3–2 (ring-to-ring) | – | (5) |
| | Neutravidin (covalent immobilization) | | 1.6 (ring-to-ring) | – | |
| | Biotinylated thrombin-binding aptamer (immobilization) & thrombin (capture) | | 2.8 (ring-to-ring) | – | |
| $Si_3N_4$ MZI | Monoclonal antibody against SARS-CoV-2 spike protein receptor binding domain in diluted serum | Analyte detection signal (phase shift) | 3.6–5.7 (between triplicate measurements of 3 samples performed on the same day) | 6.9–8.6 (across duplicate measurements of 3 samples measured on 6 different days) | (6) |
| Nanophotonic BiMW | microRNA-21 | Calibration fit parameters | – | 5.3–5.7 (in buffer)<br>2.6–3.4 (in plasma) | (7) |

| | | | | |
|---|---|---|---|---|
| | | Limit of detection | – | 51% (in buffer)<br>17% (in plasma) |

MZI: Mach-Zehnder interferometer; MRR: microring resonator; BiMW: bimodal waveguide interferometer.

†An array refers to a group of 12 sensor chips that were loaded into the same readout instrument. Each chip was addressed by two microfluidic channels. One array of 12 chips could be read out within ~2 hours (same day). For this immunoassay, 6 replicate arrays of 12 chips were tested.

### S2. Supplementary experimental details

#### S2.1. Chemicals and reagents

Ultrapure water (ASTM Type I) was acquired from a NANOpure® Diamond water purification system (Barnstead™, Thermo Fisher Scientific, Waltham, MA, USA). Compressed nitrogen gas (99.995% purity, NI 4.8-T) was purchased from Linde Canada (Mississauga, CAN). Phosphate buffered saline (PBS, Cat: 10010-023, pH 7.4, 1X), Tris buffer (0.5 M, pH 8.6, Cat: AAJ62287AP), NaCl (Cat: S271-3), and glycerol (Cat: G33-4) were purchased from Thermo Fisher Scientific (Waltham, MA, USA). Bovine serum albumin (BSA, Cat: A7906), Protein A from *Staphylococcus aureus* (Cat: P6031, Lot: 0000142821), dopamine hydrochloride (Cat: H8502), 37% HCl concentrate (Ca: 258148-2.5L-GL), glycine (Cat: G8790-100G), and Triton X-100 (Cat: T8787-100ML) were purchased from Sigma-Aldrich (St. Louis, MO, USA). Ethanol (Cat: P016EAAN) was purchased from Greenfield Global (Mississauga, CAN). Spike protein (SARS-CoV-2 (2019-nCoV) Spike S1-His Recombinant Protein, Cat: 40591-V08H, Lot: LC16AP1501) and spike protein capture antibody (SARS-CoV-2 (2019-nCoV) Spike S1 Antibody, Rabbit MAb, Cat: 40150-R007, Lot: MA14MY2901-A) were purchased from Sino Biological, Inc. (Beijing, China). Glycine-HCl buffer was prepared by dissolving 10 mM glycine and 160 mM NaCl in ultrapure water, then adjusting to pH 2.2 using HCl. PBS-BSA running buffer was prepared by dissolving 0.1 mg/mL BSA in PBS.

#### S2.2. Sensor chip design and fabrication

The SWG MRR photonic circuits were designed using KLayout mask editing software, the open-source SiEPIC tools library, SiEPIC EBeam process design kit, and the Applied Nanotools (ANT) process design kit (8–10). 500 nm-wide strip routing waveguides were used to transmit C-band light between the I/Os and resonators. Routing waveguide bends were designed with a bend radius of 5.0 µm and a Bezier bend parameter of 0.2 (11). 10 µm-long tapers were used to create smooth transitions between the routing waveguides and the SWG bus regions of the resonators. The chips were fabricated with a 220 nm silicon device layer, comprising the patterned photonic circuit, on top of a 2.0 µm $SiO_2$ buried oxide layer, which in turn sat atop a 725 µm silicon handle wafer. For this work, the chips were fabricated without cladding. No photoresist or hard mask remained on the waveguide surfaces after fabrication. The chips were stored in Gel-Pak® containers and used as received for testing. Extinction ratios (ER), quality factors (Q), and free spectral ranges (FSR) were quantified for the fabricated sensors using methods described previously (12).

**Table S2.** Resonator performance metrics for SWG MRRs measured in different cladding liquids and at different stages of surface functionalization. All MRRs included in this analysis had a coupling gap of $g_c$ = 500 nm and duty cycle of $\delta$ = 0.7.

| Resonance spectrum acquisition conditions | | Number of replicate chips included in analysis | Number of replicate MRRs analyzed per chip | ER (inter-chip mean ± standard deviation), dB | Q (inter-chip mean ± standard deviation) | FSR (inter-chip mean ± standard deviation), nm |
|---|---|---|---|---|---|---|
| Cladding liquid | Sensor chip condition | | | | | |
| Water | Bare chip | 8 | 8 | 22.3 ± 3.7 | $8.39 \times 10^3 \pm 2.13 \times 10^3$ | 4.24 ± 0.08 |
| PBS | Bare chip | 4 | 8 | 19.3 ± 2.9 | $6.65 \times 10^3 \pm 890$ | 4.16 ± 0.28 |
| PBS | PDA-coated | 3 | 8 | 17.8 ± 3.1 | $7.07 \times 10^3 \pm 182$ | 4.21 ± 0.01 |
| PBS-BSA | PDA-coated, spotted with capture antibody, blocked with BSA | 3 | 8 | 17.5 ± 4.0 | $8.23 \times 10^3 \pm 1.36 \times 10^3$ | 4.10 ± 0.14 |

#### S2.3. Microfluidic gasket design, fabrication, and assembly

The microfluidic gasket design (Fig. S1(a)) includes one straight channel, while the other channel features a bent configuration, where the inlet and outlet are connected by a 45-degree segment. The gasket includes 3 mm wide oval through hole features that can accommodate 4-40 threaded rods at a range of y-positions to facilitate alignment to the photonic chip. All procedures regarding mould design, printing, and post-processing as well as PDMS preparation, casting, curing, demoulding, and postprocessing have been previously described by our team (12).

To assemble the photonic chip-microfluidic gasket assembly for fluidic testing (Fig. S1(b–c)), the SiP chip was placed in the machined recess of an aluminium mounting plate. 4–40 threaded rods (McMaster-Carr, Elmhurst, IL, USA) were screwed into threaded holes of the mounting plate on either side of the chip. A rectangular washer of the same x-y dimensions as the fluidic gasket and with 4.5 × 2.5 mm rectangular holes aligned with the fluidic I/Os was custom laser-cut from ⅛” acrylic (McMaster-Carr, Elmhurst, IL, USA) using a Universal Laser Systems VersaLaser VLS2.30 laser cutter (Universal Laser Systems, Inc., Scottsdale, AZ, USA). The acrylic washer and PDMS gasket were manually aligned together with the aid of a 3D-printed jig, then aligned to the threaded rods in the mounting plate. The gasket and washer were lowered onto the photonic chip while visually inspecting the alignment of the gasket’s fluidic channels to the photonic chip’s MRRs using an optical microscope (Aven MicroVue Digital Microscope, Aven Tools, Ann Arbor, MI, USA). Nuts were tightened onto the threaded rods to seal the fluidics against the photonic chip. The washer served to provide even pressure to the flat PDMS gasket to maintain a good seal between the PDMS and the photonic chip.

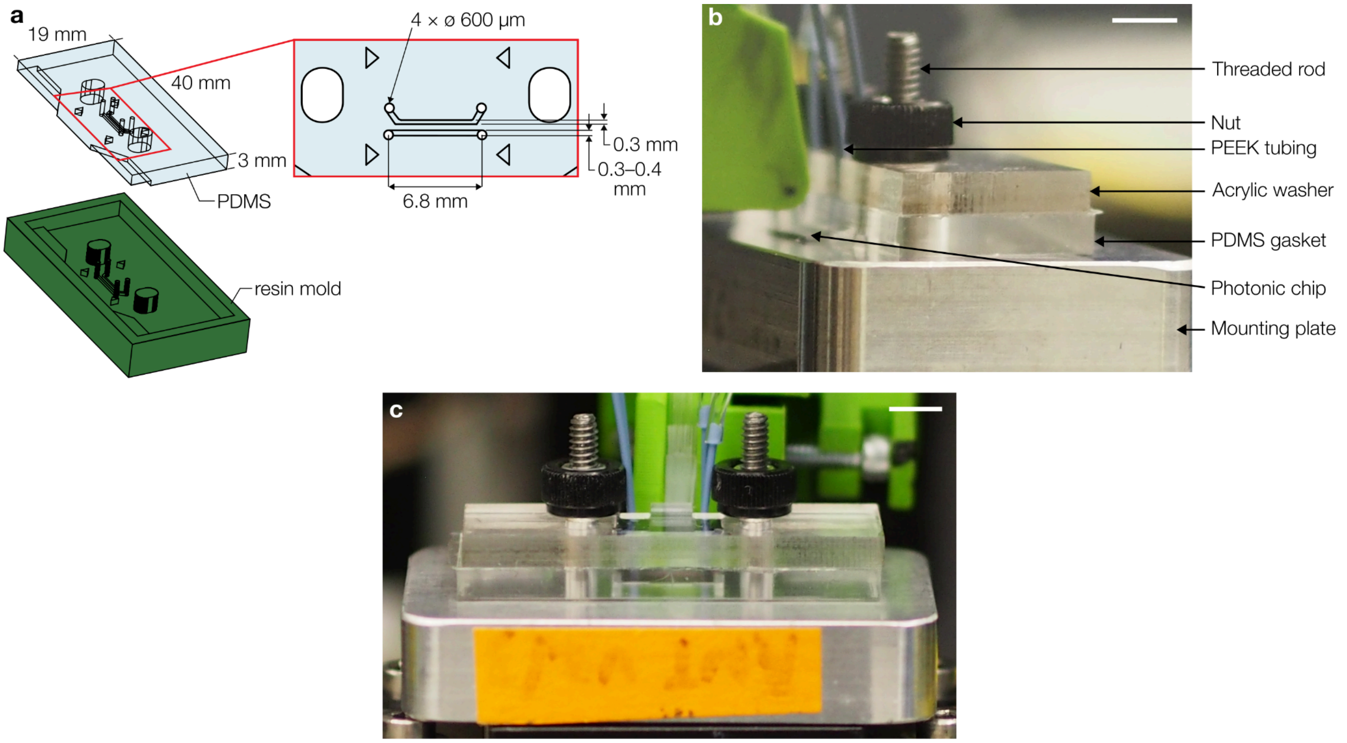

**Fig. S1.** Microfluidic gasket design and photonic chip-microfluidic gasket assembly for fluidic testing. **(a)** Illustrations of microfluidic gasket design. **(b-c)** Images of the photonic chip mounted to the assembly and coupled with the fibre array.

#### S2.4. Photonic testing setup

The chip-gasket assembly was secured on the motorized and temperature-controlled XY stage of the testing setup using thermally conductive tape. The fiber array was mounted to a motorized Z stage to enable alignment to the on-chip grating coupler inputs and outputs. The fiber array channel aligned to the chip’s input grating coupler was connected to the C-band swept tunable laser. Eight fiber array channels aligned to the output grating couplers were connected to the optical detectors. Alignment of the fiber array to the photonic chip was performed using open-source PyOptomip software (Python 2.7, 32-bit) (13). During sensing experiments, the fiber array alignment was monitored by a custom Python acquisition software GUI and adjusted every 30 sweeps using a fine align function to ensure good coupling to the on-chip grating couplers throughout each experiment. All optical and data connections between the photonic chip and external equipment are illustrated in Fig. S2.

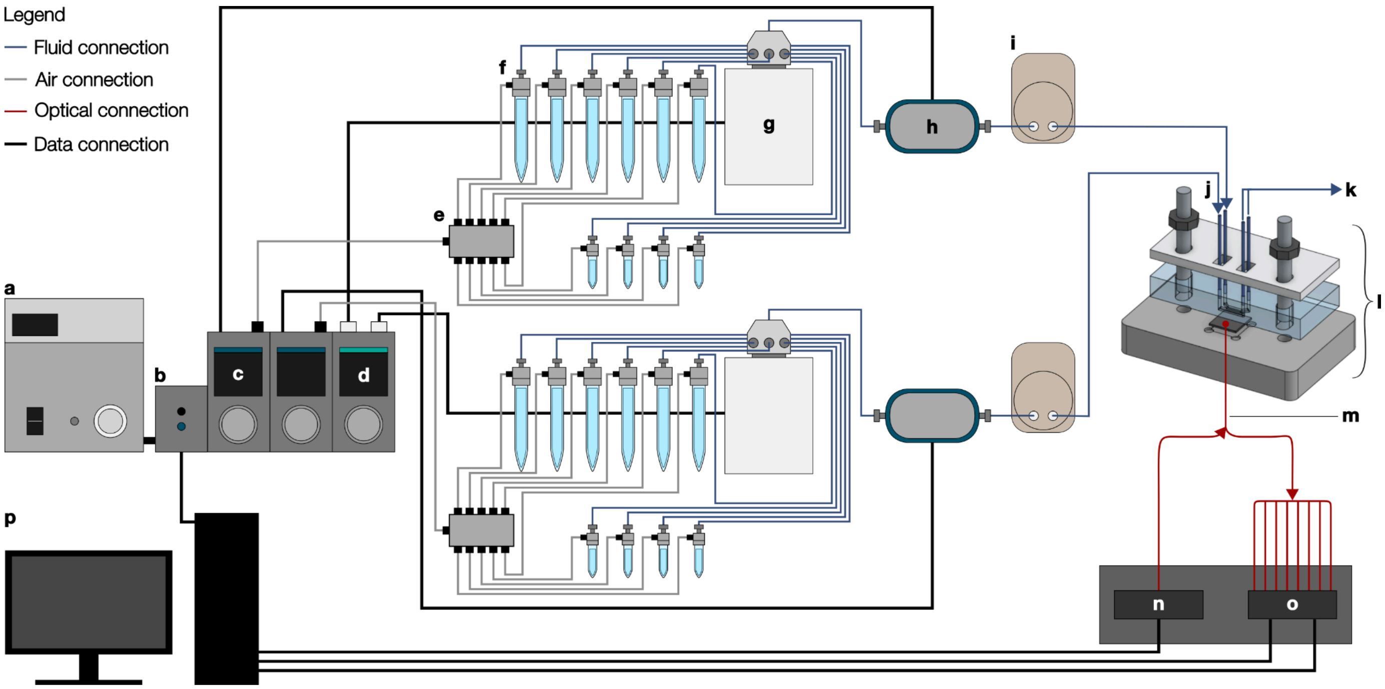

**Fig. S2.** Photonic-microfluidic testing setup. Illustration containing all fluidic and optical modules and interfaces with the PC. The fluidic control and delivery system includes two individually addressable fluidic channels and consists of the following components: **(a)** pressure source, **(b)** software control module, **(c)** pressure-based flow controller, **(d)** microfluidic valve controller, **(e)** 10-position air manifold, **(f)** fluidic reservoirs, **(g)** 10-position bidirectional valves, **(h)** bidirectional flow rate sensor, and **(i)** bubble trap. The **(l)** chip-gasket assembly is connected to the fluidic delivery system via **(j)** fluidic inlets and **(k)** fluidic outlets. The sensor chip is interfaced with an **(m)** optical fibre array that is connected to a **(n)** C-band tunable laser and **(o)** photodetectors. Overall system control is achieved via a **(p)** PC.

#### S2.5. Fluidic testing setup

The Fluigent fluidic control system (Fig. S2) consists of a FLPG Plus pressure source and LineUp™ modules including the LINK (LU-LNK-002) software control module, two Flow EZ™ (LU-FEZ) pressure-based flow controllers, two M-SWITCH™ (ESSMSW003) 10-position bidirectional valves, a SWITCH-EZ (ELUSEZ) microfluidic valve controller, and two FLOW UNIT (model M) bidirectional flow rate sensors. In each of the two fluidic channels, reagents were stored in six 15-mL Falcon® tube (Corning Inc., Corning, NY, USA) and four 2-mL Eppendorf Safe-Lock tube (Eppendorf, Hamburg, Germany) reservoirs equipped with Fluigent P-CAP airtight metal caps. The reservoirs were connected to the air pressure output of each flow controller via 4 mm OD tubing and a 10-position air manifold (Fluigent) to facilitate pressurization, and to the fluidic inlets of the bidirectional valves via 1/16” OD, 0.010” ID FEP capillary tubing (IDEX 1527L, Cole-Parmer Canada, Quebec, QC, CAN). The outlets of the two bidirectional valves were connected to the inlets of the two flow sensors using the same FEP tubing. The flow sensor outlets were each connected to bubble traps (PG-BT-REC25UL, PreciGenome, San Jose, CA, USA) via two lengths of PEEK 1/32” OD, 0.010” ID capillary tubing (IDEX 1531B, Cole-Parmer Canada, Quebec, QC, CAN), joined by a ~10 cm portion of Tygon® 0.02” ID, 0.06” OD microbore tubing (Masterflex® Microbore Transfer Tubing, Tygon® ND-100-80, Cole-Parmer Canada, Quebec, QC, CAN). The bubble trap outlets were connected to PEEK capillary tubing. In experiments using aqueous solutions for sensor pre-wetting, the ends of the PEEK capillary tubing (IDEX 1531B, Cole-Parmer Canada, Quebec, QC, CAN) connected to the bubble trap outlets were directly inserted into the inlet ports of the microfluidic gasket after priming to supply fluid to the photonic chip assembly. In experiments using ethanol-based pre-wetting, an additional pre-wetting assembly was included in the path for each microfluidic channel

For the pre-wetting assembly, a conical adapter (IDEX P-797, Cole-Parmer Canada, Quebec, QC, CAN) with FEP tubing sleeve (IDEX F-247, Cole-Parmer Canada, Quebec, QC, CAN) was attached to the ends of the PEEK tubing connected to each bubble trap. To deliver fluid from the Fluigent system to the sensor chip, the conical adaptors were each connected to a ~5 cm length of ultra chemical-resistant Tygon® 1/16” ID, 1/8” OD tubing (Tygon® 2375, Saint-Gobain Life Sciences, Solon, OH, USA), which was connected to another length of PEEK tubing via a second conical adapter and FEP tubing sleeve. This PEEK tubing was inserted into the inlet ports of the microfluidic gasket to supply fluid to the photonic chip assembly. During pre-wetting, the short lengths of 1/16” ID Tygon® tubing were disconnected from the bubble traps and, instead, each connected to a 5 mL Luer-Lok™ syringe (BD-Canada, Mississauga, ON, CAN) via a threaded Luer adapter (IDEX P-659, Cole-Parmer Canada, Quebec, QC, CAN), fingertight fitting (IDEX F-300, Cole-Parmer Canada, Quebec, QC, CAN), the same 1/16” OD, 0.010” ID FEP tubing used in the Fluigent system, and conical adapter. This facilitated delivery of ethanol from the syringe to the photonic chip assembly while avoiding contact between ethanol and fluidic components incompatible with this solvent.

#### S2.6. Fluidic priming, pre-wetting, and risk mitigation

##### S2.6.1. Priming

For all experiments, fluidic reservoirs were primed using an automated protocol that delivered 750 mbar constant pressure for 2 minutes to drive flow from each reservoir. This priming protocol was previously validated by our team to deliver ~5–10× excess of the volume required to fill the internal volume of the system between the reservoir and the flow sensor (downstream of the rotary valve) and the system was monitored (by inspecting the logged flow sensor data) to ensure that (1) the full air plug passed through the flow sensor before the end of the priming period for each reservoir and (2) the system was able to reach the expected flow rates of water (greater than the 120 μL/min maximum range of our flow sensor) for each reservoir, to ensure that none of the lines had full or partial blockages.

##### S2.6.2. Pre-wetting

For the experiments that used ethanol pre-wetting (bubble monitoring experiments testing ethanol pre-wetting and assays that detected 20 μg/mL spike protein using the Protein A and flow-mediated functionalization process), an ethanol pre-wetting process was designed to mitigate residue deposition on microfluidic sensors as well as accidental bubble introduction during solution changes (ESI S2.6.3). First, the primed Fluigent system was flushed with ultrapure water to remove any traces of salt that could precipitate in contact with residual ethanol in the gasket (750 mbar constant pressure until >5 mL water was collected). The pre-wetting assembly components were dried by flowing filtered compressed air through all components for >20 mins, to ensure that ethanol—not residual water—made first contact with the microfluidic channels to wet small crevices. The pre-wetting assemblies were then connected such that the Luer-Lock adapter and the length of PEEK tubing used to couple to the gasket inlet were connected to each other through their conical adapters and the ultra chemical-resistant Tygon® tubing. The remaining conical adapter was connected to the Fluigent system. For each microfluidic channel, a 3 mL Luer-Lock syringe (Becton Dickinson 309657) was rinsed 3× with anhydrous ethanol (by filling, waiting ~30 s, and then emptying) to remove syringe lubrication oil, and subsequently refilled and used to slowly fill and then rinse the pre-wetting assembly with ethanol, tapping the conical adapters while flowing to help release any bubbles trapped inside. The syringe was then refilled with 3 mL anhydrous ethanol, reconnected to the pre-wetting assembly, and flowed until no bubbles remained in the assembly or syringe and >2.5 mL ethanol remained in the syringe. A second 3 mL Luer-Lock syringe was rinsed 3× with ultrapure water and refilled with ultrapure water such that no bubbles were trapped inside. The PEEK tubing outlets of the ethanol-filled pre-wetting assemblies were then connected to the microfluidic gasket and very gentle manual pressure on the syringe was used to slowly fill the microfluidic channels with ethanol. Approximately 1–2 mL of ethanol was delivered through each microfluidic channel before each outlet Tygon® tubing segment was blocked using a ratchet-style pinch-clamp to stop the flow. For each channel, the ethanol syringe was carefully disconnected from the Luer-Lock adapter while supplying gentle pressure to the syringe plunger (to avoid bubble introduction into the adapter) and the ultrapure water syringe was connected after flowing a small amount of water to make sure there was no bubble at the tip of the syringe, and a convex droplet of water was produced at the syringe tip. The outlet tubing was then unblocked and the microfluidic device was rinsed with ~2 mL ultrapure water to flush the ethanol out of the pre-wetting system and gasket. Next, the gasket was connected to the Fluigent system. The outlet tubing was blocked again and the conical adapter connected to the pre-wetting syringe was very slowly and carefully disconnected from the Tygon® 2375 tubing (again while supplying gentle pressure to the syringe plunger to avoid bubble introduction). The Fluigent system was set to provide a constant pressure of 50 mbar to drive slow ultrapure water flow, and the conical adapter at the Fluigent system outlet was connected to the gasket through the Tygon® 2375 tubing after verifying that there was a water droplet forming at the tip of the adapter due to the water flow. Finally, the outlet tubing was unblocked and the Fluigent system set to flow ultrapure water at a constant 30 μL/min flow rate.

For the experiments that used Triton X-100 pre-wetting of degassed and plasma-treated microfluidic gaskets (bubble monitoring experiments testing Triton X-100 pre-wetting and all assays that detected 1 μg/mL spike protein), the pre-wetting process was simpler, more controlled, and less error-prone. The Fluigent system was primed with 0.3 mM Triton X-100 in PBS pre-wetting solution after priming all other lines. It was then set to provide a slow flow (2–10 μL/min) of pre-wetting solution. Immediately after removing the microfluidic gasket from the vacuum storage, plasma-treating it, and assembling the photonic chip and gasket, dry outlet tubing was connected to the microfluidic outlets and blocked using ratchet-style pinch-clamps. The same pinch clamps were installed on the Tygon® tubing connecting the bubble trap to the flow sensor for each Fluigent channel. After setting the output pressure for each Fluigent channel to zero, the PEEK tubing outlets from each bubble trap were raised slightly above the reservoirs to introduce ~5 s of backflow (typically at a flow rate of 5–6 μL/min) and the bubble trap Tygon® tubing lengths were quickly blocked using the pinch clamps to stop the flow and PEEK tubing connected to the inlet for each channel. The small amount of backflow was intended to ensure that the pre-wetting solution entered the channel at a controlled flow rate, rather than during PEEK tubing connection to the gasket. Finally, the bubble trap tubing and outlet tubing were unblocked and Fluigent configured to deliver the pre-wetting solution at a constant flow rate of 1–2 µL/min. The channels were monitored with a time-lapse image acquisition while the channel was filled with fluid, and the flow rate was increased to 30 µL/min when the fluid meniscus was visible in the outlet Tygon® tubing. After ~5 min of pre-wetting solution flow, the Fluigent reservoir being delivered was switched to PBS (for flow-based functionalization) or running buffer (for spotting-based functionalization).

All Triton-X-100-containing priming and pre-wetting waste was collected separately and disposed of through our institutional hazardous waste disposal program, in accordance with hazardous waste disposal guidelines.

##### S2.6.3. Residues due to salt precipitation and fluidic components when ethanol is used for pre-wetting

Fig. S3 depicts conditions that we anticipate could lead to residue formation on SiP biosensors if non-optimized ethanol pre-wetting conditions are used. Fig. S3(a) depicts salt precipitation in one microfluidic channel (#2) of a chip-gasket assembly, into which 375 mM NaCl was introduced after pre-wetting the channel with ethanol. The channel was subsequently flushed with ultrapure water before removing the gasket and drying. The other channel (#1) was pre-wetted in the same manner but salt was never introduced (only ultrapure water flow followed pre-wetting). Obvious residue formed over the resonators that were exposed to the salt solution, and this residue was not removed by flushing with water.

Fig. S3(b) depicts residue formation on a bare silicon wafer after a droplet of 100% ethanol was dispensed from a 1 mL syringe through a short length of Tygon® ND-100-80 tubing (0.02” ID, 0.06” OD). This formula of Tygon tubing is not compatible with organic solvents, and through isolating and testing the other parts of the system used in this test (syringe, needle, ethanol liquid) the bulk of the residue appears to be contributed by the tubing. Obvious residue remained on the wafer after ethanol evaporation, suggesting that ethanol pre-wetting using this type of incompatible tubing could also contribute to residue formation on SiP sensors.

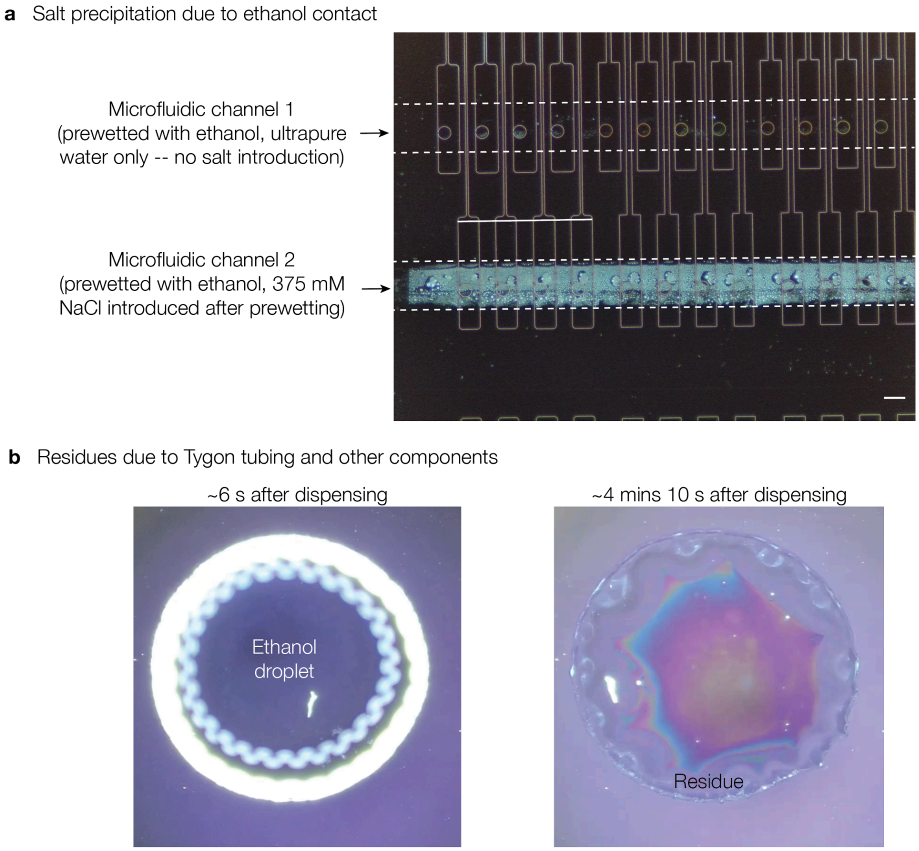

**Fig. S3.** Components of ethanol pre-wetting systems and protocols may lead to residue formation on SiP biosensors. **(a)** Micrograph depicting salt precipitation after pre-wetting in one microfluidic channel, when 0.375 mM NaCl was introduced after ethanol pre-wetting. Scale bar represents 100 µm, and dashed lines indicate approximate microfluidic channel positions. **(b)** Residue formation on a bare silicon wafer after a droplet of anhydrous ethanol evaporated from the surface. The ethanol was dispensed by a 1 mL syringe fitted with 22G blunt needle and short length of Tygon® ND-100-80 tubing (0.02” ID, 0.06” OD). By isolating the other components of the system (syringe, needle, ethanol), the dominant contributor to the residue formation appears to be the non-ethanol-compatible formulation of Tygon® tubing used in this test.

##### S2.6.4. Minimizing contamination during fluidic protocols

During reagent switches, the 10-position bidirectional rotary valves of the automated fluidic delivery system rotate to select the appropriate reservoirs for fluid delivery. As each valve rotates past primed lines connected to neighbouring reservoirs on its way to the desired reservoir, the 1.7 µL carryover fluid volume within the valve (14) may become contaminated with fluids from other reservoirs (which are pressurized throughout this process). To address this, reservoir layouts were carefully designed and valve rotation directions were defined in the automated fluidic protocols to minimize the number of reservoir lines crossed during each switch and to ensure that any contamination that might occur during these switches was tolerable. For example, reservoir layouts and valve rotation directions were selected to ensure that the valves did not pass fluidic lines containing ultrapure water or NaCl solutions during the protein-sensing portions of SARS-CoV-2 spike protein demonstration assays, minimizing the risk of protein denaturation. Additionally, in assays employing PDA/flow-mediated functionalization, the reservoir layout and fluidic protocol was designed to ensure that the valves did not pass fluidic lines containing BSA or other proteins prior to antibody immobilization. This was done to ensure that antibody immobilization was not inhibited by unintentional surface blocking prior to antibody delivery to the surface.

##### S2.6.5. Pressure-driven flow control system maintenance

Having encountered issues with inaccurate flow rate measurements and blockages in our automated pressure-driven fluidic control system, our laboratory has implemented strict protocols for working with this system to reduce the likelihood of issues that can affect assay yield. To reduce the likelihood of blockages within the small-bore tubing and components of the fluidic control system, all liquids supplied using the automated system are filtered with maximum pore size 70 µm using cell strainers (e.g., Falcon™ C352340, Thermo Fisher Scientific Canada), syringe filters (e.g., MilliporeSigma™ SLGPR33RS, Thermo Fisher Scientific Canada), or centrifugal filters (e.g., Corning™ 8160, Thermo Fisher Scientific Canada) as they are dispensed into the reservoirs. Furthermore, when using the automated system to deliver reagents containing proteins and organic materials, residues can build up on the internal surfaces of the flow rate sensors, leading to inaccurate flow rate measurements and regulation. These residues can also build up over time even when only using the system to deliver simple solutions like ultrapure water, isopropanol, and aqueous salt solutions. Residues can also contribute to blockages within the system, disrupting flow. These residues can be cleaned and further growth can be minimized by regularly cleaning the system with detergents and performing solvent flushes prior to shut-down (15). As such, our group performs a ramp-down procedure after each use of the system that involves detergent and solvent-based cleaning of the fluidic system. To perform ramp-down, we first disconnect the photonic chip-microfluidic gasket assembly from the fluidic path, then (1) flush all fluidic lines of both channels with ultrapure water at 750 mbar for 3 min. Next, (2) all fluidic lines that were used to deliver protein-containing reagents are flushed with Liquinox® Critical Cleaning Liquid Detergent (1% v/v in ultrapure water) (Alconox 1232-1, Cole-Parmer Canada, Quebec, QC, CAN) at 750 mbar for 3 min, then (3) flushed again with ultrapure water at 750 mbar for 3 min. Next, (4) the flow sensors are thoroughly cleaned by flowing ~7 mL of RBS 35 detergent (10% v/v in ultrapure water) (Cat: 27950, Lot: WB3191271, Thermo Fisher Scientific, Waltham, MA, USA) through both channels (e.g., at 120 µL/min for 60 min), followed by (5) rinsing with ~7 mL of ultrapure water. Next, (6) the bubble traps are disconnected from the fluidic path and (7) all fluidic lines are flushed with isopropanol (Cat: 270725-1L, Sigma-Aldrich, St. Louis, MO, USA) at 750 mbar for 3 min, prior to (8) flushing with air at 750 mbar for 3 min.

#### S2.7. Contact angle measurements

Contact angle measurements were performed using a custom imaging setup in which the sample surface was positioned on a lab jack to enable vertical alignment with a microscope camera (Mighty Scope USB Digital Microscope, Aven Tools, Ann Arbor, MI, USA), which imaged the side view of the liquid droplets on the PDMS surface. An LED array with built-in diffuser (Amazon) was placed behind the sample surface, opposite the microscope camera, and used as a light source to create contrast between the sample and the background. A micropipette was used to deposit 2-µL droplets of liquid on the sample surface and droplet images were captured. Left and right static contact angles were extracted using open-source OpenDrop software (16,17). For each condition, static contact angles were measured for three replicate droplets.

#### S2.8. Characterizing bubbles in microfluidic devices

**Table S3.** Experimental conditions for evaluating the efficacy of bubble mitigation strategies in PDMS microfluidic devices.

| Condition | Degassing | Plasma treatment | Pre-wetting liquid | # of trials |
|---|---|---|---|---|
| **1** | PDMS not degassed | No plasma treatment | Ultrapure water | 8 |
| **2** | PDMS not degassed | No plasma treatment | Ethanol | 24 |
| **3** | PDMS not degassed | No plasma treatment | 0.3 mM Triton X-100 in PBS | 8 |
| **4** | PDMS not degassed | Plasma treatment | 0.3 mM Triton X-100 in PBS | 18 |
| **5** | PDMS degassed | No plasma treatment | Ethanol | 3 |
| **6** | PDMS degassed | Plasma treatment | 0.3 mM Triton X-100 in PBS | 18 |

Bubble nucleation and entry within the microfluidic channels during several hours of aqueous working liquid flow was visualized using a top-view microscope camera (Pixelink, Ottawa, ON, CAN) mounted to a 12× zoom lens (Navitar, Rochester, NY, USA), positioned above the PDMS chip-gasket assembly on the photonic testing stage. Prior to starting the experiment, the camera was focused on the microfluidic channels to ensure clear capture of any bubbles that may nucleate or enter the channels. The gain and exposure thresholds were adjusted to effectively distinguish between the bubbles and the surrounding fluid. During pre-wetting, Pixelink image capture software was used to capture images at 5-second intervals. During the experiment stage, images were captured at 20-second intervals. After the experiment, the captured images were sequentially stitched together to create a time-lapse video using a custom MATLAB script. The images were resized to 25% of their original scale and a MPEG-4 format time-lapse video was generated. The following metrics were manually tracked from the videos: number of bubble nucleation sites, number of bubbles entering and exiting the microfluidic channel, duration of time during which each bubble was present in the microfluidic channel, and maximum bubble size.

#### S2.9. Polydopamine film thickness

A preliminary experiment was performed to characterize the thickness of polydopamine (PDA) films deposited on silicon wafer as a function of deposition time. In this experiment, chips diced from unpatterned silicon wafer (type: P, dopant: B, orientation: <110>, resistivity: 1–10 Ω/cm, thickness: 500 µm, polish: SSP, grade: prime, Product # 2357, UniversityWafer, Inc., Boston, MA, USA) were used as the substrate for PDA deposition. Unpatterned silicon was selected as a model substrate to represent the silicon waveguide surfaces on our photonic sensors. To prepare the chips for coating, the wafer was protected with AZ 5214 photoresist, then diced into 9.7 × 9.7 mm squares. The resist was stripped using acetone, cleaned with Nanostrip for 15 min at 80ºC, rinsed with deionized water, then dried with nitrogen. The chips were stored in a petri dish lined with cleanroom wipe until ready for use. Immediately before use, the chips were cleaned by sequential immersion in acetone, isopropanol, and ultrapure water, then dried thoroughly with nitrogen gas.

Prior to PDA deposition, the thicknesses of the native silicon dioxide films on the silicon chips were measured using a variable angle spectroscopic ellipsometer (M-2000V, J. A. Woollam, Lincoln, NE, USA). Five measurements were performed at different locations on each chip to obtain a mean native oxide thickness across the chip surface. The five measurement locations were defined by a laser-cut alignment jig (Universal Laser Systems, Inc., Scottsdale, AZ, USA), fabricated from 0.025” Delrin® Acetal sheet (McMaster-Carr, Elmhurst, IL, USA) fixed to the ellipsometer stage (Fig. S4). Measurements were performed using an angle range of 50º–70º, with increments of 10, and 25 revs per measurement. Fitting was performed in WVASE32® software (J. A. Woollam) to determine the native oxide layer thickness using a Cauchy model over a wavelength range of 500–700 nm. The Cauchy model included 2 layers: (1) the 500-µm Si wafer and (2) the native $SiO_2$ film. A total of nine chips were measured, yielding an average native oxide film thickness of 1.8 ± 0.08 nm (reported as the mean across all nine chips ± standard deviation).

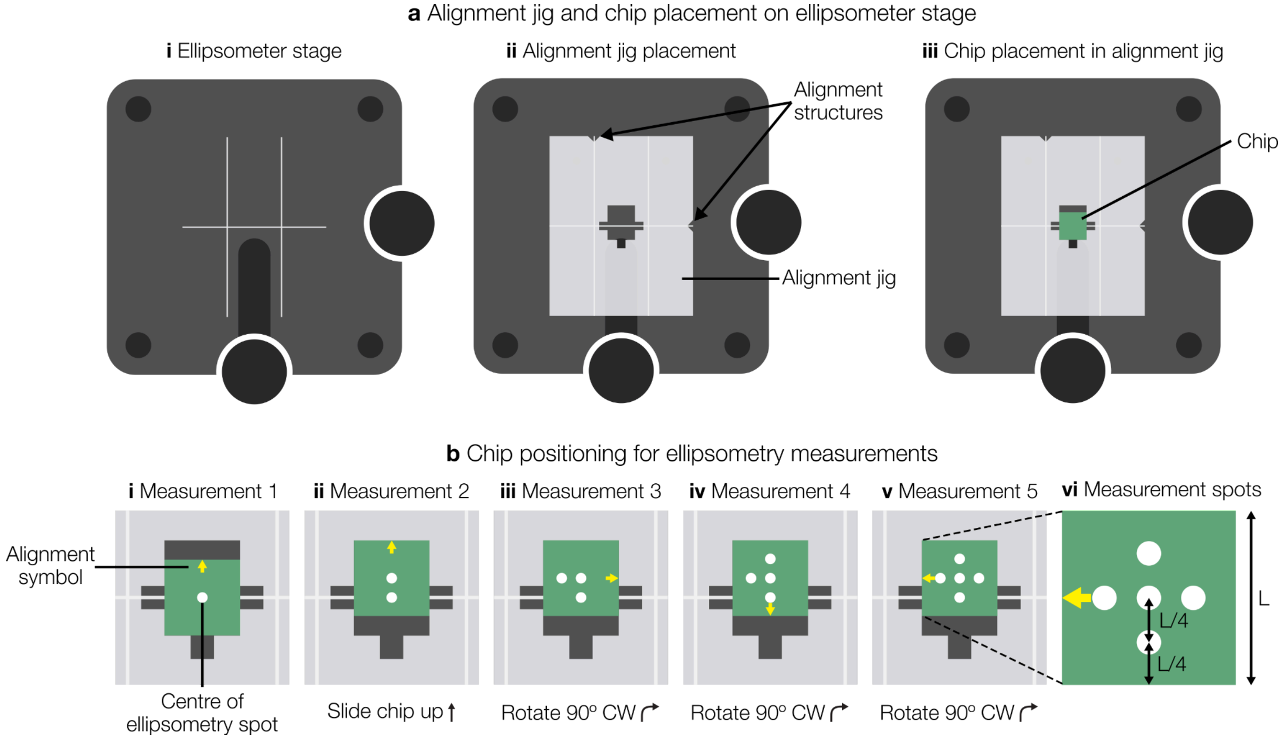

**Fig. S4.** Ellipsometry measurement procedure. **(a)** Illustration demonstrating the placement of the laser-cut alignment jig and silicon chip on the ellipsometer stage. **(b)** Illustration of chip positioning in the alignment jig to control the locations of each ellipsometry measurement and the resulting measurement spot locations.

Next, the chips were coated with PDA for 0.5, 1.0, or 1.5 h. Three replicate chips were used for each reaction time. The clean silicon chips were placed in a glass crystallization dish containing 40 mL of a freshly prepared dopamine hydrochloride solution (2 mg/mL in 0.5 M Tris buffer, pH 8.6). The chips were placed around the perimeter of the dish to avoid contact with a magnetic stir bar placed at the centre of the dish. PDA coating was allowed to proceed at room temperature under stirring at ~100 rpm with the solution open to atmospheric air. After the desired reaction time had elapsed, chips were removed from the dopamine hydrochloride solution, placed in a beaker containing fresh Tris buffer, thoroughly rinsed with ultrapure water, then dried with nitrogen gas.

The PDA films were measured via spectroscopic ellipsometry, as described for the native oxide thickness measurements. This time, the Cauchy model included 3 layers: the (1) 500-µm Si wafer, (2) native $SiO_2$ film, and (3) PDA adlayer. The previously measured native oxide thickness for each chip was inputted as the $SiO_2$ layer thickness and fitting was performed on the PDA adlayer. The results are displayed in Fig. S5, which shows that the film thickness increases almost linearly with deposition time. Reaction times of 0.5, 1.0, and 1.5 h yielded PDA films with thicknesses of 1.8 ± 0.1 nm, 4.1 ± 0.2 nm, and 6.1 ± 0.1 nm, respectively. It is generally desirable for surface functionalization chemistries on SiP microring resonators to be as thin as possible to ensure that binding reactions occur within the ~40–200 nm evanescent field of the silicon waveguides, while still achieving good uniformity and density of bioreceptors across the surface (18). Therefore, we selected the 30-min reaction time for use in the PDA-based sensor functionalization protocols reported in this work.

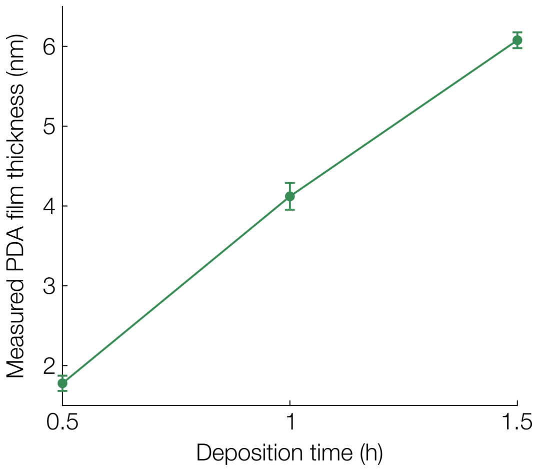

**Fig. S5.** Measured film thickness versus reaction time for PDA films deposited on unpatterned silicon chips. For each reaction time, data are plotted as the mean thickness measured via variable angle spectroscopic ellipsometry across three replicate chips ± standard deviation. For each chip, the film thickness was obtained as the average of five measurements taken at different locations across the chip surface.

#### S2.10. Bulk refractive index sensitivity testing

##### S2.10.1. Intrinsic sensor performance experiments and demonstration assays (20 µg/mL spike protein)

For each assay detecting 20 µg/mL spike protein using Protein A/flow-mediated functionalization, bulk RI sensing was completed prior to the assay to characterize each sensor's intrinsic sensing performance (Section 2.6.1). RI standard solutions of 0, 62.5, 125, 250, and 375 mM NaCl in ultrapure water were prepared and their refractive indices were measured using a visible light refractometer (Abbe-3L Refractometer, Spectronic Instruments, Rochester, NY, USA). The measured bulk refractive indices are reported in Table S4. The NaCl solutions were delivered in ascending and then descending order of concentration for 20 min per step at 30 µL/min. Resonance peak shift data from the ascending and descending concentrations were used to compute the bulk RI sensitivity, $S_{bulk}$, for each sensor. Table S5 reports the measured $S_{bulk}$ for each sensor in each trial, and Tables S6–S9 report the stability standard deviation ($\sigma_{\Delta\lambda}$), system limit of detection (sLoD), and baseline drift rate for these trials.

**Table S4.** Measured bulk refractive indices of the NaCl bulk RI standards used for the intrinsic sensor performance characterization, alongside comparison of measured refractive index step sizes (differences between adjacent NaCl concentrations, which is the metric that underlies the measured $S_{bulk}$) using the visible light refractometer with those predicted from curve fits to experimental data at 1550 nm. Refractometer measurements were conducted at room temperature (typically 22-24℃), while the published 1550 nm data were acquired between 24.0-26.7℃. Predicted RIs of the aqueous NaCl solutions at 1550 nm were calculated from their NaCl mass fractions in water (*w*) using the polynomial fit equation and fit parameters presented by Saunders *et al.* (19): $n = Aw^3 + Bw^2 + Cw + D$ ; A = -0.08, B = 0.074, C = 0.162, D = 1.3162. The differences between the average measured RI step sizes and the predicted step sizes range from 0.3-26%, and the predicted step sizes are all within 2 standard deviations of the average measured steps, suggesting that bulk RI measurement variation may dominate over the systematic error introduced through the use of the visible-light refractometer in these reference measurements.

| NaCl concentration (M) | NaCl mass fraction | Trial 1 measured RIs | Trial 2 measured RIs | Trial 3 measured RIs | Trial 4 measured RIs | Average measured RI step size | Standard deviation of measured RI step size | Saunders et al. (19) predicted RI (1550 nm) | Predicted RI step size |
|---|---|---|---|---|---|---|---|---|---|
| 0 (ultrapure water) | 0 | 1.3335 | 1.3335 | 1.3340 | 1.3335 | – | – | 1.31620 | – |
| 0.0625 | 0.0036473 | 1.3345 | 1.3341 | 1.3345 | 1.3341 | 0.0007 | 0.0002 | 1.31679 | 0.00059 |
| 0.125 | 0.0072681 | 1.3350 | 1.3350 | 1.3354 | 1.3350 | 0.0008 | 0.0002 | 1.31738 | 0.00059 |
| 0.25 | 0.0144313 | 1.3360 | 1.3362 | 1.3365 | 1.3364 | 0.00118 | 0.00017 | 1.31855 | 0.00117 |
| 0.375 | 0.0214919 | 1.3375 | 1.3375 | 1.3383 | 1.3379 | 0.0015 | 0.0002 | 1.31972 | 0.00116 |

**Table S5.** Sensor-by-sensor $S_{bulk}$, channels, and time delay between switch activation and apparent change in measured bulk RI (related to the internal volume of the fluid control system). These values were computed while excluding the bubble-containing regions from the analysis windows.

| Trial | Sensors | Channels | $S_{bulk}$ (nm/RIU) | | | | | | | | Time delay |
|---|---|---|---|---|---|---|---|---|---|---|---|
| | | | 1 | 2 | 3 | 4 | 5 | 6 | 7 | 8 | |
| 1 | 1, 2, 3, 4, 5, 6, 7, 8 | 2 | 490.7 | 458.1 | 441.3 | 460.0 | 482.3 | 457.7 | 460.2 | 458.4 | 6 mins |
| 2 | 1, 2, 3, 4, 5, 6, 7, 8 | 2 | 441.7 | 444.8 | 440.3 | 444.4 | 434.9 | 444.6 | 435.9 | 444.4 | 6 mins |
| 3 | 1, 2, 3, 4, 5, 6, 7, 8 | 2 | 427.6 | 438.4 | 427.7 | 436.1 | 427.3 | 435.6 | 426.8 | 435.6 | 6 mins |
| 4 | 1, 2, 3, 4 | 2 | 419.4 | 420.4 | 419.8 | 420.0 | | | | | 6 mins |

**Table S6.** Sensor-by-sensor $\sigma_{\Delta\lambda}$ measurements. These values were computed while excluding the bubble-containing regions from the analysis windows.

| Trial | Sensors | Channels | $\sigma_{\Delta\lambda}$ (nm) | | | | | | | |
|---|---|---|---|---|---|---|---|---|---|---|
| | | | 1 | 2 | 3 | 4 | 5 | 6 | 7 | 8 |
| 1 | 1, 2, 3, 4, 5, 6, 7, 8 | 2 | $5.43 \times 10^{-3}$ | $6.26 \times 10^{-4}$ | $5.58 \times 10^{-3}$ | $5.75 \times 10^{-4}$ | $6.72 \times 10^{-3}$ | $6.91 \times 10^{-4}$ | $5.81 \times 10^{-3}$ | $2.00 \times 10^{-3}$ |
| 2 | 1, 2, 3, 4, 5, 6, 7, 8 | 2 | $9.23 \times 10^{-4}$ | $1.40 \times 10^{-3}$ | $1.08 \times 10^{-3}$ | $1.02 \times 10^{-3}$ | $5.93 \times 10^{-4}$ | $1.28 \times 10^{-3}$ | $2.60 \times 10^{-4}$ | $1.28 \times 10^{-3}$ |
| 3 | 1, 2, 3, 4, 5, 6, 7, 8 | 2 | $5.56 \times 10^{-4}$ | $1.09 \times 10^{-2}$ | $1.00 \times 10^{-3}$ | $1.20 \times 10^{-2}$ | $5.13 \times 10^{-4}$ | $1.16 \times 10^{-2}$ | $7.83 \times 10^{-4}$ | $1.18 \times 10^{-2}$ |
| 4 | 1, 2, 3, 4 | 2 | $7.45 \times 10^{-4}$ | $1.26 \times 10^{-3}$ | $9.72 \times 10^{-4}$ | $1.55 \times 10^{-3}$ | | | | |

**Table S7.** Sensor-by-sensor sLoD measurements. These values were computed while excluding the bubble-containing regions from the analysis windows.

| Trial | Sensors | Channels | sLoD (RIU × $10^{-6}$) | | | | | | | |
|---|---|---|---|---|---|---|---|---|---|---|
| | | | 1 | 2 | 3 | 4 | 5 | 6 | 7 | 8 |
| 1 | 1, 2, 3, 4, 5, 6, 7, 8 | 2 | 33.2 | 4.10 | 37.9 | 3.75 | 41.8 | 4.53 | 37.9 | 13.1 |
| 2 | 1, 2, 3, 4, 5, 6, 7, 8 | 2 | 6.28 | 9.47 | 7.33 | 6.86 | 4.08 | 8.64 | 1.79 | 8.61 |
| 3 | 1, 2, 3, 4, 5, 6, 7, 8 | 2 | 3.90 | 75.0 | 7.03 | 82.5 | 3.60 | 79.7 | 5.50 | 81.5 |
| 4 | 1, 2, 3, 4 | 2 | 5.33 | 8.96 | 6.95 | 11.1 | | | | |

**Table S8.** Sensor-by-sensor initial baseline drift rate measurements. These values were computed while excluding the bubble-containing regions from the analysis windows.

| Trial | Sensors | Channels | Baseline drift rate (pm/min) | | | | | | | |
|---|---|---|---|---|---|---|---|---|---|---|
| | | | 1 | 2 | 3 | 4 | 5 | 6 | 7 | 8 |
| 1 | 1, 2, 3, 4, 5, 6, 7, 8 | 2 | $-1.81 \times 10^{-3}$ | $-1.17 \times 10^{-5}$ | $-1.85 \times 10^{-3}$ | $-6.94 \times 10^{-5}$ | $-2.11 \times 10^{-3}$ | $-1.05 \times 10^{-4}$ | $-1.92 \times 10^{-3}$ | $-3.59 \times 10^{-4}$ |
| 2 | 1, 2, 3, 4, 5, 6, 7, 8 | 2 | $2.92 \times 10^{-4}$ | $-3.41 \times 10^{-4}$ | $3.40 \times 10^{-4}$ | $-3.21 \times 10^{-4}$ | $1.77 \times 10^{-4}$ | $-4.02 \times 10^{-4}$ | $6.09 \times 10^{-5}$ | $-4.08 \times 10^{-4}$ |
| 3 | 1, 2, 3, 4, 5, 6, 7, 8 | 2 | $9.75 \times 10^{-5}$ | $3.44 \times 10^{-3}$ | $2.56 \times 10^{-4}$ | $3.81 \times 10^{-3}$ | $1.45 \times 10^{-4}$ | $3.68 \times 10^{-3}$ | $2.44 \times 10^{-4}$ | $3.76 \times 10^{-3}$ |
| 4 | 1, 2, 3, 4 | 2 | $2.29 \times 10^{-4}$ | $3.97 \times 10^{-4}$ | $3.05 \times 10^{-4}$ | $4.97 \times 10^{-4}$ | | | | |

**Table S9.** Sensor-by-sensor data for the Trial 1 intrinsic sensor performance characterization including the bubble-containing period in the analysis window.

| Metric | Units | Sensors | | | | | | | |
|---|---|---|---|---|---|---|---|---|---|
| | | 1 | 2 | 3 | 4 | 5 | 6 | 7 | 8 |
| $S_{bulk}$ | nm/RIU | 474.4 | 458.8 | 439.2 | 460.8 | 456.0 | 458.3 | 446.5 | 459.5 |
| $\sigma_{\Delta\lambda}$ | nm | 0.140 | $5.97 \times 10^{-4}$ | 0.0363 | $5.99 \times 10^{-4}$ | 0.217 | $2.98 \times 10^{-4}$ | 0.120 | $3.87 \times 10^{-3}$ |
| sLoD | RIU | $8.82 \times 10^{-4}$ | $3.90 \times 10^{-6}$ | $2.48 \times 10^{-4}$ | $3.90 \times 10^{-6}$ | $1.43 \times 10^{-4}$ | $1.95 \times 10^{-6}$ | $8.06 \times 10^{-4}$ | $2.53 \times 10^{-5}$ |
| Baseline drift rate | pm/min | 0.035 | $-1.90 \times 10^{-4}$ | $3.34 \times 10^{-3}$ | $1.22 \times 10^{-4}$ | 0.0571 | $-6.81 \times 10^{-6}$ | 0.0288 | $-1.15 \times 10^{-3}$ |

##### S2.10.2. Demonstration assays (1 µg/mL spike protein)

All assays demonstrating 1 µg/mL spike protein detection were followed by bulk RI sensing to quantify $S_{bulk}$ for each sensor. RI standard solutions of 0, 125, and 250 mM NaCl in ultrapure water were prepared and their refractive indices were measured using a refractometer. The NaCl solutions were delivered in ascending and then descending order of concentration for 20 min per step at 30 µL/min. This sequence was repeated twice. Resonance peak shift data from the ascending and descending concentrations were used to compute $S_{bulk}$ for each sensor. Refractive indices of the NaCl solutions and quantified $S_{bulk}$ values for each trial are provided in Table S10.

**Table S10.** Bulk refractive indices of NaCl standard solutions and measured $S_{bulk}$ for sensors used in spike protein detection assays demonstrating 1 µg/mL spike protein detection.

| | | Protein A/flow | PDA/spotting | | | PDA/flow | | |
|---|---|---|---|---|---|---|---|---|
| | | **Trial 1** | **Trial 1** | **Trial 2** | **Trial 3** | **Trial 1** | **Trial 2** | **Trial 3** |
| **Measured RIs** | **Ultrapure water (0 M NaCl)** | 1.3336 | 1.3336 | 1.3336 | 1.3336 | 1.3336 | 1.3336 | 1.3336 |
| | **125 mM NaCl** | 1.3350 | 1.3349 | 1.3349 | 1.3349 | 1.3350 | 1.3350 | 1.3350 |
| | **250 mM NaCl** | 1.3361 | 1.3360 | 1.3360 | 1.3360 | 1.3361 | 1.3361 | 1.3361 |
| **$S_{bulk}$ (nm/RIU)*** | | 433 ± 1$^{\dagger}$ | 393 ± 4 | 390 ± 13 | 415 ± 5 | 384 ± 16 | 368 ± 14 | 397 ± 20 |

*$S_{bulk}$ values are reported as the intra-assay mean and standard deviation, calculated across 8 replicate SiP MRR sensors in two fluidic channels in each trial.

$^{\dagger}S_{bulk}$ was calculated for resonators in fluidic channel 2 only, as the 10-channel bidirectional valve in channel 1 of the fluidic control system malfunctioned during the bulk RI sensing portion of this assay.

#### S2.11. Reservoir setups and fluidic protocols for demonstration assays

Bubble mitigation strategies employed for each assay format are summarized in Table S11. For each assay, the fluidic delivery system was loaded with the appropriate functionalization and assay reagents and primed with the fluids outlined in Table S12. Assays were then performed using the automated fluidic protocols outlined in Table S13. For all assays, the same microfluidic protocol was delivered to both microfluidic channels using different channels of the 2-channel fluid delivery system, with each channel having its own set of input reservoirs (prepared as duplicates by splitting each prepared solution between the two channels).

After priming and pre-wetting, Protein A/flow assays began with a 30-min PBS running buffer stabilization period, followed by Protein A deposition (100 µg/mL in PBS) to facilitate oriented immobilization of capture antibody on the sensor surface, BSA (1 mg/mL in PBS) to test for nonspecific binding and block any exposed sensor surface not covered by Protein A, and anti-spike protein capture antibody deposition (20 µg/mL in PBS). Each functionalization step was followed by a 10-min PBS rinse. Note that a lower antibody concentration was used for flow-based functionalization (20 µg/mL) compared to spotting-based functionalization (500 µg/mL) such that all sensor chips were exposed to a total mass of 10 µg of capture antibody, regardless of functionalization approach. After functionalization, spike protein detection (1 µg/mL or 20 µg/mL in PBS) was performed, followed by a final PBS running buffer stabilization period.

PDA/spotting assays began with a 30-min PBS-BSA running buffer stabilization period. PBS-BSA was used as the running buffer in these assays to mitigate the removal of surface blocking applied to the chip prior to the assay. Next, the sensor was exposed to glycine-HCl buffer to remove excess loosely bound capture antibodies (20,21), followed by a second PBS-BSA running buffer stabilization period. The sensor was challenged with BSA (1 mg/mL in PBS), rinsed with PBS-BSA for 10 min, exposed to spike protein (1 µg/mL in PBS-BSA), then underwent a final PBS-BSA running buffer stabilization period.

PDA/flow assays began with a 30-min PBS stabilization period, followed by antibody immobilization (20 µg/mL in PBS), and BSA blocking (20 mg/mL in PBS). Each functionalization step was followed by a PBS rinse. After a PBS-BSA running buffer stabilization period, the sensor was exposed to glycine-HCl buffer, followed by a second PBS-BSA stabilization period. The sensor was challenged with BSA (1 mg/mL in PBS), rinsed with PBS-BSA for 10 min, exposed to spike protein (1 µg/mL in PBS-BSA), then underwent a final PBS-BSA running buffer stabilization period.

**Table S11.** Gasket treatment conditions and pre-wetting fluids used for analyte detection assays.

| Assay type | Degassing | Plasma treatment | Pre-wetting liquid |
|---|---|---|---|
| Protein A/flow, 20 µg/mL spike protein | PDMS not degassed | No plasma treatment | Ethanol |

| | | | |
|---|---|---|---|
| Protein A/flow, 1 µg/mL spike protein | PDMS degassed | Plasma treatment | 0.3 mM Triton X-100 in PBS |
| PDA/spotting, 1 µg/mL spike protein | PDMS degassed | Plasma treatment | 0.3 mM Triton X-100 in PBS |
| PDA/flow, 1 µg/mL spike protein | PDMS degassed | Plasma treatment | 0.3 mM Triton X-100 in PBS |

**Table S12.** Fluidic control system reservoir setup for analyte detection assays.

| Reservoir | Protein A/flow (20 µg/mL spike protein) | | Protein A/flow (1 µg/mL spike protein) | | PDA/spotting | | PDA/flow | |
|---|---|---|---|---|---|---|---|---|
| | Assay | Priming | Assay | Priming | Assay | Priming | Assay | Priming |
| 1 | Ultrapure water | ⇇ | Ultrapure water | ⇇ | 0.3 mM Triton X-100 in PBS | ⇇ | PBS/ultrapure water* | PBS |
| 2 | 62.5 mM NaCl | ⇇ | 125 mM NaCl | ⇇ | 125 mM NaCl | ⇇ | 125 mM NaCl | ⇇ |
| 3 | 125 mM NaCl | ⇇ | 250 mM NaCl | ⇇ | 250 mM NaCl | ⇇ | 250 mM NaCl | ⇇ |
| 4 | 250 mM NaCl | ⇇ | 0.3 M Triton X-100 in PBS | ⇇ | Ultrapure water | ⇇ | 0.3 mM Triton X-100 in PBS | ⇇ |
| 5 | 375 mM NaCl | ⇇ | PBS | ⇇ | 1 mg/mL BSA in PBS | ⇇ | 1 mg/mL BSA in PBS | ⇇ |
| 6 | PBS | ⇇ | 1 mg/mL BSA in PBS | ⇇ | PBS-BSA | ⇇ | PBS-BSA | ⇇ |
| 7 | 1 mg/mL BSA in PBS | PBS | 100 µg/mL Protein A in PBS | PBS | 1 µg/mL spike protein in PBS-BSA | PBS-BSA | 1 µg/mL spike protein in PBS-BSA | PBS-BSA |
| 8 | 100 µg/mL Protein A in PBS | PBS | 20 µg/mL capture antibody in PBS | PBS | Glycine-HCl buffer | ⇇ | Glycine-HCl buffer | ⇇ |
| 9 | 20 µg/mL capture antibody in PBS | PBS | 1 µg/mL spike protein in PBS | PBS | PBS-BSA | ⇇ | 20 mg/mL BSA in PBS | ⇇ |
| 10 | 20 µg/mL spike protein in PBS | PBS | Glycine-HCl buffer (unused) | ⇇ | PBS-BSA | ⇇ | 20 µg/mL capture antibody in PBS | PBS |

⇇ These reservoirs were primed with the assay reagents.

*This reservoir contained PBS during the functionalization portion of the assay and was later switched to ultrapure water once functionalization was complete.

**Table S13.** Automated fluidic protocols for analyte detection assays.

| Assay step | Protein A/flow assays | | | PDA/spotting assays | | | PDA/flow assays | | |
|---|---|---|---|---|---|---|---|---|---|
| | Solution | Flow rate (µL/min) | Flow time (min) | Solution | Flow rate (µL/min) | Flow time (min) | Solution | Flow rate (µL/min) | Flow time (min) |
| 1 | PBS | 30 | 60 | PBS-BSA | 30 | 30 | PBS | 30 | 30 |
| 2 | Protein A | 30 | 17 | Glycine-HCl buffer | 30 | 7 | Antibody | 20 | 25 |
| 3 | PBS | 30 | 10 | PBS-BSA | 30 | 30 | PBS | 30 | 10 |
| 4 | 1 mg/mL BSA | 30 | 22 | 1 mg/mL BSA | 30 | 20 | 20 mg/mL BSA | 30 | 25 |
| 5 | PBS | 30 | 10 | PBS-BSA | 30 | 10 | PBS | 30 | 30 |
| 6 | Antibody | 20 | 25 | Spike protein | 30 | 17 | PBS-BSA | 30 | 40 |
| 7 | PBS | 30 | 10 | PBS-BSA | 30 | 30 | Glycine-HCl buffer | 30 | 7 |
| 8 | Spike protein | 30 | 17 | – | – | – | PBS-BSA | 30 | 30 |
| 9 | PBS | 30 | 30 | – | – | – | 1 mg/mL BSA | 30 | 20 |
| 10 | – | – | – | – | – | – | PBS/BSA | 30 | 10 |
| 11 | – | – | – | – | – | – | Spike protein | 30 | 17 |
| 12 | – | – | – | – | – | – | PBS-BSA | 30 | 30 |

#### S2.12. Sensing data analysis workflow

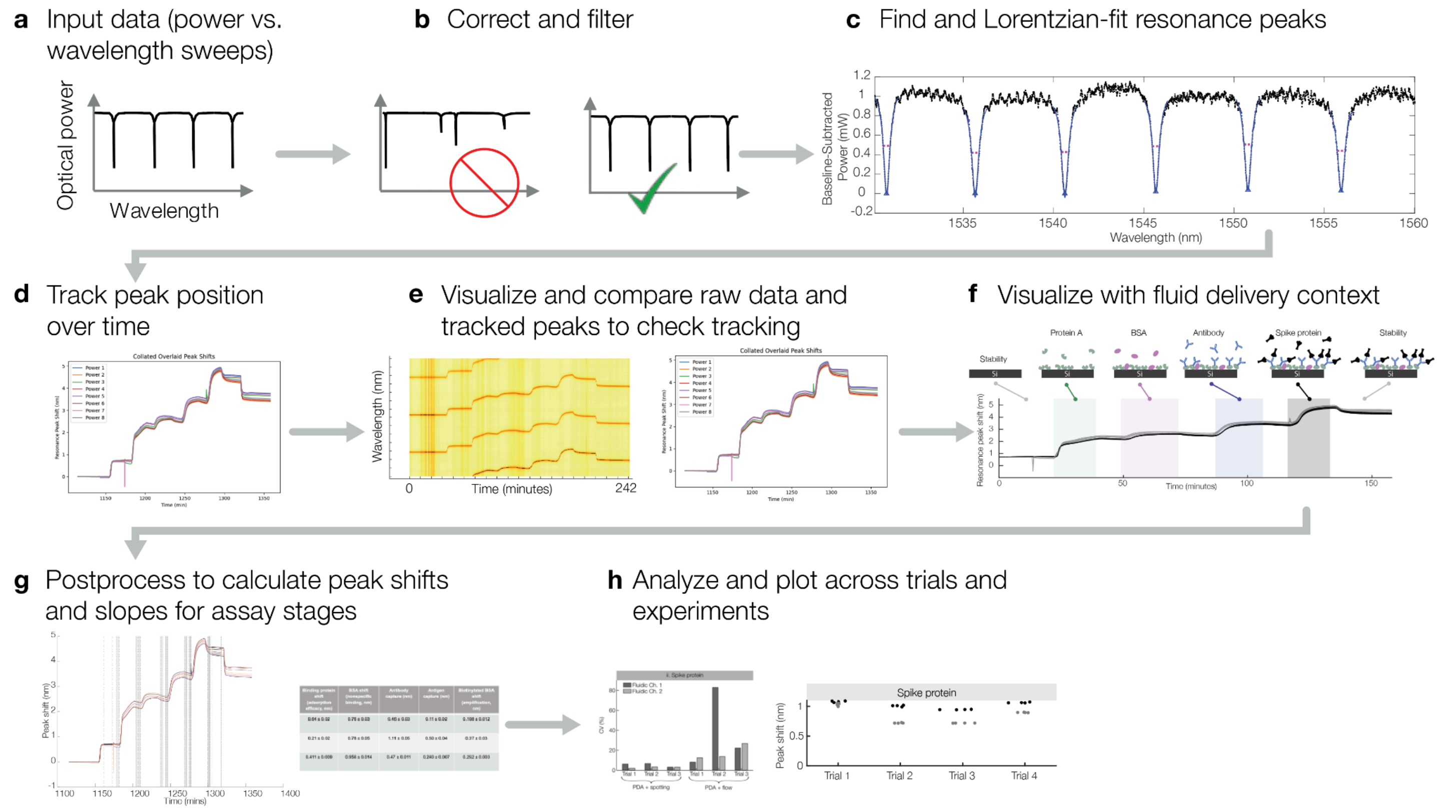

**Fig. S6.** Overview of the data analysis workflow. **(a)** individual laser sweeps at each timepoint (sampling every ~20 s) plot optical power vs. wavelength and contain the resonance peak information that is tracked to analyze the biosensor data. **(b)** The individual laser sweep data are filtered and corrected to remove sweeps that do not contain a pattern corresponding to resonance peaks, prior to tracking. **(c)** Resonance peaks are found and Lorentzian-fitted to extract a vector of information regarding each peak (central wavelength, amplitude, width, baseline), **(d)** The peak shifts over time are quantified by matching resonance peaks in consecutive sweeps based on their information vectors. **(e)** a "spectrogram" (optical power data is represented on a colour scale and each individual sweep is represented as a column of an image) is generated from the raw data, and the peak tracking data are overlaid and visually compared with the spectrogram to ensure accurate tracking. **(f)** Fluid delivery information is overlaid on the peak shift vs. time data by plotting the log file data from the fluidic delivery system. **(g)** Relevant shifts and slopes are quantified from the peak shift vs. time data. **(h)** Quantified peak shift and slope data from multiple trials are collated and compared.

##### S2.12.1. Resonance peak tracking

Acquired optical spectra were analyzed using a custom Python script to Lorentzian-fit each resonance peak and track cumulative peak shifts (Fig. S6 (a–c)). Plots and datasets of average resonance peak shift vs. time were generated for each measured microring resonator sensor (Fig. S6 (d)). Further details regarding this data processing workflow have been described previously (12).

##### S2.12.2. Bulk refractive index sensitivity extraction

Bulk RI sensitivity, $S_{bulk}$, values were computed from resonance peak shift vs. time data collected during exposure to NaCl standard solutions (Section 2.6.1) using a custom MATLAB script, as described previously by our group (12).

##### S2.12.3. Baseline correction

After processing the individual resonance spectra to yield the sensorgram (resonance peak shift vs. time) data for all binding assay experiments, these data were baseline-corrected. The purpose of this correction was to mitigate the effects of any sensor drift and functionalization dissociation on the subsequent quantification of peak shifts and baseline drift slopes for each assay stage. Using a custom MATLAB script, the sensorgram peak shift ($y$) vs. time ($t$) data for an initial ~15-min running buffer equilibration period (either PBS or PBS-BSA, depending on the assay) was nonlinear least-squares fitted to a modified bimolecular dissociation model (22) incorporating additional fit coefficients for equilibrium ($t = \infty$) peak shift ($d$) and time offset ($c$) in addition to the off-rate of the binding interaction ($c$) and the maximum signal from the bound fraction ($a$):

$$y_{fit} = a \cdot \left(1 - e^{b(t-c)}\right) + d, \qquad \text{Eq. S1}$$

The baseline data to be fitted were manually selected by clicking on a plot of the sensorgram data, and the script displayed the results of the curve fit, overlaid data, and residuals for inspection. The script generated a new baseline-corrected dataset ($y_{corr}$) equal to the fitted curve ($y_{fit}$) subtracted from the original sensorgram data ($y$): $y_{corr} = y - y_{fit}$ for the full time range (t = 0 to assay end). Supplementary Information Section S8 presents the baseline fit parameters for all of the binding assay data reported in this work, as well as the expected resulting peak shift over a 20-min time period at the end of the baseline correction period and 100 minutes later. Correction shifts are small (< 93 pm immediately after correction; < 17 pm 100 mins later) and thus expected to have low impact on most of our quantified shifts in Figures 5-8, but this correction does allow us to resolve small shifts in some cases that may be negative prior to baseline-correction.

These baseline-corrected data were used for subsequent quantification of binding peak shifts and baseline drift slopes. Plots of these baseline-corrected data were also overlaid with coloured boxes to visualize the fluid being delivered to the sensors at each assay stage (e.g., Fig. 5(a)). The start and end time points for these overlay regions were defined based on the log files from the Fluigent system alongside a delay factor that took into account the integrated measured flow rates (from the Fluigent log files) and the estimated internal volume between the Fluigent flow sensor and the microfluidic gasket. The estimated internal volume varied by assay depending on whether the ethanol pre-wetting assembly was present in the fluidic path (~390 µL with the assembly in place; ~150 µL without).

##### S2.12.4. Peak shifts and slopes quantification

After baseline-correction, a custom semi-automated MATLAB script was used to quantify the initial and final drift rate stability slopes as well as the binding signal shifts for each assay stage from the corrected sensorgram data. This script creates an overlay plot of the sensorgram data for all 8 sensors used in the assay and allows the user to click on (1) start and end timepoints to define regions over which to calculate linear slopes of the peak shift vs. time data (used here to extract drift rate stability slopes), or (2) pre-binding and post-binding regions used here to calculate resonance peak shift signal due to binding interactions.

To estimate the linear slopes of the data, the x (time) and y (baseline-corrected peak shift) data in the region defined for slope calculation were fitted to a linear regression model using MATLAB's standard 'fitlm' function. The fitted slope and the error (standard error estimate on the slope fit coefficient) were outputted by the script.

To estimate the binding peak shifts, the user clicked on the sensorgram data plot in the running buffer wash regions before and after the assay stage to be quantified, and the script averaged the peak shift vs. time data for 10 data points surrounding each click position (representing ~3.5 min of continuous data acquisition) to yield the pre-binding ($y_{pre}$) and post-binding ($y_{post}$) peak positions and their errors ( $\sigma_{y_{pre}}$ and $\sigma_{y_{post}}$, representing the standard deviations of the averaged data points in the pre- and post-binding regions). The quantified peak shift for each sensor and each assay stage was output by the script as $y_{post} - y_{pre}$, and the error on the quantified peak shift for each sensor and each assay stage was $\sqrt{\sigma_{y_{pre}}^2 + \sigma_{y_{post}}^2}$. Average peak shifts were reported as mean ± standard deviation across all sensors in the channel/trial.

The peak shift and slope data were subsequently post-processed to calculate the intra- and inter-assay CVs and plotted using Excel and MATLAB. The intra-assay CVs were calculated as the absolute value of the standard deviation of the binding shifts or slopes for all sensors (n = 8) within a single assay stage (or all sensors in a single microfluidic channel, n = 4) divided by the mean value across those same sensors. For the inter-assay CVs, we first calculated the average shift or slope across all detectors in an assay (n = 8) and subsequently calculated the inter-assay CV as the standard deviation of these averaged shifts/slopes across all trials (n = 3–4 trials) divided by the mean shift/slope across all trials.

#### S3. Statistical analysis

##### S3.1. Bubble mitigation strategy comparisons

Two-tailed Mann-Whitney U tests were performed in MATLAB (Natick, MA, USA) to evaluate the statistical significance of differences in bubble mitigation performance metrics (bubbles nucleating per hour and bubbles entering the channel per hour) between the different PDMS gasket treatment and pre-wetting strategies. Pre-wetting using Triton X-100 solution with and without plasma treatment and with and without degassing, pre-wetting using ethanol and no plasma treatment, and pre-wetting using ultrapure water and no plasma treatment were compared to one-another in a pairwise manner. Mann-Whitney U tests were performed using a significance level of $\alpha = 0.05$. Results are presented in Tables S14–S16.

**Table S14.** Statistical analysis of the bubble mitigation data presented in Fig. 3(c) (effect of plasma treatment). P-values reported in bolded text indicate metrics that were found to have a statistically significant difference between the bubble mitigation approaches based on a significance level of $\alpha = 0.05$.

| | | **Conditions compared** | | | **P-values for comparison metrics** | | |
|---|---|---|---|---|---|---|---|
| | | **Pre-wetting liquid** | **Plasma** | **Degas** | **Bubbles nucleated** | **Bubbles entering** | **Bubble duration** |
| **Triton vs. Triton + plasma** | **Group 1** | 0.3 mM Triton in PBS | N | N | 0.16 | **0.049** | 0.08 |
| | **Group 2** | 0.3 mM Triton in PBS | Y | N | | | |

**Table S15.** Statistical analysis of the bubble mitigation data presented in Fig. 3(d) (effect of gasket degassing).

| | | **Conditions compared** | | | **P-values for comparison metrics** | | |
|---|---|---|---|---|---|---|---|
| | | **Pre-wetting liquid** | **Plasma** | **Degas** | **Bubbles nucleated** | **Bubbles entering** | **Bubble duration** |
| **Triton + plasma vs. EtOH** | **Group 1** | 0.3 mM Triton in PBS | Y | N | 0.13 | 0.76 | 0.63 |
| | **Group 2** | 100% Ethanol | N | N | | | |
| **Triton + plasma vs. Triton + plasma + degas** | **Group 1** | 0.3 mM Triton in PBS | Y | N | 1 | 0.76 | 0.57 |
| | **Group 2** | 0.3 mM Triton in PBS | Y | Y | | | |
| **EtOH vs. EtOH + degas** | **Group 1** | 100% Ethanol | N | N | 0.57 | 0.49 | 0.49 |
| | **Group 2** | 100% Ethanol | N | Y | | | |
| **EtOH + degas vs. Triton + plasma + degas** | **Group 1** | 100% Ethanol | N | Y | 1 | 0.42 | 0.42 |
| | **Group 2** | 0.3 mM Triton in PBS | Y | Y | | | |

**Table S16.** Statistical analysis of the bubble mitigation data presented in Fig. 3(e) (effect of pre-wetting liquid). P-values reported in bolded text indicate metrics that were found to have a statistically significant difference between the functionalization approaches based on a significance level of $\alpha = 0.05$.

| | | **Conditions compared** | | | **P-values for comparison metrics** | | |
|---|---|---|---|---|---|---|---|
| | | **Pre-wetting liquid** | **Plasma** | **Degas** | **Bubbles nucleated** | **Bubbles entering** | **Bubble duration** |
| **Water vs. EtOH** | **Group 1** | Ultrapure water | N | N | **$3.5 \times 10^{-4}$** | **0.02** | **$7.9 \times 10^{-4}$** |
| | **Group 2** | 100% Ethanol | N | N | | | |
| **Water vs. Triton** | **Group 1** | Ultrapure water | N | N | **$3.5 \times 10^{-2}$** | 0.28 | 0.11 |
| | **Group 2** | 0.3 mM Triton in PBS | N | N | | | |
| **Water vs. Triton + plasma + degas** | **Group 1** | Ultrapure water | N | N | **$5.7 \times 10^{-5}$** | 0.05 | **$3.5 \times 10^{-5}$** |
| | **Group 2** | 0.3 mM Triton in PBS | Y | Y | | | |
| **EtOH vs. Triton** | **Group 1** | 100% Ethanol | N | N | 0.70 | **0.01** | **0.05** |
| | **Group 2** | 0.3 mM Triton in PBS | N | N | | | |
| **EtOH vs. Triton + plasma + degas** | **Group 1** | 100% Ethanol | N | N | 0.23 | 0.57 | 0.90 |
| | **Group 2** | 0.3 mM Triton in PBS | Y | Y | | | |
| **Triton vs. Triton + plasma + degas** | **Group 1** | 0.3 mM Triton in PBS | N | N | 0.16 | **0.02** | **0.01** |
| | **Group 2** | 0.3 mM Triton in PBS | Y | Y | | | |

#### S3.2. Demonstration assay comparisons

Two-tailed Mann-Whitney U tests were performed in MATLAB (Natick, MA, USA) using the ranksum function and a significance level of $\alpha = 0.05$ to evaluate the statistical significance of differences in demonstration assay performance metrics (initial and final drift rates, BSA binding shifts, and spike protein detection shifts) between the different functionalization strategies. Due to the relatively low number of assay replicates analyzed in this study for each functionalization approach, we decided to perform statistical analysis using the non-parametric Mann-Whitney U test, which does not make any assumptions about whether or not the data are normally distributed (23). Protein A/flow (using 20 μg/mL spike protein), PDA/spotting, and PDA/flow assays were compared to one-another in a pairwise manner. The Protein A/flow assay involving 1 μg/mL spike protein detection was not included in this statistical analysis, as only a single replicate was performed for this assay format. For these tests, the null hypothesis was stated as $H_0$: there is no statistical difference in the given assay performance metric between the two functionalization approaches. Shift and slope data for each of the 8 MRR sensors in each trial were used as the input data to the significance test for each functionalization strategy. The resulting p-values for each test are provided in Table S17.

**Table S17.** P-values obtained from two-tailed Mann-Whitney U tests comparing assay performance metrics for different functionalization approaches. P-values reported in bolded text indicate metrics that were found to have a statistically significant difference between the functionalization approaches based on a significance level of $\alpha = 0.05$.

| | **Protein A/flow vs. PDA/spotting** | **Protein A/flow vs. PDA/flow** | **PDA/spotting vs. PDA/flow** |
|---|---|---|---|
| **Initial drift rate** | 0.85 | 0.55 | 0.059 |
| **BSA challenge shift** | **0.0058** | **0.0016** | **$4.1 \times 10^{-4}$** |
| **Spike protein detection shift** | **$2.2 \times 10^{-10}$*** | **$2.2 \times 10^{-10}$*** | **$3.1 \times 10^{-9}$** |
| **Final drift rate** | **$2.9 \times 10^{-9}$** | **$2.2 \times 10^{-10}$** | **$4.9 \times 10^{-8}$** |

*Note that different concentrations of spike protein were employed for the Protein A/flow (20 μg/mL) and PDA/spotting and PDA/flow (1 μg/mL) assays.

### S4. Sensor instability due to bubble exposure

In trial 1 of the four 20 μg/mL SARS-CoV-2 spike protein assays using Protein A/flow functionalization, there were bubbles nucleating and detaching around the inlet of channel 2 throughout the bulk RI sensitivity characterization and binding assay, directly affecting sensors 1, 3, 4, and 5. As described in Section 3.2, the data were analyzed both by including the periods in which the bubbles impacted the sensor signal, as well as by selecting periods that excluded the bubble(s) from analysis. Fig. S7 depicts micrographs collected by the experiment-monitoring time-lapse capture that show the bubble in the microfluidic channel as well as the corresponding regions of the peak shift plot where the resonance peak wavelength is impacted by the bubble. As the bubble passes over the sensors, an abrupt increase in resonance peak wavelength is observed for all channel 2 sensors, followed by a decrease as the bubble is cleared from the channel.

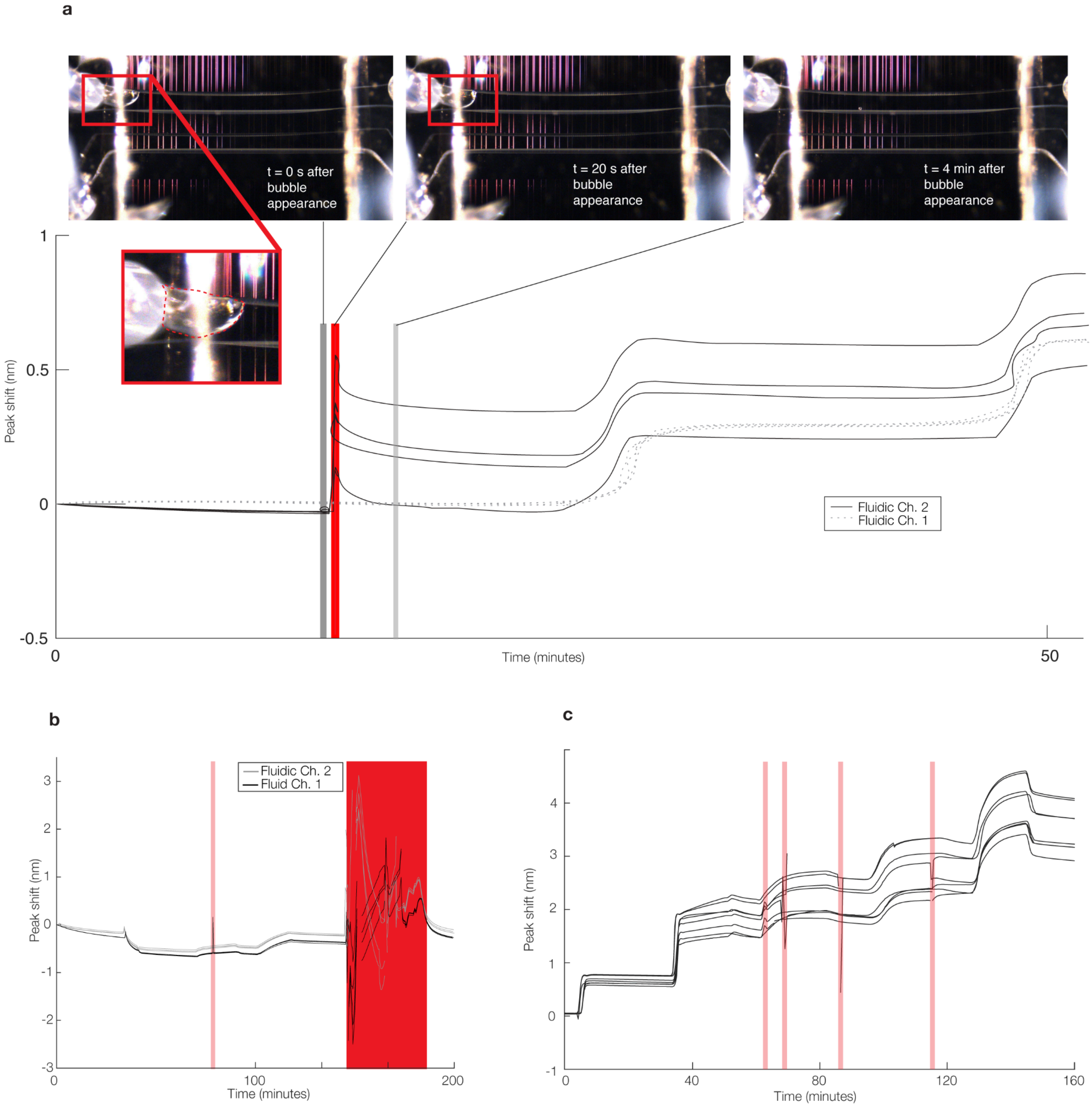

**Fig. S7.** Micrographs depicting instability caused by bubbles entering the microfluidic channel(s). **(a)** Micrographs depicting the bubble that impacted fluidic channel 2 in trial 1 of the four 20 μg/mL SARS-CoV-2 spike protein assays using Protein A/flow functionalization. Regions of the peak shift plot corresponding to each micrograph are labelled (n = 8 sensors). **(b)** Micrograph depicting instability due to an equipment malfunction that resulted in air introduction to the microfluidic channels during a PDA-mediated assay. Equipment malfunction discontinuities in peak shift measurement are highlighted in dark red and shorter, non-discontinuity-causing instability periods caused by bubbles and characterized by sudden increase then normalization of peak shift back to the baseline are highlighted in light red (n = 8 sensors). **(c)** Micrograph of bubble-induced instability during a Protein A mediated assay. Brief regions of bubble-induced instability are highlighted in light red (n = 8 sensors). Although the peak shift baseline returns to its previous position in some cases after the bubbles pass through, in other cases there is a more permanent offset introduced by the bubble, potentially due to residual air or damage to the functionalized surface.

#### S5. Sensor signal dependence on channel position

To assess whether analyte depletion along the length of the microfluidic channel might increase our measured intra-assay CVs, we plotted the 20 μg/mL spike protein detection signal as a function of sensor position along the length of the microfluidic channel for each channel-replicate of the data presented in Fig. 5. These data are presented in Fig. S8 below. No obvious relationship between sensor signal and position along the length of the channel was observed, suggesting that analyte is not depleted along the ~800 μm traveled by the fluid between the first and last sensor in the group.

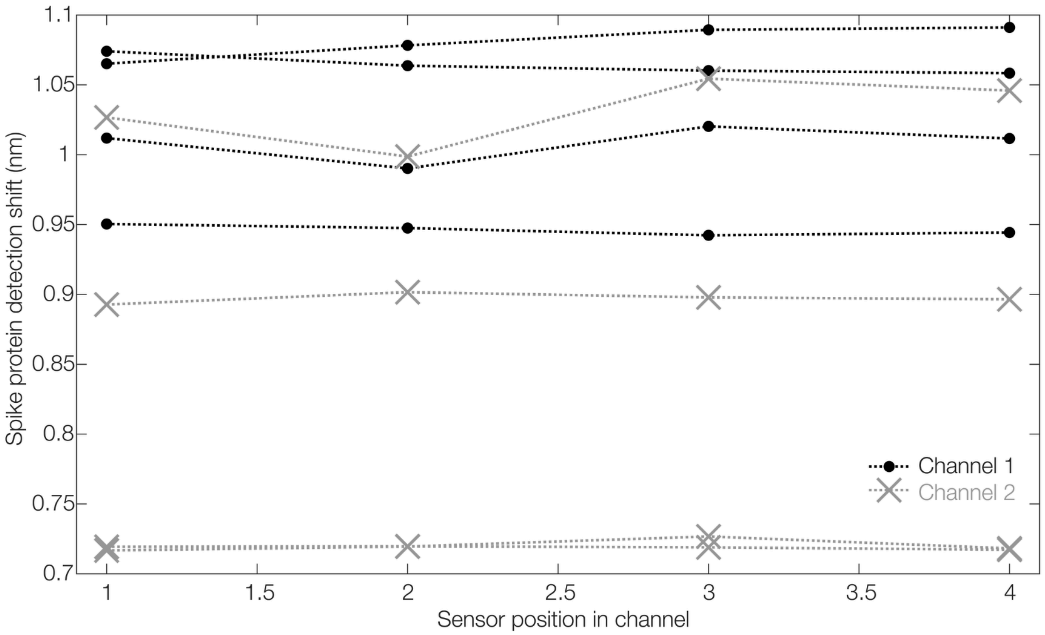

**Fig. S8.** Spike protein detection signal vs. sensor position along the length of the microfluidic channel, for the 20 μg/mL spike protein assays using Protein A-based functionalization presented in Fig. 5. Sensor position is denoted as 1, 2, 3, 4 where 1 is the most upstream sensor and 4 is the most downstream sensor, with approximately 800 μm channel length between sensors 1 and 4. Eight datasets are plotted, representing the sensors in the 2 microfluidic channels in each of 4 trials. We do not observe any obvious consistent relationship between sensor signal and position along the length of the microfluidic channel.

**Table S18.** Pearson correlation coefficients (r) and p-values (p) testing the null hypothesis that there is no relationship between the sensor position along the length of the channel and the detected signal. Only 1 out of 8 channel-replicates shows a significant correlation ($p < 0.05$).

| **Trial** | **1** | | **2** | | **3** | | **4** | |
|---|---|---|---|---|---|---|---|---|
| **Channel** | **1** | **2** | **1** | **2** | **1** | **2** | **1** | **2** |
| **r** | 0.96 | 0.59 | 0.29 | 0.33 | -0.85 | -0.84 | -0.93 | 0.27 |
| **p** | 0.04 | 0.41 | 0.71 | 0.66 | 0.15 | 0.16 | 0.07 | 0.73 |

#### S6. Mass transport estimates

Mass transport calculations were performed to obtain order-of-magnitude estimates of the amount of capture antibody transported to the photonic sensor surface by the spotting- and flow-mediated antibody immobilization protocols used in this work, as well as to quantify the relative magnitudes of convective and diffusive transport during in-flow spike protein detection. This analysis was based on models described by Squires et al. (24) for biomolecule transport in surface capture-based biosensor systems. In the spotting-mediated antibody immobilization protocol, we assumed that mass transport of antibodies to the sensor surface was driven solely by diffusion, while in the flow-mediated protocol, it was driven by diffusion and convection. For simplicity, reaction kinetics were not considered in this analysis and it was assumed that all antibody molecules transported to the surface were instantaneously immobilized.

#### S6.1. Spotting-mediated antibody immobilization

The diffusivity ($D$) of the capture antibody in the spotting buffer (10% glycerol, 0.005% Triton X-100 in PBS) was first calculated according to the Stokes-Einstein-Smoluchowski equation (25,26) (Eq. S2).

$$D = k_B T / 6\pi\mu R_h \quad \text{Eq. S2}$$

Here, $k_B$ is the Boltzmann constant, $T$ is the temperature (293.15 K), $\mu$ is the dynamic viscosity of the spotting buffer, and $R_h$ is the hydrodynamic radius of the antibody (assumed to be 5.5 nm (27–29)). Based on data reported by Takamura et al. (30), the dynamic viscosity of the spotting buffer was approximated as that of an aqueous brine solution (brine salinity II) containing 10% glycerol, giving $1.465 \times 10^{-3}$ Pa·s. This yielded an approximate diffusivity of $2.66 \times 10^{-11}$ m$^2$/s.

Next, we estimated the mass of antibody transported to the sensor surface during the 1-hour incubation time (24). The diffusive flux of antibodies is given by $j_D = - D\nabla c$, where $\nabla c$ is the concentration gradient of antibodies in the droplet. For scaling purposes, this was approximated as $\nabla c \sim c_0/\delta$, where $c_0$ is the initial antibody concentration in the spotting solution (500 µg/mL) and $\delta$ is the depletion zone thickness, which scales like $\delta \sim \sqrt{Dt}$. The diffusive collection rate, $J_D$, which is the mass of antibodies transported to the sensor surface per unit time, can thus be approximated as $J_D = j_D A_{spot} \sim A_{spot} c_0 \sqrt{D/t}$, where $A_{spot}$ is the surface area of the droplet. This was measured to be roughly 0.29 cm$^2$, based on micrographs captured during antibody spotting. Integrating $J_D$ over time gives the total mass of antibody transported to the sensor surface, $m = 2A_{spot} c_0 \sqrt{Dt}$. For a 1-hour incubation time, this gives ~9.0 µg of antibody.

We then compared this to the theoretical mass of antibody required to fully saturate the sensor surface. We assumed that the maximum density of antibody molecules immobilized on the sensor surface, $b_m$, is $2 \times 10^{12}$ molecules/cm$^2$ (24). This value represents the maximum active site density that can be achieved for a near monolayer of Fab fragments immobilized on a surface in an oriented manner. As our functionalization protocol employs whole antibodies immobilized in random orientations, this value of $b_m$ represents an impossible upper limit; the active site density that can realistically be achieved is likely an order of magnitude or more lower (24). Using this upper limit value of $b_m$, the maximum mass of antibodies that can be immobilized over the spotted area is given by $m_{max} = b_m A_{spot} M_{Ab} / N_A = \mathbf{0.14}\ \boldsymbol{\mu g}$, where $M_{Ab}$ is the molar mass of the antibody (~150,000 g/mol) and $N_A$ is Avogadro's number. This is ~64× smaller than the 9.0 µg of antibodies that can be delivered to the surface by diffusion in this protocol. Hence, we can conclude that the antibody concentration and incubation time employed in this spotting-mediated immobilization protocol are in excess of what is needed to saturate the surface with receptors.

Finally, we extended this analysis to the case of piezoelectric inkjet-based antibody spotting, whereby 0.1–1 nanoliter-scale spots with diameters on the order of 100 µm can be deposited on individual ring resonator sensors to facilitate precise multiplexed functionalization (18,31,32). Considering a spot volume of 1 nL and diameter of 150 µm, performing the same analysis reveals that ~0.87 ng of antibody can be transported to the sensor surface in the spotting region during the first 1.5 minutes of incubation. This is 9.8× greater than the ~0.088 ng of antibody required to fully saturate the surface in the spotting region. After 1 hour of incubation, roughly ~5.5 ng of antibody can be transported to the sensor surface in the spotting region, which is ~62× greater than that required for surface saturation.

#### S6.2. Flow-mediated antibody immobilization

The model system geometry for this protocol consists of a rectangular fluidic channel of height $H$ = 500 µm, width $W_c$ = 300 µm, length $L$ = 6772 µm, and cross-sectional area $A_{c,cross} = HW_c$ = $1.5 \times 10^{-3}$ cm$^2$. The surface area of the photonic sensor enclosed by the fluidic channel is given by $A_{c,surf} = LW_c$ = $2.0 \times 10^{-2}$ cm$^2$. The ratio of the channel length to height is defined as $\lambda = L/H$ = 13.5. In this system, a capture antibody solution of concentration $c_0$ = 20 µg/mL in PBS ($\mu$ = $1.02 \times 10^{-3}$ Pa·s (33)) was delivered to the sensor at a flow rate of $Q$ = 20 µL/min for 25 min. The antibody diffusivity in the running buffer (PBS) was calculated according to the Stokes-Einstein-Smoluchowski equation (25,26) (Eq. S2). This yielded a diffusivity of $3.82 \times 10^{-11}$ m$^2$/s.

To compare the relative effects of diffusion and convection on mass transport in this system, we first calculated the Peclet number, $Pe_H = Q/DW_c$, which is the ratio of the diffusive and convective time scales (24). This gave a value of $Pe_H$ = $2.9 \times 10^4$, indicating that mass transport in this system is diffusion-limited. The shear Peclet number, $Pe_S$, which depends on the sensor shear rate and sensor length, was calculated as $Pe_S = 6\lambda^2 Pe_H$ = 3.2 × 10$^7$, indicating that the depletion zone is thin compared to the sensor length. For this scenario where $Pe_H$, $Pe_S \gg 1$, the flux of antibodies to the sensor surface can be estimated as $J_D \approx Dc_0 W_s Pe_S^{1/3}$ = 7.3 × 10$^{-5}$ µg/s. Multiplying this by the 25-minute flow time gives a total mass of ~0.11 µg of antibody delivered to the sensor surface. This is nearly 11× the theoretical maximum loading of antibodies on the sensor surface, which is given by $m_{max} = b_m A_{c,surf} M_{Ab} / N_A$ = 0.010 µg. As the actual achievable surface loading of antibodies is likely an order of magnitude or more smaller than $m_{max}$, the conditions employed in this flow-mediated antibody immobilization protocol should deliver ample excess antibody to the sensor to achieve saturation.

##### S6.3. In-flow analyte detection

The diffusivity of spike protein was calculated according to the Stokes-Einstein-Smoluchowski equation using the fluid properties of PBS (25,26) (Eq. S2) and assuming a hydrodynamic radius of 3.15 nm, which was estimated based on the number of amino acid residues (681 residues) (34). This yielded a diffusivity of 6.67 × 10$^{-11}$ m/s. To compare the relative effects of diffusion and convection on mass transport in this system, we calculated the Peclet number, considering a flow rate of 30 µL/min, which gave a value of $Pe_H$ = 2.5 × 10$^4$, indicating that convection is dominant and mass transport in this system is diffusion-limited (24). As $Pe_H \gg 1$, we may assume that convection is adequately large that variations in flow velocity across the channel width and height have a small effect on the dominant mass transport regimes across the channel.

##### S6.4. Limitations

These calculations represent order-of-magnitude estimates of mass transport phenomena occurring in our biosensor system; detailed models of these phenomena are outside the scope of this work. Simplifying assumptions were made to enable this analysis where empirical data were not available, which may limit the accuracy of the calculated values. For example, in the antibody spotting model system, the geometry of the antibody spot was simplified from a fluid cap to a cylinder, while additional mechanisms of mass transport in the fluid other than diffusion, such as capillary or Marangoni flows (35), were not accounted for. These factors could affect the uniformity of antibody capture across the spotted area, as well as transport kinetics. In the flow-mediated antibody immobilization model, adsorption of antibodies to the microfluidic tubing upstream of the sensor and absorption of antibodies by the PDMS gasket were not accounted for (36). These phenomena may be expected to partially deplete antibodies from the functionalization solution, effectively decreasing the concentration and slowing mass transport. Additionally, we made the assumption that all antibodies are immediately immobilized upon encountering the surface, which discounts reaction kinetics between the antibodies and PDA film or Protein A layer (24). We also made the assumption that antibodies are immobilized at the maximum possible surface density, $b_m$ = 2 × 10$^{12}$ molecules/cm$^2$ (24), as described in the sections above, and did not account for conformational changes and interactions between proteins on the surface (37). In reality, the rate at which antibodies are immobilized to the sensor surface may be limited by these factors.

#### S7. Summary of sensor-level, channel-level, and assay-level CVs

**Table S19.** Summary of reported experimental CVs in our manuscript, along with the factors expected to impact them. Factors that may be expected to vary on multiple levels (e.g., some variation at the sensor-level and additional higher variation at the chip/assay level) are reported in multiple columns.

| | Metric | Single-channel intra-assay CV | Single-channel intra-assay CV contributing factors (sensor-specific) | 2-channel intra-assay CV | 2-channel intra-assay CV: additional contributing factors (channel-specific) | Inter-assay CV | Inter-assay CV: additional contributing factors (assay-specific) |
|---|---|---|---|---|---|---|---|
| Intrinsic performance | Stability $\sigma_{\Delta\lambda}$ | 26 ± 30% for ch. 1<br>28 ± 18% for ch. 2 | • Surface cleanliness<br>• Waveguide wetting | 61 ± 30% | • Bubbles<br>• Surface etching or material adsorption<br>• Waveguide wetting<br>• Reagent depletion/ contamination<br>• Flow rate stability<br>• Channel alignment | 83.1% | • Optical and thermal noise<br>• Surface cleanliness<br>• Variability in prepared fluids |
| | Baseline drift rate ($\Delta\lambda/\Delta t$) | 40 ± 50% for ch. 1<br>30 ± 20% for ch. 2 | • Surface cleanliness<br>• Waveguide wetting | 160 ± 180% | • Bubbles<br>• Surface etching or material adsorption<br>• Waveguide wetting<br>• Reagent depletion/ contamination | 83.1% | • Optical and thermal noise<br>• Surface cleanliness<br>• Variability in prepared fluids |

| | Metric | Single-channel intra-assay CV | Single-channel intra-assay CV contributing factors (sensor-specific) | 2-channel intra-assay CV | 2-channel intra-assay CV: additional contributing factors (channel-specific) | Inter-assay CV | Inter-assay CV: additional contributing factors (assay-specific) |
|---|---|---|---|---|---|---|---|
| | | | | | • Flow rate stability | | |
| | Bulk RI sensitivity $S_{bulk}$ | 0.16 ± 0.13% for ch. 1<br>1 ± 2% for ch. 2 | • Surface cleanliness<br>• Fabrication variation<br>• Waveguide wetting | 1.4 ± 1.4% | • Waveguide wetting<br>• Bubbles<br>• Reagent depletion/ contamination<br>• Flow rate stability<br>• Channel alignment | 4.2% | • Optical and thermal noise<br>• Surface cleanliness<br>• Fabrication variation<br>• Oxide-open variation<br>• Variability in prepared fluids |
| | System limit of detection sLoD | 26 ± 30% for ch. 1<br>28 ± 18% for ch. 2 | All factors above for $S_{bulk}$ and $\sigma_{\Delta\lambda}$ | 60 ± 30% | All factors above for $S_{bulk}$ and $\sigma_{\Delta\lambda}$ | 83.8% | All factors above for $S_{bulk}$ and $\sigma_{\Delta\lambda}$ |
| Analyte-detection performance (Protein A/flow, 20 µg/mL) | Pre-functionalization baseline drift rate | 31 ± 7% for ch. 1<br>500 ± 800% for ch. 2 | • Surface cleanliness<br>• Waveguide wetting | 300 ± 300% | • Bubbles<br>• Surface etching or material adsorption<br>• Waveguide wetting<br>• Reagent depletion/ contamination<br>• Flow rate stability<br>• Functionalization stability | 117.7% | • Optical and thermal noise<br>• Surface cleanliness<br>• Variability in prepared fluids<br>• Surface cleanliness |
| | Functionalization (capture antibody attachment) peak shift $\Delta\lambda_{func}$ | 1.2 ± 0.7% for ch. 1<br>3 ± 4% for ch. 2 | • Surface cleanliness<br>• Fabrication variation<br>• Waveguide wetting<br>• In-channel depletion | 14 ± 14% | • Bubbles<br>• Reagent depletion or contamination<br>• Flow rate stability<br>• Fluidic channel alignment<br>• Waveguide wetting | 22.3% | • Bioreceptor solution factors (e.g., aggregates, pH)<br>• Bioreceptor stability<br>• Oxide-open variation<br>• Surface cleanliness |
| | Analyte-binding peak shift $\Delta\lambda_{binding}$ | 0.9 ± 0.4% for ch. 1<br>0.9 ± 1.0% for ch. 2 | • Surface cleanliness<br>• Fabrication variation<br>• Waveguide wetting<br>• In-channel depletion | 11 ± 7% | • Bubbles<br>• Bioreceptor orientation and density<br>• Reagent depletion or contamination<br>• Flow rate stability<br>• Fluidic channel alignment<br>• Waveguide wetting<br>• Nonspecific and off-target binding | 11.1% | • Bioreceptor solution factors (e.g., aggregates, pH)<br>• Bioreceptor stability<br>• Oxide-open variation<br>• Surface cleanliness<br>• Reagent variability |
| | Nonspecific binding peak shift $\Delta\lambda_{nonspec}$ | 2.7 ± 2.0% for ch. 1<br>10 ± 8% for ch. 2 | • Surface cleanliness<br>• Fabrication variation<br>• Waveguide wetting<br>• In-channel depletion | 24 ± 20% | • Reagent depletion or contamination<br>• Bioreceptor orientation and density<br>• Nonspecific and off-target binding<br>• Waveguide wetting | 76.5% | • Reagent variability<br>• Bioreceptor stability |
| | Post-assay drift rate | 2.8 ± 1.2% for ch. 1<br>5 ± 7% for ch. 2 | • Surface cleanliness<br>• Waveguide wetting | 24 ± 20% | • Bubbles<br>• Surface etching or material adsorption<br>• Waveguide wetting<br>• Flow rate stability<br>• Functionalization and binding interaction stability<br>• Bound analyte signal | 16.6% | • Bound analyte signal<br>• Reagent variability |

| | Metric | Single-channel intra-assay CV | Single-channel intra-assay CV contributing factors (sensor-specific) | 2-channel intra-assay CV | 2-channel intra-assay CV: additional contributing factors (channel-specific) | Inter-assay CV | Inter-assay CV: additional contributing factors (assay-specific) |
|---|---|---|---|---|---|---|---|
| Analyte-detection performance (PDA/spotting, 1 μg/mL) | Initial baseline drift rate (post-functionalization, pre-assay) | 37.8 ± 15.8% for ch. 1<br>133.0 ± 100.1% for ch. 2 | • Surface cleanliness<br>• Waveguide wetting<br>• Functionalization stability | 112.3 ± 71.5% | • Bubbles<br>• Surface etching or material adsorption<br>• Reagent depletion/ contamination<br>• Flow rate stability<br>• Biorecetor orientation and density<br>• Functionalization stability | 1333.4% | • Optical and thermal noise<br>• Sample preparation<br>• Reagent variability<br>• Biorecetor orientation and density<br>• Functionalization stability |
| | Analyte-binding peak shift $\Delta\lambda_{binding}$ | 5.4 ± 1.9% for ch. 1<br>2.8 ± 0.7% for ch. 2 | • Biorecetor orientation and density<br>• Functionalization stability<br>• Fabrication variation<br>• Waveguide wetting<br>• In-channel depletion | 4.7 ± 0.6% | • Bubbles<br>• Reagent depletion or contamination<br>• Flow rate stability<br>• Fluidic channel alignment<br>• Biorecetor orientation and density<br>• Functionalization stability<br>• Waveguide wetting<br>• Nonspecific and off-target binding | 18.1% | • Sample preparation<br>• Reagent variability<br>• Biorecetor orientation and density<br>• Functionalization stability<br>• Fabrication variation |
| | Nonspecific binding peak shift $\Delta\lambda_{nonspec}$ | 9.0 ± 7.9% for ch. 1<br>2.1 ± 1.0% for ch. 2 | • Surface cleanliness<br>• Fabrication variation<br>• Waveguide wetting<br>• In-channel depletion | 6.7 ± 5.5% | • Bubbles<br>• Reagent depletion or contamination<br>• Functionalization stability<br>• Nonspecific and off-target binding<br>• Waveguide wetting | 34% | • Sample preparation<br>• Reagent variability<br>• Functionalization stability<br>• Fabrication variation |
| | Post-assay drift rate | 26.0 ± 15.0% for ch. 1<br>18.2 ± 12.5% for ch. 2 | • Surface cleanliness<br>• Waveguide wetting<br>• Functionalization stability | 25.5 ± 4.1% | • Bubbles<br>• Surface etching or material adsorption<br>• Reagent depletion/ contamination<br>• Flow rate stability<br>• Biorecetor orientation and density<br>• Functionalization stability | 10% | • Optical and thermal noise<br>• Bound analyte signal<br>• Sample preparation<br>• Reagent variability |
| Analyte-detection performance (PDA/flow, 1 μg/mL) | Functionalization (capture antibody attachment) peak shift $\Delta\lambda_{func}$ | 10.7 ± 12.1% for ch. 1<br>8.1 ± 4.6% for ch. 2 | • Surface cleanliness<br>• Waveguide wetting<br>• In-channel depletion | 51.7 ± 42.1% | • Bubbles<br>• Reagent depletion/ contamination<br>• Flow rate stability<br>• Fluidic channel alignment<br>• Waveguide wetting | 76% | • Sample preparation<br>• Reagent variability<br>• Biorecetir stability<br>• Surface cleanliness |
| | Initial baseline drift rate (post-functionalization, pre-assay) | 133.0 ± 100.1% for ch. 1<br>205.1 ± 47.9% for ch. 2 | • Surface cleanliness<br>• Waveguide wetting<br>• Functionalization stability | 271.4 ± 279.9% | • Bubbles<br>• Surface etching or material adsorption<br>• Reagent depletion/ contamination<br>• Flow rate stability<br>• Biorecetor orientation and density<br>• Functionalization stability | 120.6% | • Optical and thermal noise<br>• Sample preparation<br>• Reagent variability<br>• Biorecetor orientation and density<br>• Functionalization stability |
| | Analyte-binding peak shift $\Delta\lambda_{binding}$ | 37.8 ± 39.7% for ch. 1<br>17.6 ± 8.0% for ch. 2 | • Biorecetor orientation and density<br>• Functionalization stability<br>• Fabrication variation<br>• Waveguide wetting<br>• In-channel depletion | 642.7 ± 999.4% | • Bubbles<br>• Reagent depletion or contamination<br>• Flow rate stability<br>• Fluidic channel alignment<br>• Biorecetor orientation and density<br>• Functionalization stability<br>• Waveguide wetting<br>• Nonspecific and off-target binding | 125% | • Sample preparation<br>• Reagent variability<br>• Biorecetor orientation and density<br>• Functionalization stability<br>• Fabrication variation |

| | Metric | Single-channel intra-assay CV | Single-channel intra-assay CV contributing factors (sensor-specific) | 2-channel intra-assay CV | 2-channel intra-assay CV: additional contributing factors (channel-specific) | Inter-assay CV | Inter-assay CV: additional contributing factors (assay-specific) |
|---|---|---|---|---|---|---|---|
| | Nonspecific binding peak shift $\Delta\lambda_{nonspec}$ | 7.9 ± 2.5% for ch. 1<br>3.7 ± 1.9% for ch. 2 | • Surface cleanliness<br>• Fabrication variation<br>• Waveguide wetting<br>• In-channel depletion | 14.4 ± 11.5% | • Bubbles<br>• Reagent depletion or contamination<br>• Functionalization stability<br>• Nonspecific and off-target binding<br>• Waveguide wetting | 39% | • Sample preparation<br>• Reagent variability<br>• Functionalization stability<br>• Fabrication variation |
| | Post-assay drift rate | 15.7 ± 12.4% for ch. 1<br>9.5 ± 11.2% for ch. 2 | • Surface cleanliness<br>• Waveguide wetting<br>• Functionalization stability | 25.3 ± 9.3% | • Bubbles<br>• Surface etching or material adsorption<br>• Reagent depletion/ contamination<br>• Flow rate stability<br>• Biorecetor orientation and density<br>• Functionalization stability | 41% | • Optical and thermal noise<br>• Bound analyte signal<br>• Sample preparation<br>• Reagent variability |

### S8. Baseline-correction fit functions

**Table S20.** Summary of the baseline correction fit parameters (for fit function $y_{fit} = a \cdot \left(1 - e^{b(t-c)}\right) + d$) as well as the expected correction shift over a 20-min period at the end of the baseline correction period ($\Delta y_{fit,\,0-20min}$) and 100 minutes later ($\Delta y_{fit,\,100-120min}$). Using a 20-min period immediately following the baseline correction period is expected to overestimate the baseline-correction shift for later binding steps, since the functions tend to level off over time. Estimated correction shifts are overall small, with all shifts < 39 pm in the period immediately following baseline correction, and < 11 pm 100 minutes later.

| Assay series | Trial | Sensor | a (nm) | b (1/min) | c (min) | d (nm) | $\Delta y_{fit,\,0-20min}$ (nm) | $\Delta y_{fit,\,100-120min}$ (nm) |
|---|---|---|---|---|---|---|---|---|
| Analyte-detection performance (Protein A/flow, 20 μg/mL) | 1 | 1 | 12.1 | -1 | 500 | -11.4 | 0.000E+00 | 0.000E+00 |
| | | 2 | 10.3 | -1 | 500 | -9.6 | 0.000E+00 | 0.000E+00 |
| | | 3 | -7.29 | -1 | 500 | 8 | 0.000E+00 | 0.000E+00 |
| | | 4 | 10.2 | -1 | 500 | -9.44 | 0.000E+00 | 0.000E+00 |
| | | 5 | 9.22 | -1 | 500 | -8.55 | 0.000E+00 | 0.000E+00 |
| | | 6 | 7.98 | -1 | 500 | -7.25 | 0.000E+00 | 0.000E+00 |
| | | 7 | 0.751 | -1 | 500 | -0.081 | 0.000E+00 | 0.000E+00 |
| | | 8 | 7.17 | -1 | 500 | -6.44 | 0.000E+00 | 0.000E+00 |
| | 2 | 1 | 10.3 | -1 | 500 | -9.46 | 0.000E+00 | 0.000E+00 |
| | | 2 | 0.211 | -1 | 500 | 0.55 | 0.000E+00 | 0.000E+00 |
| | | 3 | 13.3 | -1 | 500 | -12.5 | 0.000E+00 | 0.000E+00 |
| | | 4 | 0.684 | -1 | 500 | 0.084 | 0.000E+00 | 0.000E+00 |
| | | 5 | 12.1 | -1 | 500 | -11.3 | 0.000E+00 | 0.000E+00 |
| | | 6 | -0.00622 | -1 | 500 | 0.774 | 0.000E+00 | 0.000E+00 |
| | | 7 | 9.78 | -1 | 500 | -8.96 | 0.000E+00 | 0.000E+00 |
| | | 8 | 0.42 | -1 | 500 | 0.351 | 0.000E+00 | 0.000E+00 |
| | 3 | 1 | 0.168 | -0.626 | 262 | 0.607 | 8.528E-08 | 0.000E+00 |
| | | 2 | 12.6 | -0.0947 | 214 | -11.6 | 1.341E-02 | 1.035E-06 |
| | | 3 | 0.184 | -0.713 | 263 | 0.586 | 2.301E-08 | 0.000E+00 |
| | | 4 | 12.6 | -0.0938 | 214 | -11.6 | 1.373E-02 | 1.159E-06 |
| | | 5 | -0.161 | -0.451 | 259 | 0.932 | -1.555E-06 | 0.000E+00 |

| Assay series | Trial | Sensor | a (nm) | b (1/min) | c (min) | d (nm) | $\Delta y_{fit,\,0-20min}$ (nm) | $\Delta y_{fit,\,100-120min}$ (nm) |
|---|---|---|---|---|---|---|---|---|
| | | 6 | 12.4 | -0.0904 | 212 | -11.5 | 1.396E-02 | 1.654E-06 |
| | | 7 | 0.15 | -0.523 | 260 | 0.618 | 3.695E-07 | 0.000E+00 |
| | | 8 | 12.5 | -0.0952 | 215 | -11.5 | 1.297E-02 | 9.521E-07 |
| | 4 | 1 | 0.185 | -0.478 | 84.9 | 0.597 | 7.060E-07 | 0.000E+00 |
| | | 2 | 0.744 | -0.117 | 51.4 | 0.0631 | 6.498E-04 | 5.663E-09 |
| | | 3 | -0.0998 | -0.67 | 88.1 | 0.88 | -2.168E-08 | 0.000E+00 |
| | | 4 | 1.03 | -0.118 | 49.5 | -0.221 | 6.676E-04 | 5.127E-09 |
| | | 5 | -0.203 | -0.0816 | 33.1 | 0.983 | -2.835E-04 | -8.136E-08 |
| | | 6 | 0.546 | -0.161 | 64.3 | 0.257 | 2.847E-04 | 2.867E-11 |
| | | 7 | -0.0265 | -1.29 | 91.9 | 0.808 | -4.841E-13 | 0.000E+00 |
| | | 8 | 0.643 | -0.152 | 60 | 0.157 | 2.633E-04 | 6.691E-11 |
| Analyte-detection performance (PDA/spotting, 1 µg/mL) | 1 | 1 | -12.8 | -0.0391 | -87 | 12.2 | -1.570E-02 | -3.161E-04 |
| | | 2 | -16 | -0.0353 | -97.1 | 15.3 | -2.294E-02 | -6.702E-04 |
| | | 3 | -12.7 | -0.0448 | -72.1 | 12.1 | -1.345E-02 | -1.522E-04 |
| | | 4 | -16.2 | -0.0135 | -271 | 15.4 | -3.876E-02 | -1.002E-02 |
| | | 5 | -12.3 | -0.0508 | -60.3 | 11.7 | -1.094E-02 | -6.790E-05 |
| | | 6 | -15.8 | -0.0247 | -147 | 15.1 | -2.924E-02 | -2.464E-03 |
| | | 7 | -13.1 | -0.0401 | -84.1 | 12.5 | -1.558E-02 | -2.829E-04 |
| | | 8 | -16.2 | -0.0349 | -98.4 | 15.5 | -2.383E-02 | -7.298E-04 |
| | 2 | 1 | -7.94 | -0.0523 | -58.6 | 7.25 | -6.194E-03 | -3.329E-05 |
| | | 2 | -4.84 | -0.00584 | -549 | 3.99 | -1.437E-02 | -8.013E-03 |
| | | 3 | -8.18 | -0.0456 | -71.6 | 7.48 | -7.638E-03 | -7.968E-05 |
| | | 4 | -5.35 | -0.017 | -210 | 4.6 | -1.320E-02 | -2.404E-03 |
| | | 5 | -11.8 | -0.049 | -61.5 | 11.1 | -1.183E-02 | -8.847E-05 |
| | | 6 | -5.96 | -0.00842 | -405 | 5.15 | -1.687E-02 | -7.267E-03 |
| | | 7 | -7.83 | -0.0407 | -84.3 | 7.12 | -8.155E-03 | -1.390E-04 |
| | | 8 | -8.03 | -0.00763 | -443 | 7.15 | -2.268E-02 | -1.058E-02 |
| | 3 | 1 | -0.658 | -0.715 | 36.5 | 0.585 | -1.118E-10 | 0.000E+00 |
| | | 2 | 2.06 | -0.00888 | -376 | -2.07 | 6.510E-03 | 2.680E-03 |
| | | 3 | -0.0796 | -1.07 | 40.5 | 0.0206 | -1.255E-14 | 0.000E+00 |
| | | 4 | 3.2 | -0.00888 | -385 | -3.19 | 9.314E-03 | 3.831E-03 |
| | | 5 | -0.868 | -0.543 | 33.8 | 0.807 | -7.467E-09 | 0.000E+00 |
| | | 6 | 2.39 | -0.00728 | -448 | -2.4 | 7.562E-03 | 3.650E-03 |
| | | 7 | -1.26 | -0.421 | 30.6 | 1.2 | -1.806E-07 | 0.000E+00 |
| | | 8 | 1.72 | -0.0269 | -126 | -1.77 | 3.874E-03 | 2.625E-04 |

| Assay series | Trial | Sensor | a (nm) | b (1/min) | c (min) | d (nm) | $\Delta y_{fit,\,0-20min}$ (nm) | $\Delta y_{fit,\,100-120min}$ (nm) |
|---|---|---|---|---|---|---|---|---|
| Analyte-detection performance (PDA/flow, 1 µg/mL) | 1 | 1 | -4.21 | -0.104 | 160 | 6.97 | -1.063E-03 | -3.314E-08 |
| | | 2 | -1.78 | -0.146 | 176 | 3.56 | -1.815E-04 | -8.666E-11 |
| | | 3 | -4.19 | -0.122 | 168 | 7.05 | -6.670E-04 | -3.310E-09 |
| | | 4 | -3.92 | -0.0769 | 144 | 6.37 | -2.075E-03 | -9.537E-07 |
| | | 5 | -4.36 | -0.112 | 164 | 7.21 | -8.690E-04 | -1.180E-08 |
| | | 6 | -3.43 | -0.11 | 164 | 5.76 | -8.185E-04 | -1.347E-08 |
| | | 7 | -4.32 | -0.107 | 162 | 7.22 | -9.760E-04 | -2.171E-08 |
| | | 8 | -1.74 | -0.131 | 172 | 3.43 | -2.452E-04 | -5.180E-10 |
| | 2 | 1 | -1.45 | -0.167 | 143 | 3.52 | -1.573E-04 | -8.904E-12 |
| | | 2 | -4.09 | -0.115 | 128 | 5.47 | -1.395E-03 | -1.439E-08 |
| | | 3 | -1.63 | -0.0397 | 45.3 | 3.77 | -2.159E-03 | -4.075E-05 |
| | | 4 | -3.35 | -0.0488 | 68.5 | 4.78 | -3.980E-03 | -3.036E-05 |
| | | 5 | -2.19 | -0.0956 | 119 | 4.03 | -1.056E-03 | -7.426E-08 |
| | | 6 | -4.94 | -0.0917 | 117 | 6.13 | -2.687E-03 | -2.795E-07 |
| | | 7 | -1.26 | -0.0319 | 17.5 | 3.46 | -1.926E-03 | -7.895E-05 |
| | | 8 | -4.86 | -0.0919 | 117 | 6.03 | -2.564E-03 | -2.609E-07 |
| | 3 | 1 | -8.45 | -0.0597 | 162 | 10.7 | -8.691E-03 | -2.221E-05 |
| | | 2 | -6.07 | -0.105 | 197 | 8.5 | -2.224E-03 | -6.336E-08 |
| | | 3 | -4.45 | -0.0773 | 179 | 6.94 | -2.839E-03 | -1.248E-06 |
| | | 4 | -6.47 | -0.0888 | 188 | 8.89 | -3.310E-03 | -4.618E-07 |
| | | 5 | -4.84 | -0.0793 | 181 | 7.43 | -3.074E-03 | -1.110E-06 |
| | | 6 | -6.94 | -0.096 | 193 | 9.42 | -3.170E-03 | -2.146E-07 |
| | | 7 | -9 | -0.0564 | 157 | 11.5 | -9.973E-03 | -3.548E-05 |
| | | 8 | -10.1 | -0.0768 | 180 | 12.5 | -7.316E-03 | -3.393E-06 |
| (PrA/flow, 1 µg/mL) | 1 | 1 | -0.155 | -0.0549 | 123 | 3.07 | -2.657E-02 | -1.098E-04 |
| | | 2 | -0.161 | -0.0604 | 123 | 2.85 | -2.525E-02 | -6.032E-05 |
| | | 3 | -0.149 | -0.0585 | 123 | 3.08 | -2.412E-02 | -6.965E-05 |
| | | 4 | -0.156 | -0.0639 | 123 | 2.86 | -2.296E-02 | -3.844E-05 |
| | | 5 | -0.149 | -0.0588 | 123 | 3.1 | -2.399E-02 | -6.684E-05 |
| | | 6 | -0.16 | -0.0613 | 123 | 2.85 | -2.461E-02 | -5.381E-05 |
| | | 7 | -0.159 | -0.0508 | 123 | 3.09 | -2.892E-02 | -1.804E-04 |
| | | 8 | -0.159 | -0.0589 | 123 | 2.83 | -2.556E-02 | -7.069E-05 |

### S9. Demonstration of a sandwich immunoassay for IL-8 detection in complex media on polydopamine-functionalized SiP sensors

#### S9.1. Methods

##### S9.1.1. IL-8 detection assay methods

Sandwich immunoassays to detect interleukin 8 (IL-8) in simple and complex media were run using methods similar to those reported in section S2 for the simpler demonstration assays reported in this manuscript (SARS-CoV-2 spike protein in buffer). This assay included the following stages, again with 10-min running buffer rinse steps between each microfluidic assay stage:

1. PDA/spotting-mediated functionalization with IL-8 capture antibody
2. Blocking with either BSA in PBS or BSA in cell culture media containing 10% fetal bovine serum (FBS)
3. Integration with microfluidics
4. Initial rinse with regeneration buffer to remove loosely attached bioreceptors or blocking proteins
5. Nonspecific binding challenge with BSA and the sample matrix (either PBS or cell culture medium)
6. Detection of IL-8 in running buffer or in complex media (cell culture media)
7. Binding of biotinylated IL-8 detection antibody
8. Signal amplification using high-sensitivity streptavidin-HRP (SA-HRP)

A cross-sectional schematic visualizing the assay stages is presented in Figure S9.

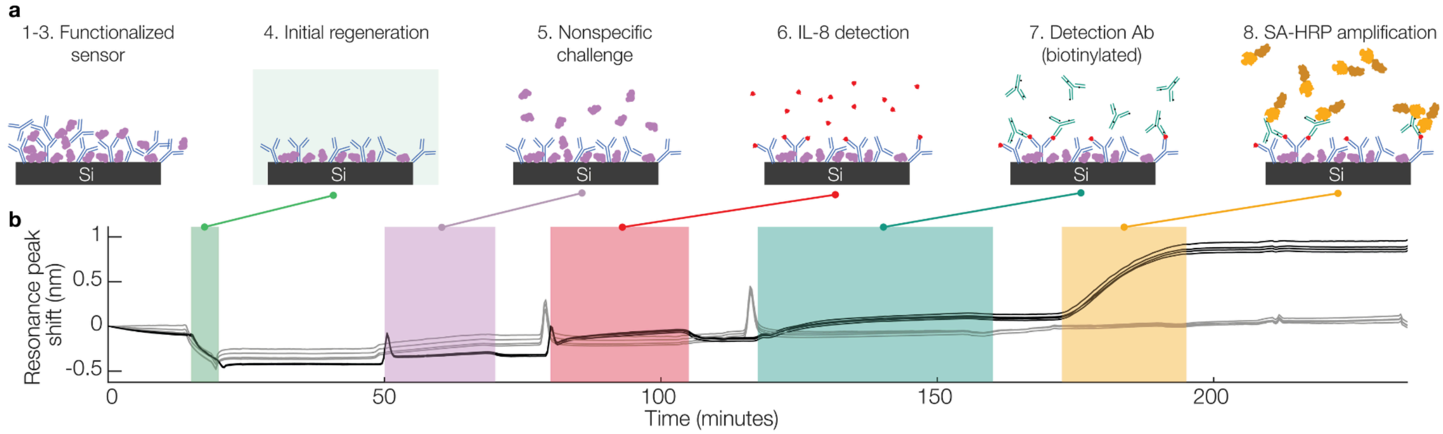

**Fig. S9.** Schematics showing the stages of the sandwich assay used to detect IL-8 as well as example sensorgrams. **(a)** cross-sectional diagrams showing each stage of the sandwich assay. **(b)** example sensorgrams for a sample containing IL-8 (black) and a negative control sample (grey) in simple buffer solution.

The specific reagents used for this detection assay included: IL-8 capture antibody (included in ELISA kit R&D Systems DY208-05, lots ASJ3822051, ASJ3824031); IL-8 (included in ELISA kit R&D Systems DY208-05, lots ASJ3822051, ASJ3824031); IL-8 detection antibody (R&D Systems BAF208, lots UM2322052, UM221031), high-sensitivity streptavidin-HRP (Pierce 21130 lot XJ361814); Dulbecco's Modified Eagle's Medium (Sigma-Aldrich D6421, lot RNBM3386); Fetal Bovine Serum (Gibco 12483020, lot 2357664RP); L-glutamine (Sigma-Aldrich G7513, lot RNBM3694); 100× Antibiotic-Antimycotic (Thermo Scientific 15240-062, lot 2585910). Complete cell culture medium (CCM) was prepared by mixing 44 mL DMEM F/12, 5 mL FBS, 0.5 mL 200 mM L-glutamine, and 0.5 mL 100× Antibiotic-Antimycotic. This complete medium was stored at 4℃ for a maximum of one month and subsequently used for the sample matrix and blocking solution preparation. For these assays, an initial rinse with an acidic sensor regeneration buffer was used to remove loosely attached functionalization and blocking reagents from the sensor surface prior to the binding assay. This regeneration buffer was prepared using ultrapure water from a Nanopure Diamond system and included final concentrations of 10 mM glycine (Sigma-Aldrich G8790-100G, lot SLCG2930) and 160 mM NaCl (Fisher S271-3, lot 166221), adjusted to pH 2.2 using 1M HCl stock (prepared using ultrapure water and HCl concentrate: Sigma-Aldrich 258148-2.5L-GL, PCode: 4102988462, Source: MKCS8144). Other reagents (e.g., dopamine-HCl, Tris buffer, PBS) used the same stocks as described previously for the spike protein detection assays.

Sensors were coated with PDA and functionalized using a spotting-mediated functionalization process similar to that described in Section 2.5, using a 1-hour spotting incubation with a concentration of 240 µg/mL IL-8 capture antibody in spotting buffer (10% glycerol, 0.005% Triton X-100 in PBS). After rinsing in PBS, the sensors were immersed in blocking solution (20 mg/mL BSA in either PBS or CCM) for 1 hour, rinsed again in PBS, dried, and integrated with the microfluidic gasket (pre-treated by storing in vacuum overnight and exposure to plasma as previously described). Microfluidic channels were pre-wetted using 0.3 mM Triton X-100 in PBS before exposing the sensors to the automated binding assay protocol. The conditions tested are outlined in Table S21, and Table S22 presents the automated protocol for the IL-8 detection assay.

**Table S21.** Conditions tested for IL-8 detection assays.

| Condition | Name | IL-8 concentration (ng/mL) | Sample matrix | Blocking solution | Number of channel replicates tested (n) |
|---|---|---|---|---|---|

| | | | | | |
|---|---|---|---|---|---|
| 1a | PBS-BSA, BSA block | 3.125 | PBS-BSA running buffer | 20 mg/mL BSA in PBS | 5 |
| 1b | PBS-BSA, BSA block, neg control | 0 | PBS-BSA running buffer | 20 mg/mL BSA in PBS | 4 |
| 2a | CCM, BSA block | 3.125 | CCM | 20 mg/mL BSA in PBS | 2 |
| 2b | CCM, BSA block, neg control | 0 | CCM | 20 mg/mL BSA in PBS | 4 |
| 3a | CCM, BSA+CCM block | 3.125 | CCM | 20 mg/mL BSA in CCM | 2 |
| 3b | CCM, BSA+CCM block, neg control | 0 | CCM | 20 mg/mL BSA in CCM | 6 |

**Table S22.** Automated fluidic protocols for IL-8 detection assays.

| Assay step | Solution | Flow rate (µL/min) | Flow time (min) | Solution details |
|---|---|---|---|---|
| 1 | PBS-BSA | 30 | 60 | Assay running buffer: 0.1 mg/mL BSA in PBS |
| 2 | Regeneration | 30 | 5 | pH 2.2 Glycine-HCl buffer |
| 3 | PBS-BSA | 30 | 60 | Assay running buffer: 0.1 mg/mL BSA in PBS |
| 4 | Nonspecific challenge | 30 | 20 | 1 mg/mL BSA in PBS or 1 mg/mL BSA in CCM |
| 5 | PBS-BSA | 30 | 10 | Assay running buffer: 0.1 mg/mL BSA in PBS |
| 6 | IL-8 or negative control | 20-30 | 25 | 0 ng/mL or 3.125 ng/mL IL-8 in assay running buffer or in CCM |
| 7 | PBS-BSA | 30 | 10 | Assay running buffer: 0.1 mg/mL BSA in PBS |
| 8 | IL-8 detection antibody | 20 | 45 | 1 µg/mL detection antibody in assay running buffer |
| 9 | PBS-BSA | 30 | 10 | Assay running buffer: 0.1 mg/mL BSA in PBS |
| 10 | SA-HRP | 20 | 25 | 2 µg/mL high-sensitivity SA-HRP in assay running buffer |
| 11 | PBS-BSA | 30 | 30 | Assay running buffer: 0.1 mg/mL BSA in PBS |

##### S9.1.2. Assays incorporating reference sensors

To investigate the efficacy of performing reference-subtraction using sensors not functionalized with IL-8 capture antibody, we used a photonic chip architecture that included the same sensor architecture described in section S2.2, but with 12 sensors positioned in 3 distinct groups on the microfluidic chip, far enough apart that each group could separately be functionalized by manual spotting using a micropipette. Two groups of resonators (at the upstream and downstream sides of the chip) were each spotted with 1 µL IL-8 capture antibody solution (240 µg/mL IL-8 capture antibody in spotting buffer), and one group (in the middle of the chip) was spotted with 1 µL of a control solution (here, we used a simple blocking solution of 20 mg/mL BSA in spotting buffer but a framework for choosing an optimal reference sensor functionalization has been described previously by others (38)). The central sensors served as references for reference-subtraction in order to mitigate noise sources such as thermal drift, nonspecific binding, and bulk refractive index changes. Figure S10 highlights the sensor architecture used for this multiplexed functionalization process and shows the droplets spotted during functionalization as well as integration of the microfluidic gasket.

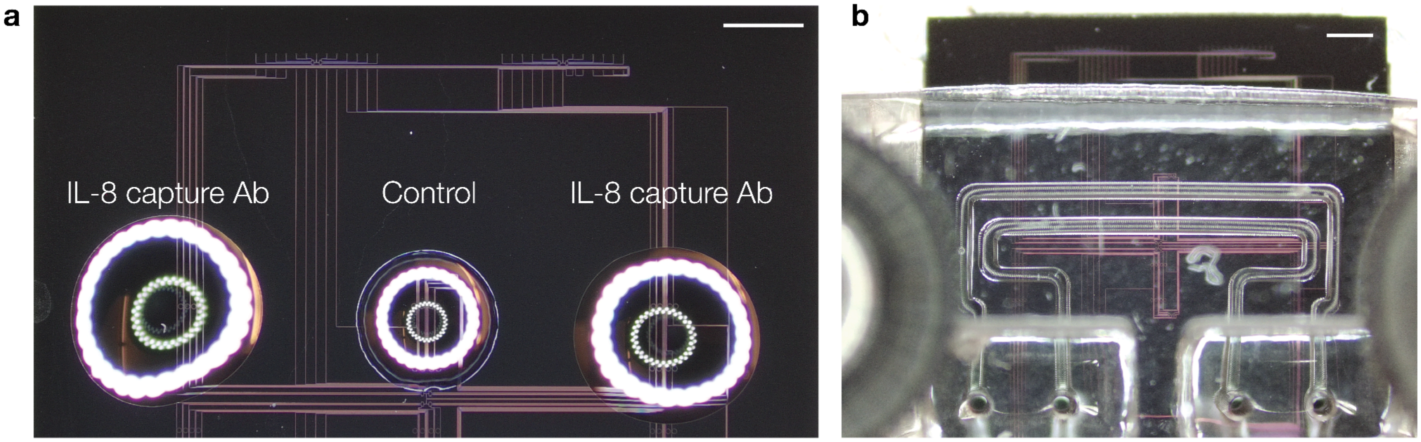

**Fig. S10**. Multiplexed SiP sensor architecture used for assays incorporating reference sensors. **(a)** Micrograph depicting the SiP chip during spotting, showing the 3 functionalization droplets (two with capture antibody (Ab) and one with BSA as a control) addressed by micropipette spotting. **(b)** Micrograph depicting the chip post-functionalization and after microfluidic gasket integration, showing alignment of the 2 microfluidic channels with the resonator sensors. Scale bars in both micrographs represent 1 mm.

After acquisition of the raw data for all sensors and analysis to track the resonance peak position as described in S2.12.1, the peak position vs. time data for the two reference sensors in each microfluidic channel were averaged. These averaged reference shift data were subtracted from the peak position vs. time data for all sensors in that channel. These reference-subtracted data for the functionalized sensors were subsequently analyzed as described in sections S2.12.3-4, and compared to the raw data.

#### S9.2. Performance comparison for IL-8 detection in simple and complex media

Fig. S11 presents the signal quantification for the sample (IL-8 detection) and amplification (SA-HRP detection) assay stages, for the 6 tested groups described in Table S21. The corresponding quantified average signals and CVs are presented in Table S23. During the IL-8 detection stage (Fig. S11(a)), the measured signal is a combination of the IL-8 binding shift and shift due to nonspecific binding of other proteins and materials in the sample matrix. When we move from a simple buffer matrix to the more complex CCM, the negative control shifts increase from close to zero (-0.0053 ± 0.0441 nm) to 1.0 ± 0.1 nm – approximately 5× the 0.19 ± 0.11 nm shifts that we observe for the IL-8-containing simple buffer sample. As such, we only observe a significant difference between the signal resulting from the IL-8-containing sample (3.125 ng/mL) and the negative control (0 ng/mL) in the case where our sample was the simple PBS-BSA buffer solution and not in the more complex CCM containing additional confounding proteins using either of the tested blocking strategies. Despite reducing the nonspecific binding signal by 5.4 ± 1.2× compared to the BSA blocking strategy, blocking with CCM+BSA yielded negative control signals that remained similar to the IL-8 detection shifts in simple media. Further blocking optimization and/or the use of reference sensors would likely be required to enable direct label-free detection of the intrinsic IL-8 binding shift at this concentration in complex media.

Using the sandwich assay format and quantifying the SA-HRP binding signal (Fig. S11(b)), we are able to resolve significant differences between the IL-8-containing sample and the negative control for all tested matrix and blocking conditions. These results demonstrate the specificity advantage afforded by the sandwich assay, where the signal quantification takes place during a later stage of the assay and is thus less likely to be confounded by nonspecific binding. Our analysis of the variability of this assay is limited by the low number of replicates included in this pilot study. However, even in the simple buffer sample, we observe a higher inter-assay CV for the SA-HRP assay stage of this IL-8 detection assay compared to the direct label-free detection of 1 μg/mL spike protein reported in Fig. 8. The ~320× lower concentration used in this assay likely contributes to this higher variability, as it creates conditions where analyte depletion may be more pronounced and binding shifts are closer to the intrinsic noise levels of the sensors. The more complex assay format may also contribute to this higher variability (with SA-HRP binding shifts dependent not only on the capture antibody-IL-8 binding interaction but also on the detection antibody-IL-8 interaction and the biotin-streptavidin interaction).

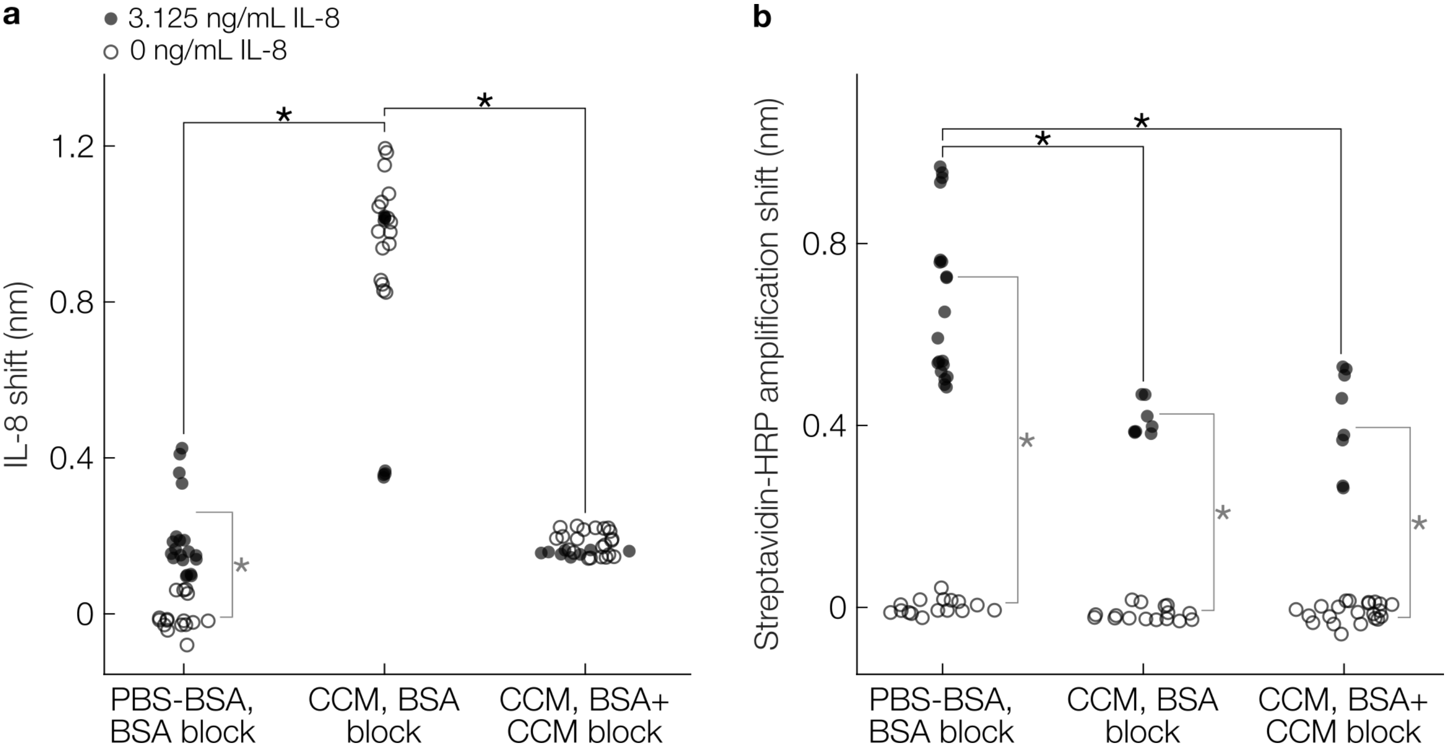

**Fig. S11.** Demonstration of IL-8 detection on PDA-functionalized SiP sensors using a sandwich assay format. Beeswarm plots comparing resonator peak shifts for **(a)** IL-8 detection and **(b)** SA-HRP-based signal amplification assay stages. Data are shown for sensors exposed to samples containing 3.125 ng/mL or 0 ng/mL (neg. control) IL-8, prepared in simple buffer (PBS-BSA) or complex media (complete cell culture medium, CCM). Data are presented for sensors blocked with BSA or BSA+CCM. *$p<0.05$, Mann-Whitney U-tests. Black (horizontal) significance indicators indicate significant differences observed between different assay conditions (buffer type and blocking strategy) for sensors exposed to 3.125 ng/mL Il-8. Vertical (grey) significance indicators indicate significant differences in signal observed between sensors exposed to 3.125 ng/mL IL-8 and the corresponding negative controls (0 ng/mL IL-8) for each assay condition. Further details regarding significance testing can be found in Section S9.3.

Fig. S12 presents the impact of reference subtraction on nonspecific binding signal in a negative control experiment (IL-8 binding assay with a sample consisting of 0 ng/mL IL-8 in CCM). This assay used BSA in PBS blocking and showed high nonspecific binding signal during the sample introduction stage, for both the functionalized and reference sensors. With reference subtraction, we are able to decrease the non-specific binding signal from the sample by an average factor of 13.3 ± 3.6 (from 0.96 ± 0.02 nm to 0.08 ± 0.02 nm). This large reduction in nonspecific signal highlights the utility of reference subtraction, particularly when detecting in complex media with large amounts of background signal. Along with blocking optimization strategies and optimized negative control probe selection as discussed in excellent detail by Bucukovski and Miller (38), the implementation of reference sensors may permit future direct label-free detection of IL-8 without the sandwich assay format.

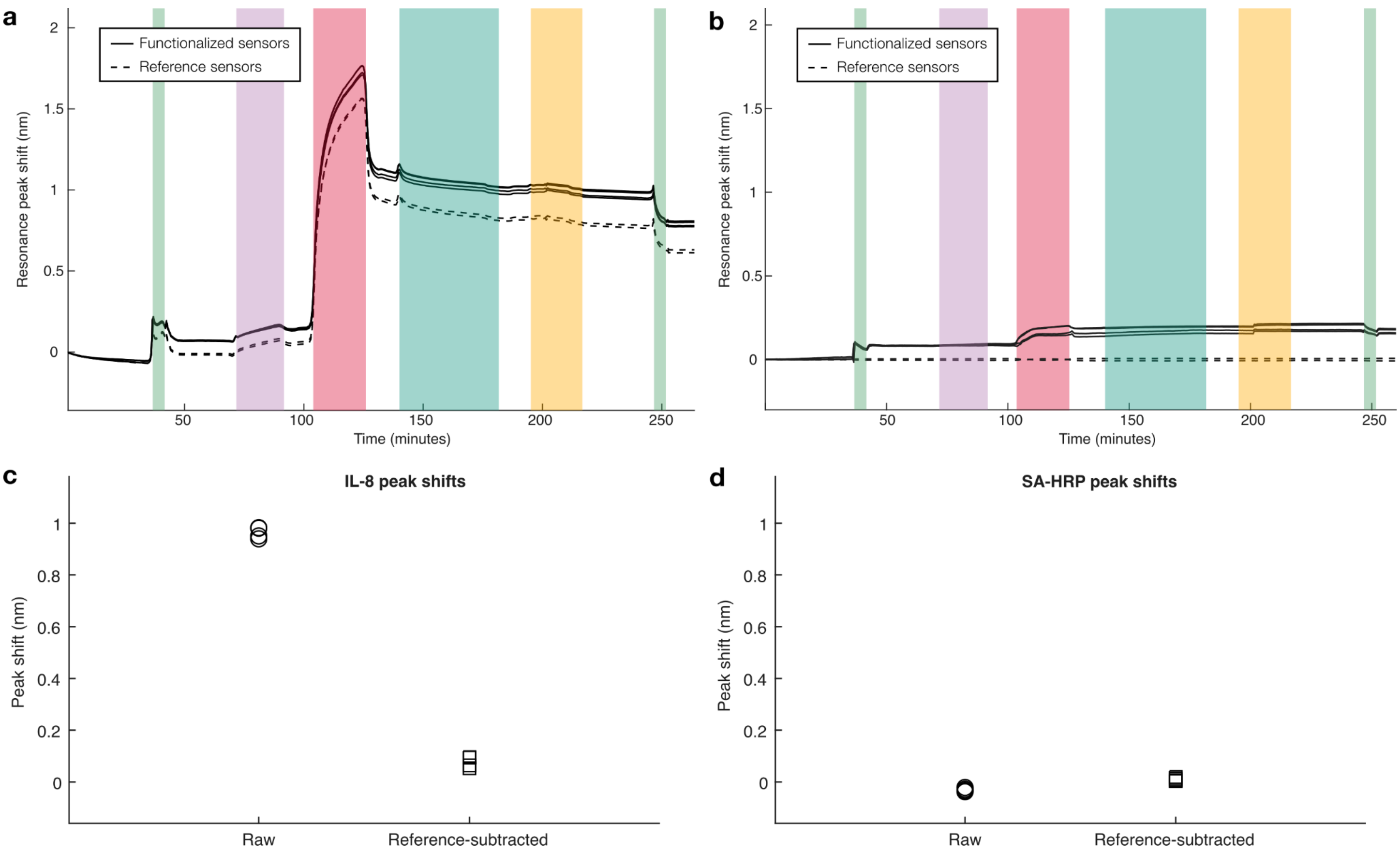

**Fig. S12.** Demonstration of reference-subtraction in a negative control experiment (IL-8 binding assay with a sample consisting of 0 ng/mL IL-8 in CCM) on PDA/spotting-functionalized multiplexed SiP sensors blocked with BSA in PBS. Upstream and downstream sensor groups were functionalized with IL-8 capture antibody, and the middle sensing group was blocked with BSA to serve as a reference as shown in Figure S10. **(a-b)** Comparison of **(a)** raw and **(b)** reference-subtracted sensorgrams for the functionalized and reference sensors, where solid lines indicate signal from the functionalized sensors and dashed lines indicate signal from reference sensors. **(c-d)** Beeswarm plots showing resonance peak shifts measured for the functionalized sensors during **(c)** IL-8 sample exposure and **(d)** SA-HRP signal amplification. Raw signals are represented by the circular markers while reference-subtracted signals are represented by the square markers. Data are shown for n = 4 functionalized resonator replicates and n = 2 reference resonators. These data illustrate that reference-subtraction is able to reduce the nonspecific binding signal from the sample by an average factor of 13.3 ± 3.6 in this pilot assay, potentially complementing optimized blocking strategies to improve detection performance in complex media.

**Table S23.** Quantified IL-8 binding and SA-HRP amplification shifts and their replicability in sandwich immunoassays performed on PDA-functionalized SiP sensors. Data are tabulated for sensors exposed to 3.125 ng/mL and 0 ng/mL IL-8 samples prepared in simple (PBS-BSA) and complex (CCM) buffers. Blocking was performed with BSA or BSA+CCM.

| | | | | | **IL-8 shift** | | | **SA-HRP amplification shift** | | |
|---|---|---|---|---|---|---|---|---|---|---|
| **Running buffer** | **Blocking** | **IL-8 conc. (ng/mL)** | **Number of channel replicates** | **Number of assay replicates** | **Inter-trial mean and std. dev. (nm)** | **Single-ch. intra-assay CVs** | **Inter-assay CV** | **Inter-trial mean and std. dev. (nm)** | **Single-ch. intra-assay CVs** | **Inter-assay CV** |
| **PBS-BSA** | **BSA** | **3.125** | 5 | 5 | 0.19 ± 0.11 | 5.5 ± 3.7% | 57% | 0.67 ± 0.18 | 4.6 ± 5.8% | 27% |
| | | **0** | 4 | 4 | -0.0053 ± 0.0441 | 34 ± 30% | 837% | 0.0016 ± 0.0124 | 96 ± 164% | 791% |
| **CCM** | **BSA** | **3.125** | 2 | 2 | 0.69 ± 0.46 | 1.2 ± 1.0% | 68% | 0.41 ± 0.03 | 5.4 ± 5.2% | 8.3% |
| | | **0** | 4 | 3 | 1.0 ± 0.1 | 2.7 ± 1.3% | 13% | -0.014 ± 0.016 | 25 ± 30% | 116% |
| **CCM** | **BSA+CCM** | **3.125** | 2 | 2 | 0.16 ± 0.00 | 4.1 ± 1.5% | 1.6% | 0.41 ± 0.13 | 13 ± 10% | 32% |
| | | **0** | 6 | 4 | 0.18 ± 0.03 | 4.0 ± 2.9% | 16% | -0.012 ± 0.019 | 50 ± 26% | 161% |

#### S9.3. Statistical testing

Two-tailed Mann-Whitney U tests were performed as described in Section S3 to evaluate the statistical significance of differences in IL-8 detection and SA-HRP amplification shifts between IL-8-containing samples (3.125 ng/mL IL-8) and negative controls (0 ng/mL IL-8), different sample buffers (PBS-BSA vs. CCM), and different blocking strategies (BSA vs. BSA+CCM). These different assay conditions were compared to one-another in a pairwise manner. The resulting p-values for each test are provided in Table S24.

**Table S24.** Statistical analysis of the IL-8 and SA-HRP shift data presented in Fig. S11. P-values reported in bolded text indicate metrics that were found to have a statistically significant difference between the compared assay conditions based on a significance level of $\alpha = 0.05$.

| | | | Conditions compared | | | P-values | |
|---|---|---|---|---|---|---|---|
| | | | **IL-8 concentration (ng/mL)** | **Sample buffer** | **Blocking** | **IL-8 detection shifts** | **SA-HRP amplification shifts** |
| **IL-8 sample vs. negative control** | **PBS-BSA, BSA block** | **Group 1** | 3.125 | PBS-BSA | BSA | **$3.82 \times 10^{-7}$** | **$3.82 \times 10^{-7}$** |
| | | **Group 2** | 0 | PBS-BSA | BSA | | |
| | **CCM, BSA block** | **Group 1** | 3.125 | CCM | BSA | $1.34 \times 10^{-1}$ | **$1.01 \times 10^{-4}$** |
| | | **Group 2** | 0 | CCM | BSA | | |
| | **CCM, BSA+CCM block** | **Group 1** | 3.125 | CCM | BSA+CCM | $7.10 \times 10^{-2}$ | **$4.07 \times 10^{-5}$** |
| | | **Group 2** | 0 | CCM | BSA+CCM | | |
| **PBS-BSA, BSA block vs. CCM, BSA block** | **IL-8 sample** | **Group 1** | 3.125 | PBS-BSA | BSA | **$4.95 \times 10^{-4}$** | **$5.28 \times 10^{-5}$** |
| | | **Group 2** | 3.125 | CCM | BSA | | |
| | **Negative control** | **Group 1** | 0 | PBS-BSA | BSA | **$1.54 \times 10^{-6}$** | **$3.09 \times 10^{-3}$** |
| | | **Group 2** | 0 | CCM | BSA | | |
| **CCM, BSA block vs. CCM, BSA+CCM block** | **IL-8 sample** | **Group 1** | 3.125 | CCM | BSA | **$1.55 \times 10^{-4}$** | $8.78 \times 10^{-1}$ |
| | | **Group 2** | 3.125 | CCM | BSA+CCM | | |
| | **Negative control** | **Group 1** | 0 | CCM | BSA | **$2.12 \times 10^{-7}$** | $4.33 \times 10^{-1}$ |
| | | **Group 2** | 0 | CCM | BSA+CCM | | |
| **PBS-BSA, BSA block vs. CCM, BSA+CCM block** | **IL-8 sample** | **Group 1** | 3.125 | PBS-BSA | BSA | $9.39 \times 10^{-1}$ | **$8.65 \times 10^{-4}$** |
| | | **Group 2** | 3.125 | CCM | BSA+CCM | | |
| | **Negative control** | **Group 1** | 0 | PBS-BSA | BSA | **$2.12 \times 10^{-7}$** | $9.48 \times 10^{-2}$ |
| | | **Group 2** | 0 | CCM | BSA+CCM | | |
